# Congenital Zika virus infection impairs corpus callosum development

**DOI:** 10.1101/2021.11.11.468315

**Authors:** Raissa R. Christoff, Jefferson H. Quintanilha, Raiane O. Ferreira, Jessica C. C. G. Ferreira, Daniel M. Guimarães, Bruna Valério-Gomes, Luiza M. Higa, Átila D. Rossi, Janaina M. Vasconcelos, João L.S.G. Vianez, Maria Bellio, Amilcar Tanuri, Roberto Lent, Patricia P. Garcez

## Abstract

Congenital Zika Syndrome (CZS) is a set of birth defects caused by Zika virus (ZIKV) infection during pregnancy. Microcephaly is its main feature, but other brain abnormalities are found in CZS patients, such as ventriculomegaly, brain calcifications, and dysgenesis of the corpus callosum. Many studies have focused on microcephaly, but it remains unknown how ZIKV infection leads to callosal malformation. To tackle this issue, we infected mouse embryos *in utero* with a Brazilian ZIKV isolate and found that they are born with a reduction in callosal area and density of callosal neurons. ZIKV infection also causes a density reduction of PH3+ cells, intermediate progenitor cells and SATB2+ neurons. Moreover, axonal tracing revealed that callosal axons are reduced and misrouted. Also, ZIKV infected cultures show a reduction of callosal axon length. GFAP labelling showed that *in utero* infection compromises glial cells responsible for midline axon guidance. The RNA-Seq data from infected brains identified downregulation of axon guidance and axonogenesis related genes. In sum, we showed that ZIKV infection impairs critical steps of corpus callosum formation by disrupting not only neurogenesis but also axon guidance and growth across the midline.

**Summary Statement:** Zika virus infection during development impairs the formation of corpus callosum by disturbing axon guidance and growth of callosal neurons.

## Introduction

Zika virus (ZIKV) has gained worldwide attention since its devastating impact on brain development in humans was identified (Mlakar *et al*., 2016). The virus infects neural progenitor cells *in vitro* and *in vivo*, impairing cell cycle, causing cell death, and leading to microcephaly (Cugola *et al*., 2016; Garcez *et al*., 2016; Tang *et al*., 2016). Although microcephaly is the main feature of ZIKV infection, other brain abnormalities have been identified (De Oliveira Melo *et al*., 2016; Chimelli *et al*., 2017). The set of signs present in affected newborns whose mothers were infected during pregnancy are now known as Congenital Zika Syndrome (CZS). Besides microcephaly, CZS patients also display ventriculomegaly, brain calcifications, and corpus callosum dysgenesis (De Fatima Vasco Aragao *et al*., 2016; de Souza *et al*., 2018).

The corpus callosum is the main telencephalic commissure of placental mammals connecting the two cerebral hemispheres and contributing to the proper functioning of different sensorimotor and cognitive abilities (Tomasch, 1954). Corpus callosum dysgenesis is a brain malformation characterized by absence or size reduction of the commissure, with abnormal formation of ectopic bundles (Tovar-Moll *et al*., 2007; 2014; Edwards *et al*., 2014). In typical neurodevelopment, callosal projection neurons arise from neural progenitor cells and migrate to their final position on the cortical plate where axons are extended to a contralateral target following guidance molecules including those secreted by midline glial cells (Gobius and Richards, 2011). In mice, the midline glial populations are generated from radial glial progenitors between embryonic days E13 to E17 and express the astroglial marker GFAP. These structures are known as glial wedge, *indusium griseum* glia; and midline zipper glia, positioned close or at the boundary between hemispheres (Gobius and Richards, 2011). Defects of midline structures compromise axon guidance of callosal neurons, resulting in corpus callosum malformations (Shu *et al*., 2003).

During corticogenesis, ZIKV early infection disrupts radial glial progenitor cells by altering their proliferation, as shown by the reduction of proliferation markers such as Ki67 and PH3. Also, viral infection reduces intermediate progenitors that are responsible for cerebral cortex expansion (Cugola *et al*., 2016; Garcez *et al*., 2018). This scenario leads to a reduced cortical thickness by the loss of neuronal cells of different cortical layers. In later infection, ZIKV targets progenitor cells that will give rise to the oligodendrocytes, causing impairment in proliferation and differentiation (Li *et al*., 2018). Primarily, ZIKV impairs proliferation, but it can also infect different types of neural cells such as astrocytes, postmitotic neurons, and microglia (Lum *et al*., 2017; van den Pol *et al*., 2017; Limonta *et al*., 2018; Ledur *et al*., 2020). Since ZIKV is capable of infecting different types of glial cells and progenitors, both at early and late stages of brain development, it is conceivable that ZIKV infection indirectly disturbs corpus callosum development.

Although many studies have shown the impact of ZIKV in neurogenesis and its link to microcephaly, it is unknown how ZIKV interferes with corpus callosum development. Therefore, the aim of this study was to identify the cellular mechanisms that could lead to corpus callosum dysgenesis in ZIKV infection in a *in utero* animal model. Here, we provide evidence that ZIKV infection during corticogenesis causes corpus callosum developmental abnormalities by not only reducing the callosal neurons production but also impairing axon guidance and outgrowth.

## Results

### Congenital ZIKV infection leads to a reduction in the corpus callosum in mice

To investigate the cellular mechanisms underpinning corpus callosum developmental abnormalities after ZIKV infection, embryonic mice were inoculated with 10^4^ PFU of Brazilian ZIKV (ZIKV) or culture medium (MOCK) at E15 *in utero* via an intracerebroventricular approach and harvested through postnatal day 4 (P4), when they were euthanized and had their brains removed (Fig. 1A-B). To characterize this infection model, we measured the brain weight at P4 and found that the infected brains were lighter than controls (Fig. 1C). Also, when measuring only cerebral cortex weight, the effect was evident (Fig. 1D). Using ZIKV NS1 immunostaining, ZIKV protein was detected in the cerebral cortex at P4 (Fig. 1E,1F). Also, plaque assay confirmed the presence of infectious viral particles in cortical samples (Fig. 1G). To analyse the corpus callosum formation, we compared sagittal brain sections of infected animals to the controls and found that the corpus callosum area is severely reduced in infected animals (Fig.1H-J). Corpus callosum dysgenesis could be caused by multiple developmental defects. To investigate whether the *in utero* infection impacted the number of callosal neurons, we used the isotropic fractionator method to estimate the total amount of cortical SATB2+ cells (Supplementary Fig. 1). At P4, we observed lower counts of both the total nuclei (DAPI+) and SATB2+ cells, when compared to controls (Supplementary Fig.1C-D). Together, these results show that 9 days after infection (dpi) animals present abnormal corpus callosum development, with a significant reduction in SATB2+ callosal cells, which is consistent with the callosal dysgenesis found in some patients with Congenital Zika Syndrome (De Fatima Vasco Aragao et al., 2016).

**Figure 1.**
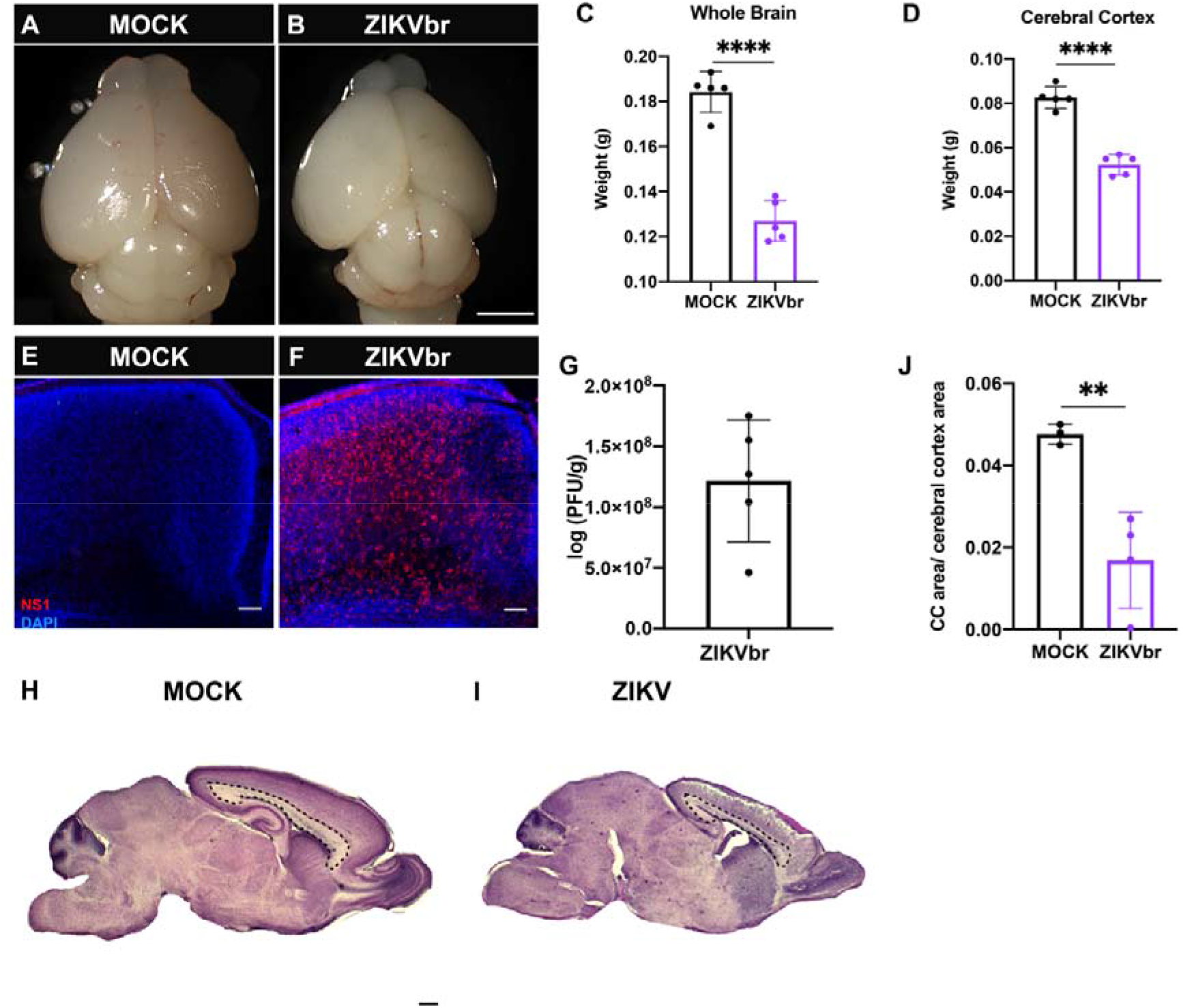
Zika virus congenital infection at E15 causes microcephaly and corpus callosum dysgenesis at P4. (A, B) Dorsal views of MOCK animals and ZIKV-infected mice. Scale bar = 1mm. (C, D) Quantitative analysis of brain and forebrain weights (g) of ZIKV and MOCK groups. N= MOCK (5) ZIKV (5). (E, F) Immunohistochemical labelling for NS1 (red) and DAPI staining (blue) on MOCK and ZIKV-infected cerebral cortex. Scale bars = 50μm (G) Quantitative analysis of infected particles in ZIKV animals by plaque assay. N= ZIKV (5). (H, I) Sagittal sections of MOCK and ZIKV-infected brains, stained with cresyl-violet. The corpus callosum area is delimited by dashed lines. Scale bar = 100μm. (J) Quantification of corpus callosum area normalized to cerebral cortex area in quasi-sagittal brain sections of ZIKV and MOCK animals. N= MOCK (3) ZIKV (3). Student’s t-test* = p < 0.05; ** = p < 0.005; **** = p < 0.00005.

### Congenital ZIKV infection causes cell death at the cortical plate

ZIKV infection is known to cause cell death, contributing to the microcephalic phenotype (Tang et al., 2016, Garcez et al., 2016, Cugola et., 2016). To verify if in our model cell death could account for the SATB2+ callosal cell reduction, we stained the cerebral cortex with cleaved caspase-3 (CASP3+) apoptotic marker at different time points. Interestingly, after 5dpi (50 PFU, low titre infection), there were no CASP3+ cells in the cerebral cortex (Fig. 2A-C). Cell death was only noticed after 7dpi (Fig. 2D-F), and, more pronouncedly, at 10dpi (Fig. 2G-I). In addition, animals infected with ZIKV presented an increased number of pycnotic nuclei at P4 (Supplementary Fig.1E), confirming an increased cell death after 10dpi. These results suggest that loss of cortical cells at P4 (Supplementary Fig. 1C-D) could be due to cell death induced by ZIKV infection.

**Figure 2.**
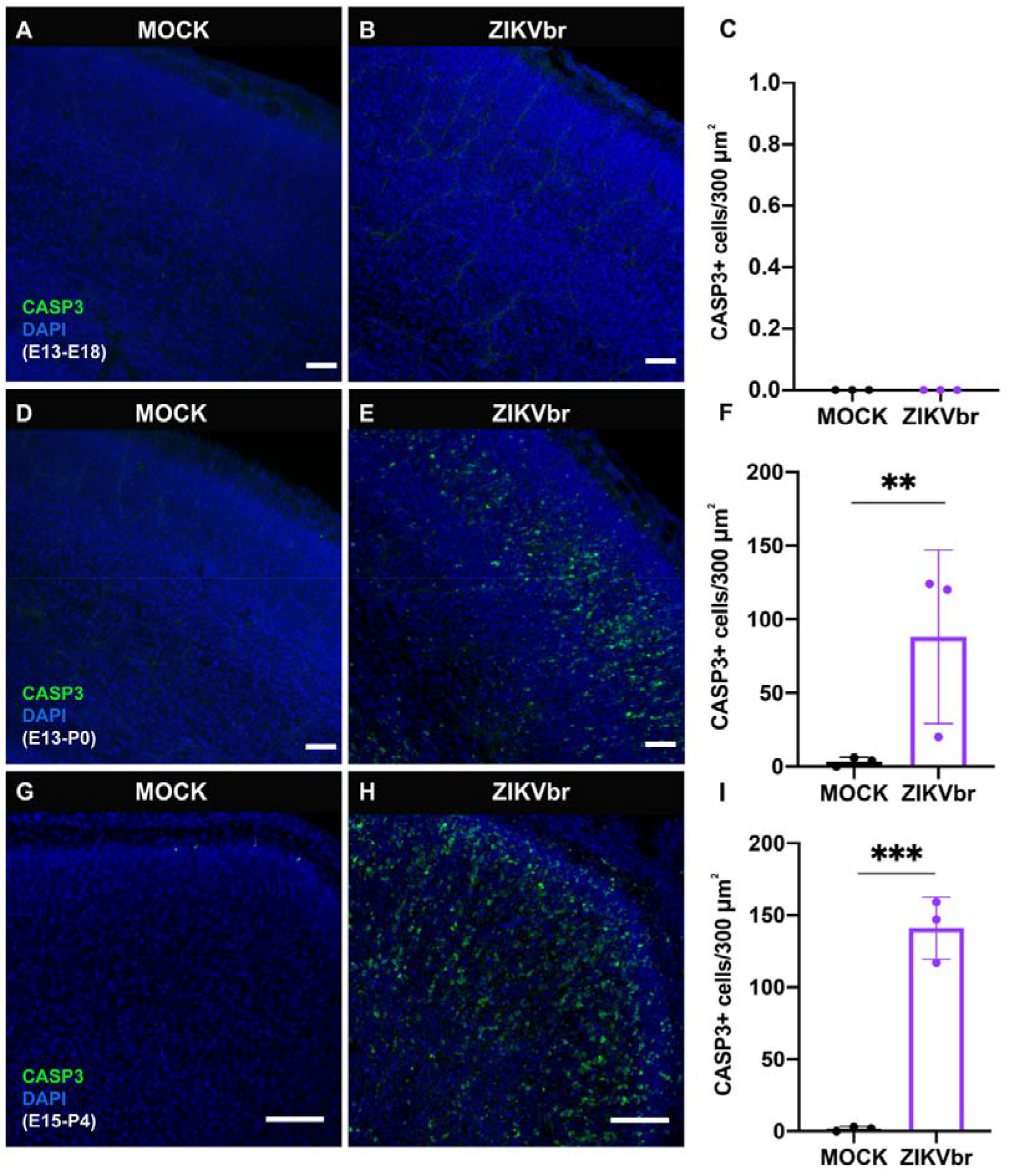
Zika virus congenital injection results in cell death only after 7 days post-infection. (A, B) Immunohistochemistry for activated Caspase 3 (CASP3) on coronal sections of MOCK compared to ZIKV-infected E13 embryos, harvested at E18. DAPI staining in blue. (C) Quantification of CASP3+ cell density in 300 μm^2^ at E18. N= MOCK (3) ZIKV (3). (D, E) Immunohistochemistry for CASP3 (green) on coronal sections of MOCK compared to ZIKV-infected E13 embryos, harvested at P0. Scale bars = 50 μm (F) Quantification of CASP3+ cell density in 300 μm^2^ at P0. N= MOCK (3) ZIKV (3). (G, H) Immunocytochemistry for CASP3 (green) on coronal sections of MOCK compared to ZIKV-infected E15 embryos, harvested at P4. Scale bars = 50μm. (I) Quantification of of CASP3+ cell density in 300 μm^2^ at P4. N= MOCK (3) ZIKV (3). Data presented here as the mean ± SEM from at least six sections prepared from three animals obtained from two or three litters for each condition in this analysis and subsequent quantifications, unless stated otherwise. Student’s t-test ** = p < 0.005; *** = p < 0.0005.

### ZIKV infection leads to defect in callosal neurons production

In addition to cell death, ZIKV is also known to impair neural progenitor’s cell cycle (Tang et al., 2016). In the low titre ZIKV infection model, cell death was only observed after birth (Fig. 2D-F). Therefore, we investigated whether ZIKV embryonic infection would impair callosal neurons production by impacting the proliferation of neural progenitors. At E15, cortical layer V neurons migrate towards the cortical plate and callosal II/III neurons are generated. After 3dpi (E18), although infected brains show SATB2+ neurons positioned similarly to controls (Fig. 3A-C), their density is significantly reduced (Fig. 3D). The cortical plate was divided into three sectors of equal areas encompassing the whole cortical thickness, and SATB2+ neurons were quantified in relation to DAPI+ nuclei. To quantify the progenitor density, we immunostained sections with PH3 G2/M marker. We observed that ZIKV infection reduces the density of progenitors at the ventricular zone after 3dpi (Fig 3E-G). After 5dpi, the intermediate progenitor density (TBR2+) is also reduced (Fig. 3H-J). Taken together, these results show that ZIKV infection impairs proliferation and reduces intermediate progenitor cells, responsible for generating upper granular layers, where the majority of callosal cells are found. Also, ZIKV infection reduces the callosal neurons density within the cortical plate, but not their location (Fig. 3D).

**Figure 3.**
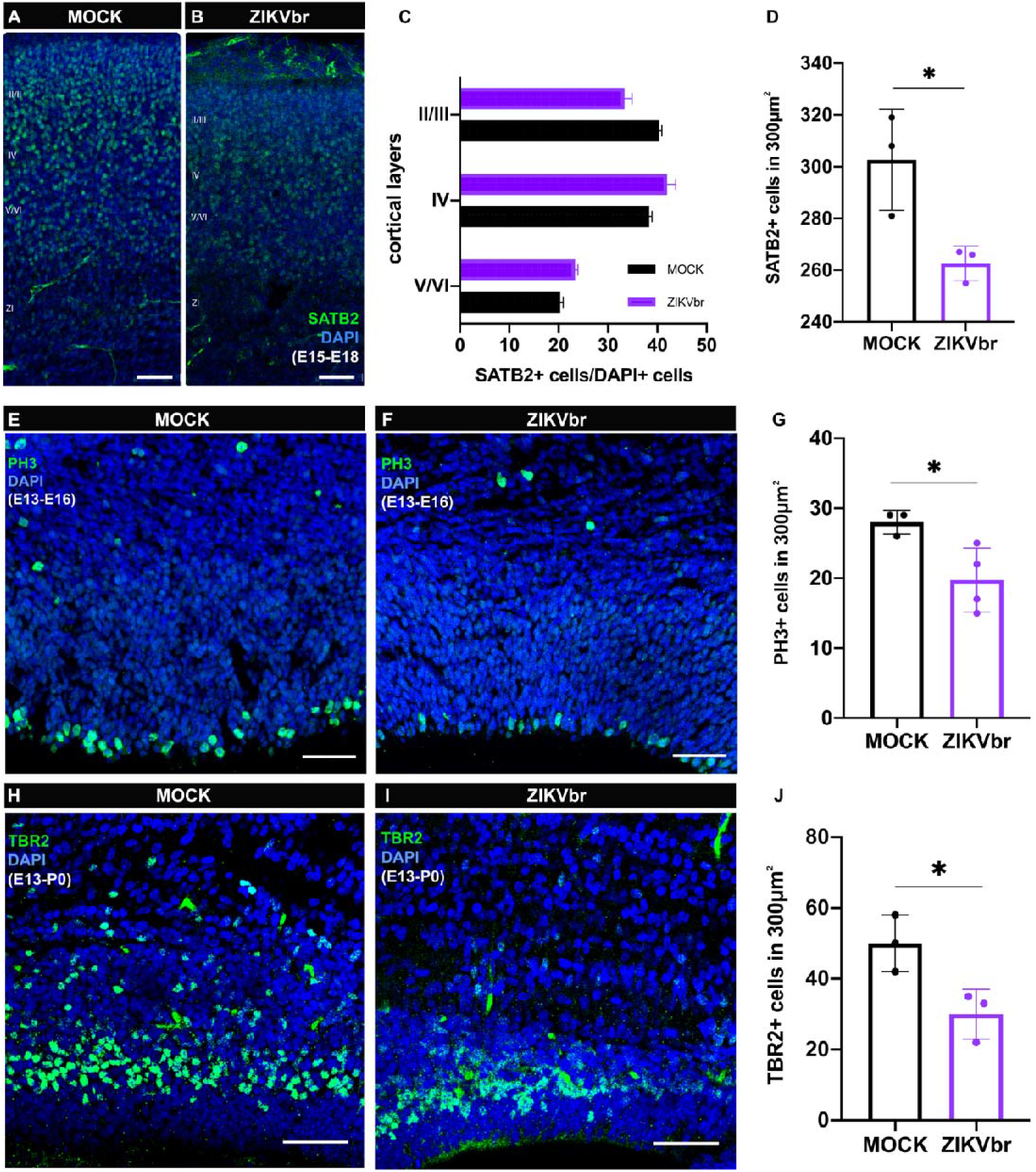
Zika virus *in utero* injection leads to intermediate progenitors and callosal neuron density reduction. (A, B) Immunohistochemistry for callosal nuclear marker SATB2 (green) counterstained with DAPI (blue) in cortical coronal sections of MOCK and ZIKV mice injected *in utero* at E15 and harvested at E18. (C) Quantifications of SATB2+/DAPI+ cells in the different cortical layers (I to VI). N= MOCK (3) ZIKV (3). (D) Quantifications of SATB2+ cell density in 300 μm^2^. N= MOCK (3) ZIKV (3). (E, F) Immunohistochemistry for phospho-histone 3 (PH3) of MOCK compared to ZIKV-infected E13 embryos, harvested at E16 in 300 μm^2^. (G) Quantification PH3+ cell density in 300 μm^2^. N= MOCK (3) ZIKV (3). (H, I) Immunohistochemistry for intermediate progenitors’ marker TBR2 (green) in cortical sections of MOCK and ZIKV-infected E13 embryos, harvested in P0. (J) Quantification of TBR2+ cell density in 300 μm^2^. N= MOCK (3) ZIKV (3). Scale bars = 50μm. Student’s t-test * = p < 0.05.

### ZIKV infection impairs axonal growth of callosal neurons

Axon growth is a critical step in corpus callosum development (Gobius & Richards, 2011). To establish whether ZIKV infection directly impairs the axon extension of callosal neurons, we used E14 cortical neurons primary culture and infected with ZIKV compared to controls. After 4 days *in vitro* (div), no difference was observed in the number of SATB2+ neurons or the presence of cell death (data not shown). However, we quantified the length of callosal axons using ImageJ tracing tools and found that they were ∼25% shorter compared to mock exposed axons (Fig. 4A-C). Also, Sholl analysis revealed that dendrites of these ZIKV exposed neurons are less arborized compared to controls, suggesting that ZIKV infection impairs neurite growth (Fig. 4D-F). To verify whether ZIKV affects callosal axon growth *in vivo*, a DiI crystal, a lipophilic dye that diffuses within fixated membranes, was inserted and incubated into the dorsolateral cerebral cortex surface to allow anterograde axonal tracing (Fig. 5A-F). After 20 days of DiI diffusion, the E18 brain was sectioned, and axons could be analysed. Interestingly, infected animals show not only a reduction of the callosal bundle at the midline but also signs of axonal misrouting (Fig. 5J-K). To accurately measure the width of the callosal axons, we stained axons with L1CAM and analyzed the corresponding rostrocaudal levels at 5dpi. Infected animals display a reduction of the midline bundle width compared to controls (Fig. 5G-I).

**Figure 4.**
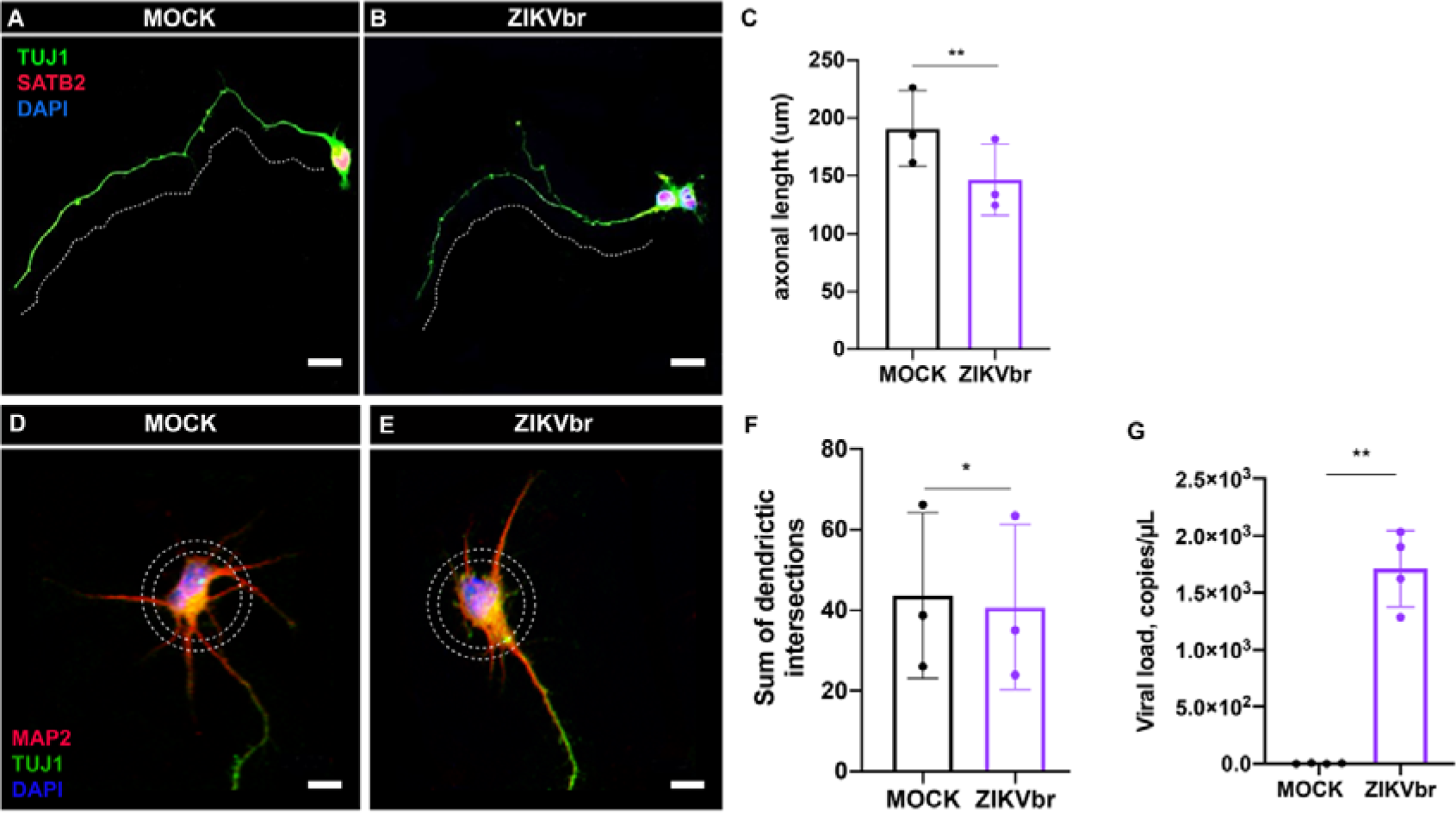
Zika virus *in vitro* infection reduces axon extension and dendritic arborization of callosal neurons. (A, B) Primary dorsal cortical progenitors exposed to ZIKV or MOCK after 24 hours and cultivated for 3 days post infection (3dpi). Co-immunolabelling for TUJ1 (green) and SATB2 (red) to measure the longest neurite (dashed lines). Scale bars = 10 μm (C) Quantification of axonal length in ZIKV and MOCK primary neuron cultures. N= 3 independent experiments. (D, E) Co-immunolabelling for MAP2 (red) and TAU1 (green). Sholl analysis was performed using concentric dashed circles to access arborization complexity in MOCK or ZIKV. Scale bars = 10μm (F) Quantification of the number of dendritic intersections with the concentric circles in ZIKV and MOCK primary neuron cultures. Blue = DAPI. N= 3 independent experiments. (G) ZIKV viral RNA was determined by real-time PCR after 3dpi from 4 independent experiments. Student’s t-test * = p < 0.05; ** = p < 0.005.

**Figure 5.**
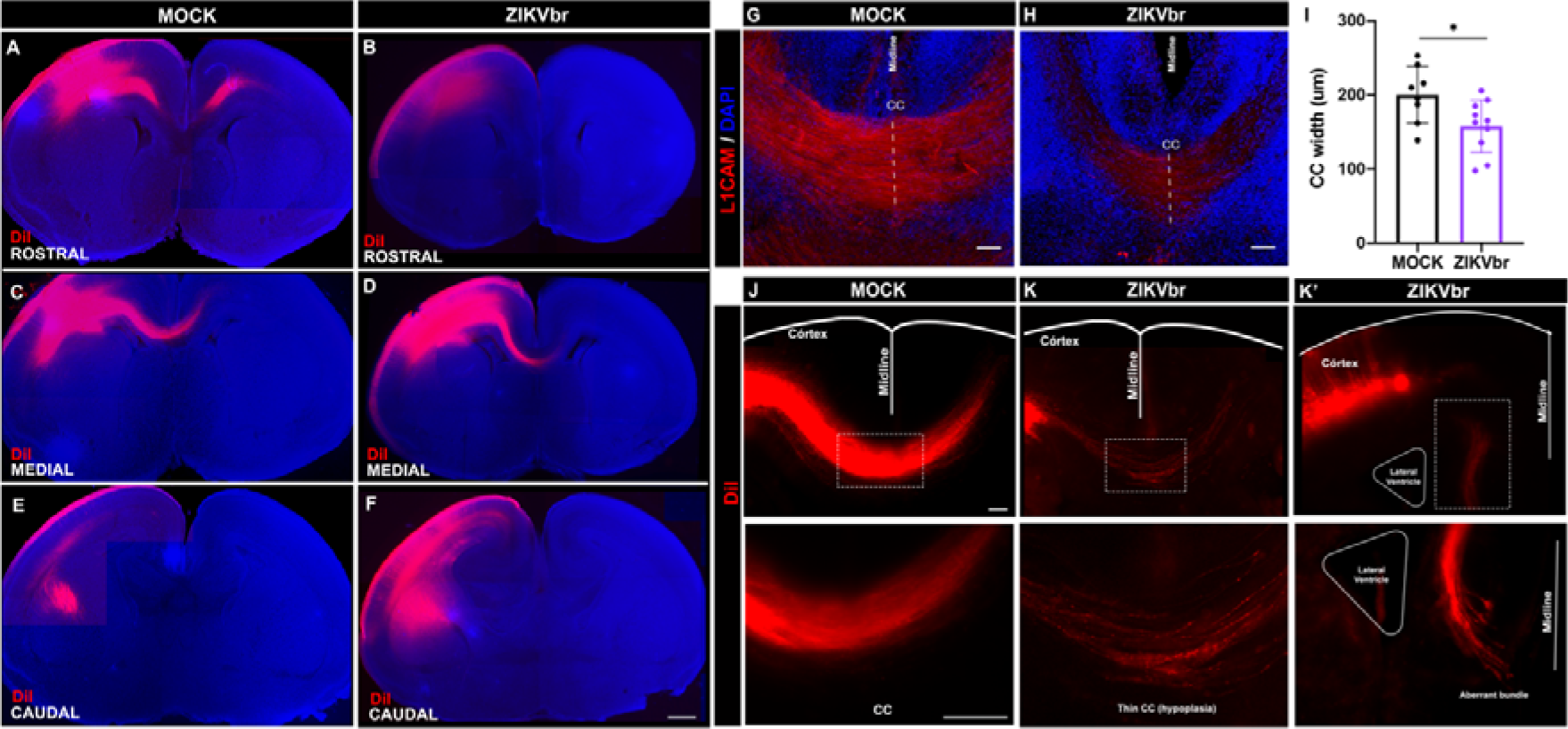
Zika virus congenital infection impairs midline crossing of callosal axons at E18 *in vivo*. (A-F) Rostro-caudal levels of DiI anterograde tracer (red) in E18 brains of MOCK and ZIKV infected animals at E13. Scale bar = 100μm. (G, H) L1CAM labeling of callosal axons at midline region of MOCK or ZIKV infected animals at E13 and harvested at E18. Scale bars = 50μm (I) Width measurement of callosal axons labeling for L1CAM (red) at midline region. Blue = DAPI. N= MOCK (8) ZIKV (10). Student’s t-test * = p < 0.05. (J, K) Brain coronal section of MOCK animal with DiI labelling fibers compared to the ZIKV-infected animal with defasciculated fibers at midline. Scale bars = 50μm (K’) Brain coronal section of ZIKV-infected animal with DiI labelling fibers showing misrouted axon fibers that fail to cross the midline. Scale bar = 50μm. CC = corpus callosum

### ZIKV infection impairs midline glia cells

Our *in vitro* results show that ZIKV infection generates callosal neurons with shorter axons. We also observed that *in vivo*, axons that are crossing the midline through corpus callosum are reduced and misrouted. These findings suggest that axon guidance could be altered. Midline glia consists of transient cell populations that hold the role of guiding axons across the midline during development. We measured the signal intensity of GFAP+, a marker of glial cells (Fig. 6A-D). In ZIKV-infected mice, GFAP+ intensity is low, suggesting that this cell population is compromised, which may explain the defasciculated axons we observed.

**Figure 6.**
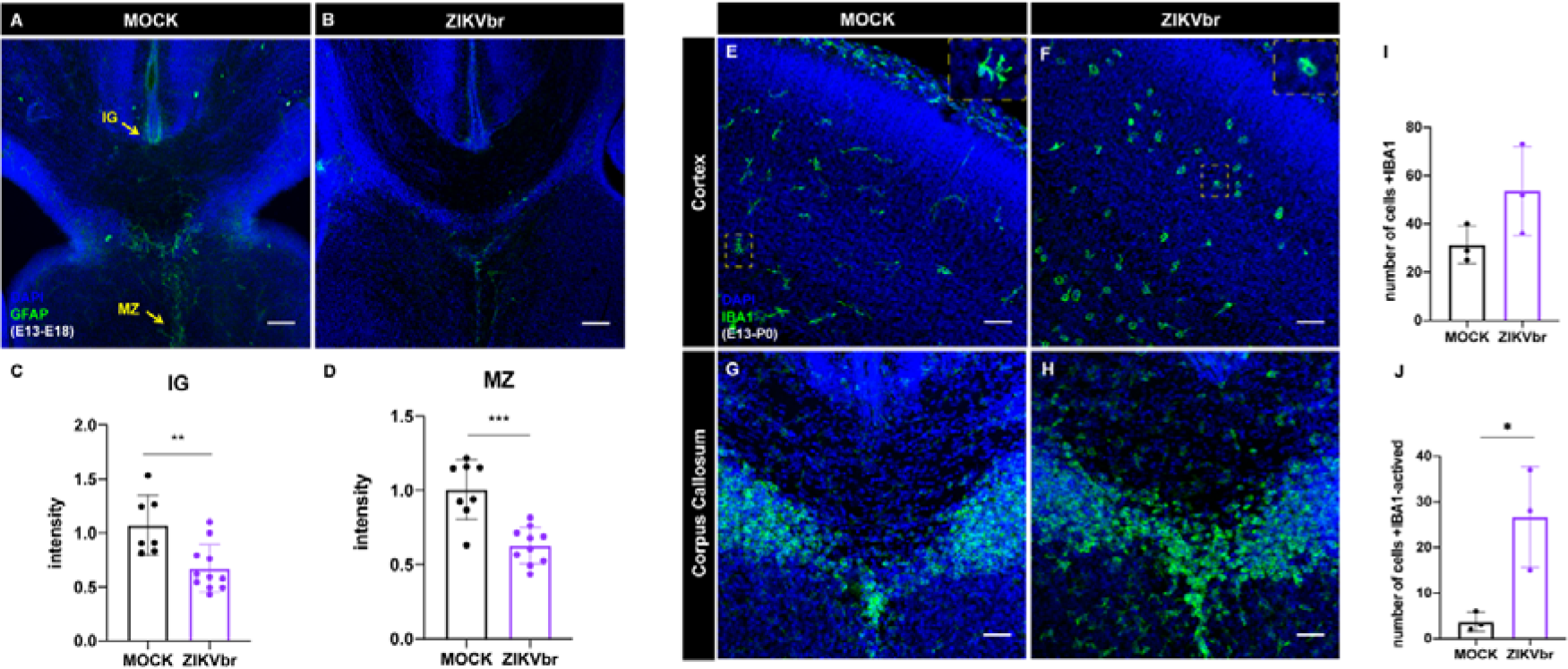
Zika virus infection at E13 alters midline glial populations and leads to microglial activation. (A, B) Immunocytochemistry for GFAP (green) at the midline regions of embryonic brain slices of MOCK (A) and ZIKV (B) animals. (C) Quantification of GFAP signal intensity at the *indusium griseum glia* (IG) close to the midline of ZIKV and MOCK. N= MOCK (8) ZIKV (10). (D) Quantification of analysis of GFAP signal intensity at the *midline zipper glia* (MZ) region of ZIKV and MOCK. N= MOCK (8) ZIKV (10). (E-H) Immunocytochemistry for microglial marker IBA1 (green) at the cortical plate of embryonic brain slices of MOCK (E) and ZIKV (F) and midline regions of MOCK (G) an ZIKV (H) infected *in utero* at E13 and analyzed at P0. (I) Quantification of IBA1+ density at cortical plate in 450 μm^2^. N= MOCK (3) ZIKV (3). (J) Quantification of IBA1+ density at cortical plate with amoeboid morphology (active microglia) in 450 μm^2^. N= MOCK (3) ZIKV (3). Scale bars = 50μm. Blue = DAPI. IG = *indusium griseum glia*; MZ = *midline zipper glia*. Student’s t-test * = p < 0.05; ** = p < 0.005; *** = p < 0.0005.

### ZIKV infection deregulates gene expression of embryonic brain

To investigate the overall unbiased effect of ZIKV infection on gene expression and cortical development, animals’ brains infected at E15 (10^4^PFU) were analyzed by RNA-Seq. After 3dpi, ZIKV-infected animals already show a range of differentially expressed genes (DEGs), with 1,414 significantly up-regulated and 1,040 down-regulated [false discovery rate (FDR) < 0.05]. A list of differentially expressed genes is available in Table S1. In total, the DEGs are enriched in more than 800 terms in Gene Ontology analysis (GO). DEGs upregulated are mainly overrepresented in viral infection and immune response, while down-regulated genes are associated with neural development, metabolic processes, and synaptic formation (Supplementary Fig. 2). Circos plot analyses indicate a set of downregulated hub genes that are important for axonal formation, axonal projection, and axonal guidance (Fig. 7A). For example, genes of the *β*-tubulin family (e.g., *tubb3* and *tubb2b*) that are crucial for cytoskeleton assembly, therefore important for axonal growth, appear altered. In addition, *actb* is a protein-coding gene that encodes for *β*-actin, a key compound of the axon growth cone. In contrast, circos plot of upregulated genes reveal genes that modulate neuronal death and microglial activation (Fig 7B). In fact, microglia staining for IBA1+ presents an activated morphology at cortical region after 7 days of *in utero* infection (Fig. 6E-J)

**Figure 7.**
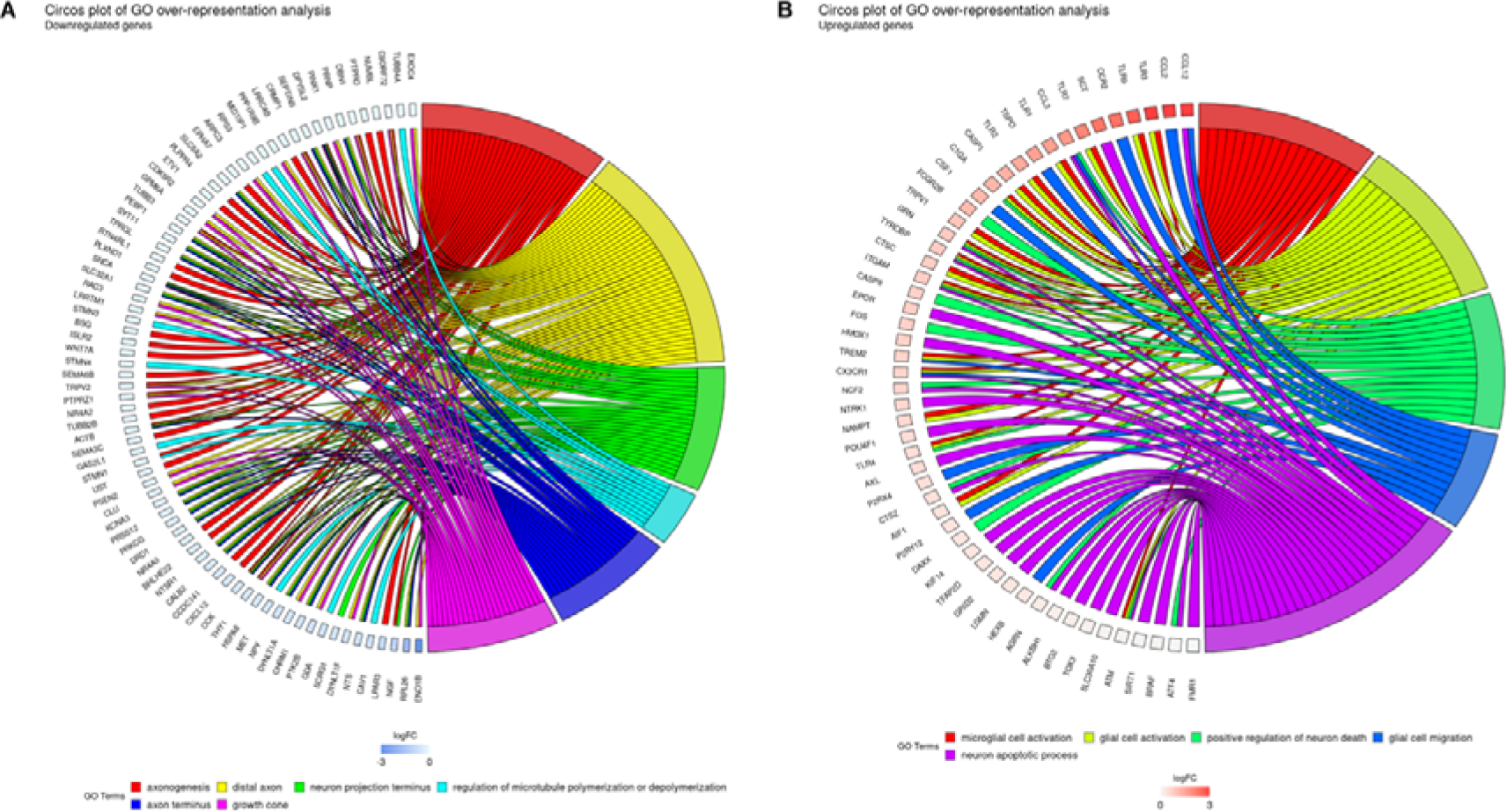
Zika virus congenital infection at E15 deregulates gene expression in embryonic brains after 3dpi. (A, B) Circos plot illustrating the hub genes in ZIKV-infected brain when compared to MOCK arranged in order of their expression level sharing relevant cellular processes. (A) Down-regulated genes. (B) Up-regulated genes. N= MOCK (5) ZIKV (3).

## Discussion

Congenital Zika Syndrome is a set of birth defects caused by ZIKV infection during pregnancy that has microcephaly as its hallmark. ZIKV studies focus mainly on the viral impact on neurogenesis and its relation to microcephaly. Yet, other brain abnormalities are present in CZS, such as corpus callosum dysgenesis. In this study, we identified defects in proliferation and axonal extension of callosal cells, as well as alterations in midline glial cells, which can contribute to our understanding of the cellular mechanisms by which ZIKV infection leads to malformation of the corpus callosum.

We used *in utero* infection to assess the brain abnormalities caused by ZIKV. Like other *in vivo* studies that used this approach, ZIKV was capable of replicating in the mouse cerebral cortex and causing microcephaly (Fig. 1A-C) (Li *et al*., 2016, 2018; Zhang *et al*., 2019). Besides microcephaly, we found that the corpus callosum size was significantly reduced in infected animals (Fig.1H-J). This result is consistent with reported cases of CSZ in humans (De Fatima Vasco Aragao *et al*., 2016). In addition, we found an important reduction in the total number of cells in the cerebral cortex which agrees with reduced cortical thickness found in other studies of embryonic ZIKV infection (Li *et al*., 2016, 2018; Shao *et al*., 2016; Wu *et al*., 2016). Moreover, SATB2+ cells in particular, which are known to represent cortical callosal projection neurons, were severely reduced in number (Supplementary Fig.1B-C). The significant change in the number of callosal neurons explains, at least partially, the reduced size of the corpus callosum in infected animals.

ZIKV infection is associated with massive cell death ratios on the cortical plate at 10dpi (Fig 2C-F). This effect could account for the loss of callosal neurons at P4 (Supplementary Fig.1B-C). As previously shown, extensive neuronal death is detected in different mouse models of infection (Li *et al*., 2016; Shi *et al*., 2018; Zhang *et al*., 2019). In these models, cell death appears to be concentrated in the cortical plate and takes place both for immature and mature neurons. In our study, we also observed the same phenomenon.

ZIKV infection impairs neurogenesis by disrupting cell cycle and reducing proliferation (Garcez *et al*., 2016; Tang *et al*., 2016). In this study, we observed that apoptosis typically occurred in the cortical plate only after birth, therefore we asked if ZIKV infection disrupts callosal neurons during the embryonic period by disturbing neural progenitor cell proliferation. At E15, callosal neurons that occupy cortical layers II/III are originated from progenitors located at the ventricular surface of the cerebral cortex (Shen *et al*., 2006). We observed that infecting animals at this age reduces the number of SATB2+ neurons at E18 (Fig. 3A-B), a timepoint at which cell death was not noticed (Fig. 2A). In agreement with previous studies (Li *et al*., 2016; Shi *et al*., 2018; Zhang *et al*., 2019), we also found that ZIKV infection reduces proliferation at the ventricular zone, which leads to less intermediate progenitors (TBR2+) (Fig. 3D-G). In later stages of corticogenesis, TBR2+ progenitors are responsible for producing most glutamatergic projection neurons in all cortical layers (Hevner, 2019). Therefore, in our model of infection, ZIKV reduces the number of callosal neurons by targeting proliferation and intermediate progenitors’ cells without causing cell death at the embryonic stage.

Microcephaly is mainly caused by a reduction in the number of neurons due to impaired cell proliferation (Doobin, Dantas and Vallee, 2017). Although a reduction of the callosal commissure may be directly caused by this key feature of microcephaly, in mice, we found that corpus callosum malformation is better explained by the disruption of developmental steps after neurogenesis, such as defective midline patterning, axonal misrouting and growth (Shu *et al*., 2003). To evaluate if ZIKV directly impairs axonal extension of callosal neurons, we infected primary cell cultures of embryonic mice at E14. As observed at 3dpi, callosal neurons exhibit shorter axons compared to controls (Fig. 4A-C). Also, dendritic branching was compromised (Fig. 4D-F). Our results agree with those of Goodfellow *et al*. (2018), who infected cultured mature neurons with ZIKV and found that neurite length and branching were reduced in size. Here, we show that embryonic SATB2+ callosal neurons have a shorter axon and less arborized dendrites. In addition, using an axonal tracer to analyze callosal tracts *in vivo,* we found axonal misrouting at the midline (Fig. 5K). These results indicate that ZIKV infection compromises an important step of corpus callosum formation.

Axonal misrouting could be explained by the lack of axonal guidance signaling, which is provided by midline glial cells (Shu *et al*., 2003). In mouse models of ZIKV infection, glial cells are also disrupted. Li and collaborators (2018) noticed that mice infected at E15 and evaluated at P5 showed a reduced number of oligodendrocytes in the corpus callosum. Here, we observed that during development infected animals showed a reduction of GFAP+ midline astrocytes (Fig. 6A-D), suggesting that these cells populations are impaired, and therefore indicating that the virus is capable of infecting different cell populations in the brain.

An important limitation of our work is to clearly show that the observed effects were caused directly by the viral infection and were not derived from inflammatory responses. ZIKV infection produces a strong inflammatory response and during embryonic development, inflammatory molecules are known to contribute to neurodevelopmental disorders (Jiang *et al*., 2018) In agreement with other studies that investigate the effect of ZIKV infection on gene expression, our results demonstrated an upregulation of genes that are involved in viral infection, immune and inflammatory response (Li *et al*., 2016; Shao *et al*., 2016; Chang *et al*., 2021). Despite the strong presence of inflammatory factors, ZIKV infection also deregulates key genes important for neurodevelopment (Barbeito-Andrés *et al*., 2020; Chang *et al*., 2021), which was also observed in our work (Fig.7). Semaphorins and Ephrins are a family of proteins that act on axonal guidance by attracting or repelling growing axons at the midline (Hu *et al*., 2003; Mire *et al*., 2018). Interestingly, our transcriptome analysis showed that genes that belong to these families are downregulated (*sema3c, sema6b, epha7*). Together, the reduction of GFAP+ marker of midline glia and the lower expression levels of cue molecules may generate a poor signaling environment for crossing axons at the midline. Moreover, *tubb3* is downregulated and has been described in the literature as an important gene to the formation of commissural structures (Tischfield *et al*., 2010). Also, our data showed that *actb*, a key protein-coding gene is deregulated. *β*-actin is a cytoskeleton protein that structurally composes the axon growth cone, which is responsible for identifying the molecular guidance cues. For example, Netrin1, a well-established guidance molecule, is known to modulate the levels of *β*-actin local expression at the growth cone (Leung *et al*., 2006). Another gene that was downregulated in our analysis is *calb2* that encodes for a calcium-binding protein calretinin (CR). A transient population of neurons that express CR is necessary to guide axons through the midline region together with the midline glia (Niquille *et al*., 2009). Our results demonstrated that in addition to the inflammatory response, genes important for axon guidance and outgrowth are deregulated. Our GO enrichment analysis pointed that the genes involved in synapse formation and axon terminal structure were downregulated in infected brains. Similar processes involving axon and dendrite development were found to be deregulated in previous work (Chang *et al*., 2021).

As the formation of corpus callosum comprises complex steps of development, this is the first study to suggest a mechanism by which ZIKV infection interferes with the formation of corpus callosum during embryonic development. These findings contribute to our understanding of the pathophysiology of ZIKV infection. CSZ patients exhibit a range of brain abnormalities that are not always associated with microcephaly (Nielsen-Saines *et al*., 2019). The ability of ZIKV to disturb critical steps of corpus callosum development reveals the importance of monitoring pregnant women who have been exposed to ZIKV and doing a follow-up of their newborns since corpus callosum continues to develop postnatally (Moldrich *et al*., 2010).

Further studies should address whether the changes in axonal extension and axon guidance can be explained by the lack of key players that are crucial for cortical development, either produced by the midline glia, which are altered in our work, or by other cells that also play this role.

## Materials and Methods

### Animals and ZIKV infection

Pregnant Swiss mice were housed in the Animal Care Facility of the Microbiology Institute of the Federal University of Rio de Janeiro and maintained in standard animal housing with food and water *ad libitum* and circadian cycles of 12 hours light/12 dark. All procedures were approved by the Committee on Ethics of Animal Use for Research of the Federal University of Rio de Janeiro, protocol 040/19. Pregnant dams were anesthetized, and the uterus exposed to handle the embryos carefully. A volume of 1.5 μl of a Brazilian ZIKV isolate (Recife/Brazil, ZIKV PE/243, accession no: KX197192.1) or MOCK was injected into one side of the lateral ventricle of the embryonic mouse brain. On E15, mice were injected with 10^4^ PFU of ZIKV brain tissues were harvested either E18 and P4. On E13, mice were injected with 30 PFU of ZIKV and harvested either E16, E18, and P0.

### Virus detection

Viral RNA was extracted from primary neuronal culture supernatant after four days *in vitro* (4DIV) with 3 days post infection (3dpi), with the RNeasy Plus Mini Kit (QIAGEN), following the recommendations of the manufacturer. To determine the viral load of these samples, reverse transcription of viral RNA followed by quantitative PCR was performed with GoTaq^®^ Probe 1-Step RT-qPCR System (Promega) on a 7500 Real-Time PCR System (Applied Biosystems), using primers and probe described by Lanciotti *et al*. (Lanciotti *et al*, 2008). ZIKV RNA copies were calculated by interpolation onto a standard curve composed of eight 10-fold serial dilutions of a synthetic ZIKV RNA based on the region targeted by the set of primers and probe. The ZIKV quantification was expressed as ZIKV RNA copies per gram of tissue.

Quantification of infectious particles was performed by plaque assay. Brain tissue was mechanically homogenised with ultra turrax T10 (ika) in 1 ml of DMEM supplemented with 2% fetal bovine serum and 2x penicillin/streptomycin. Brain tissue homogenates were centrifuged at 3400 x g for 5 minutes. The clarified supernatants were serially diluted and incubated with confluent monolayers of vero cells. Virus titers were determined by plaque assay performed on Vero cells. Virus stocks or samples were serially diluted and adsorbed to confluent monolayers. After 1 h, the inoculum was removed, and cells were overlaid with semisolid medium composed of alpha-MEM (GIBCO) containing 1.25% carboxymethyl cellulose (Sigma Aldrich) and 1% FBS (GIBCO). Cells were further incubated for 5 days when cells were fixed in 4% formaldehyde. Cells were stained with 1% crystal violet in 20% ethanol for plaque visualization. Titers were expressed as plaque-forming units (PFU) per gram.

### Primary neuronal culture and infection

Pregnant dams were euthanized by cervical dislocation after anesthesia. E14 brains were harvested and dissected, followed by mechanical dissociation of the cerebral cortex with a micropipette into a cell suspension. The dissociation of cortical tissue was done in Neurobasal in a final volume of 1ml. 100,000 cells were added per plate with Neurobasal culture medium supplemented with 2% B27 and antibiotics, then kept in the incubator under 5% CO_2_ at 37°C. After 1DIV, cortical neurons were exposed to ZIKV (MOI 1) or MOCK. After 1 hour, the original culture medium was returned to the culture and the neurons were incubated for further 3 days (3dpi). With 4DIV, the culture was fixed with 4% paraformaldehyde for further immunohistochemical analysis.

### Immunohistochemistry and imaging

Embryonic brains were fixed with 4% paraformaldehyde overnight, cut coronally at 70 µm with a vibratome (VT1000S, Leica, Germany), and stored in 0.01% PBS. After standard antigenic retrieval, coronal sections were permeabilized with 0.1% Triton X-100 (Sigma-Aldrich, USA) and incubated with 3% bovine serum albumin (Sigma-Aldrich, USA) for 2h. Next, the following primary antibodies were incubated overnight: mouse anti-SATB21:200 (GenWay 20-372-60065); rabbit anti-PH3 1:500 (Millipore 06-570); mouse anti-GFAP 1:200 (Sigma G3893); rabbit anti-TBR2 1:500 (Abcam AB23345); rabbit anti-Caspase3 1:300 (Cell signaling 9664); anti-IBA1 1:1000 (Wako 019-19741) and mouse anti-NS1 1:10. Subsequently, samples were washed with PBS and incubated with secondary antibodies: goat anti-rabbit Alexa Fluor 488 1:500 AP132JA4 (Millipore, Etobicoke, CAN) and goat anti-mouse Alexa 546 1:500 AP192SA6 (ThermoFisher Scientific, USA). Nuclei were stained with DAPI (0.5 mg/ml) for 20 min. Images were acquired with a TCS SP8 confocal microscope (Leica, Germany) with an oil immersion 20x objective of high numerical apertures (NA). Analyses of the acquired images were carried out using Fiji software.

### Immunocytochemistry and quantitative analyses

Three different experiments were carried out in duplicates for further analysis. Immunocytochemistry was carried out similarly after vibratome brain sections. The following primary antibodies were used: rabbit SATB2 1:200 (Abcam AB34735); mouse Tuj1 1:100 (Millipore MAB1637). For evaluation of the axonal length, approximately 30 cells per well were equally analyzed under the experimental conditions. To establish a comparison criterion, the largest axons of positive DAPI/SATB2/TUJ1 neurons were chosen to be analysed in the two experimental groups. Axonal length was measured on ImageJ, outlining the extension of the largest neurite from the cell body to the growth cone. To quantify the number of cells, images of five different fields were taken at random (20x magnitude) in each well. The number of DAPI+ and DAPI+/SATB2+ cells was counted in each photo and the percentage of DAPI+/SATB2+ cells in relation to DAPI+ is extracted.

### Histology

Nissl staining (cresyl-violet) was used to quantify the corpus callosum area of animals. After fixation, brains were split at the midline and sectioned parasagittally in the vibratome (150 μm). The sections were then stained with cresyl-violet, (Sigma-Aldrich) for 8 minutes, then dehydrated with ethanol at different concentrations (75%, 95%, and 100%) for 5 minutes each. After dehydration, the sections were immersed in 100% Xylol (Sigma-Aldrich) for 10 minutes, and the slides were cover slipped with Entellan.

### Dye labelling

DiI (1,1’-dioctadecyl-3,3,3’,3-tetramethyl-indocarbocyanine perchlorate, Invitrogen) is a lipophilic dye used as an axonal tracer for fixated tissue.After tissue collection and fixation, a DiI crystal was inserted into the pial surface of the dorsolateral parietal cortex of E18 embryos and incubated for 2 weeks in PBS for diffusion from the cell body to the fibers. After incubation, the brains were serially sectioned in a vibratome (150 μm), at the coronal plane, and analyzed under a fluorescence Zeiss Axioimager Microscope.

### Isotropic Fractionator for total and neuronal cell counting

To assess the total number of cells, the isotropic fractionator technique was performed as described by Valério-Gomes et al., 2018. Briefly, brains of P4 mouse pups were collected and fixed in 4% PFA by immersion for two weeks. Then, the cerebral cortex was dissected, weighed, and mechanically dissociated for 15 min using a glass tissue homogenizer (Pyrex^TM^) with buffer-detergent solution (40 ml sodium citrate + 1% Triton X-100). A suspension of nuclei was transferred to 15ml falcon tubes and then prepared by adding 0.1M of phosphate buffer saline (PBS) until reaching a 5ml of final volume. From this homogeneous suspension, a 1mlaliquotwas taken and 2% of DAPI (20mg/L) was added for total cell quantification in the Neubauer chamber using a Axioimager fluorescence microscope (Zeiss). The total number of cells was estimated by multiplying the mean nuclear density by the total suspension volume and 15.625 (Neubauer factor number for 16 squares counted). Pyknotic nuclei were counted in parallel with total DAPI nuclei by observations of the presence of small and degenerated DAPI+ nuclei. From this same suspension, other aliquots of 1ml were taken for immunocytochemistry. To identify and count callosal neurons, immunocytochemistry was performed using the nuclear SATB2 antibody. For that, aliquots were centrifugated and washed trice in 0.1M PBS (1,500RPM, 5 min each) and then incubated with 1:200 mouse anti-SATB21:200 (GenWay 20-372-60065) in blocking solution (2μl BSA 5%, 30μl NGS and PBS to a complete volume of 200) for up to 48h in agitation at 4-8°C. After three centrifugations, aliquots were incubated with 1:500 Alexa 546 anti-mouse in the same blocking solution in agitation at room temperature for 2h. To estimate the SATB2 absolute number, the percentage obtained by counting SATB2+ positive nuclei in 500 DAPI+ nuclei was multiplied by the total number of cells.

### Total RNA isolation, library construction, and sequencing

A total of 8 frozen brain samples (five controls and three infected with ZIKV) were processed in TissueLyser® (Qiagen) for 30 seconds at 30Hz for cell disruption, using Stainless Steel Grinding Balls. Total RNA isolation was performed with an inhouse protocol based on TRIzol® Reagent (Thermo Fisher Scientific), Chloroform and isopropyl alcohol. Total RNA quantitation was performed with Qubit RNA HS Assay kit (Thermo Fisher Scientific, USA) and RNA quality was assayed on an Agilent 2100 Bioanalyzer using the Agilent RNA 6000 Pico Kit (Agilent Technologies, USA). Libraries were built using TruSeq Stranded Total RNA Library Prep Kits (Illumina, USA) according to the manufacturer’s instructions. To analyse final libraries fragment size distribution and quantify them, we used Agilent 2100 BioAnalyzer and High Sensitivity DNA Kit (Agilent Technologies, USA), and a qPCR-based KAPA library quantification kit (KAPA Biosystems, USA). Finally, libraries were pooled using a final concentration of 20 pM and subjected to paired-end sequencing using Illumina HiSeq 2500 (Illumina, USA) platform and HiSeq Paired-End Cluster Kits v4.

### RNA-Seq analysis

The quality of the raw sequenced Illumina reads was assessed using FastQC 0.11.9 (Andrews *et al*., 2012). Quality and adapter trimming was performed with Trim Galore! 0.6.6 (Krueger, 2015) using a default phred score of 20. After that, the trimmed reads were mapped to the GRCm39 (M27) mouse genome reference assembly (Frankish *et al*., 2019) using STAR 2.7.9a (Dobin *et al*., 2013) and counted with HTseq (Anders, Pyl and Huber, 2014). The edgeR (Robinson, McCarthy and Smyth, 2009) package was used to normalize the data with the TMM method and to test for differentially expressed genes using the exactTest function (p-value < 0.05). Then, the differentially expressed genes were used to perform GO over-representation analysis (Benjamini Hochberg adjusted p-value<0.05) using clusterProfiler v3.18.1 [7] in the R statistical environment [8]. The relationships between enriched GO terms and genes were visualized using circos plots generated by the R package GO plot 1.0.2 [9].

## Statistical Analysis

Statistical analysis was performed using GraphPad Prism 8. *In vivo* experimental data were analysed using unpaired Student’s t-test. For *in vitro* experiments statistical analysis was considered as paired. Data are expressed as mean ± S.D. For each independent experiment at least 3 animals were used per group. *p < 0.05, **p < 0.005 and *** p < 0.0005; **** p < 0.00005.

## Acknowledgements

We thank Dr Danielle Rayee for her intellectual and experimental insights on this work. We thank Dr. Joice Stipurky and Professor Flavia Gomes for sharing their lab facility. We thank Adiel Batista, Camila Lopes, Fabio Jorge Moreira da Silva, Grasiela Matioszek and Lena Dalva Rubim for their valuable technical assistance. We thank Publicase for the intensive scientific writing workshop.

## Author contribution

**R.R.C.** concept/design, data acquisition, analysis and interpretation, J.H.Q, R.O.F, J.C.C.G.F., D.M.G., B.V-G, data acquisition and analysis; **L.M.H** and **A.D.R.**, provided the viral titration and viral qPCR, **J.M.V** and J.L.S.G. performed the RNA-Seq experiments and bioinformatics, **M.B., A.T., R.L** provided supervision and lab facility; **P.P.G** supervised, conceived and designed the experiments. All authors contributed to the manuscript writing and critical review.

---

**Supplementary Figure 1.**
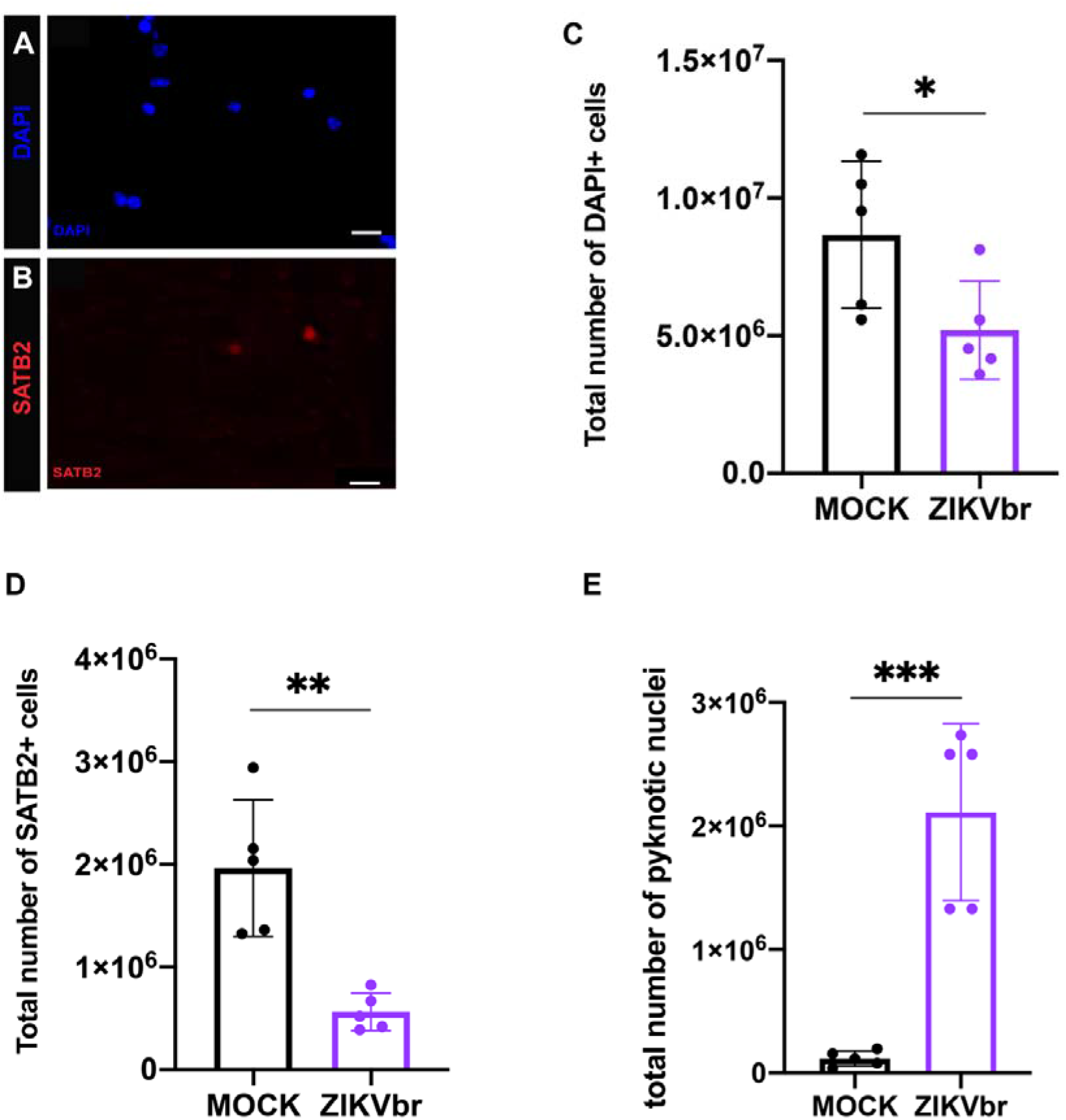
Zika virus congenital infection at E15 causes reduces the total number of callosal neurons and increases pycnotic nuclei at P4. (A, B) Cortical brain nuclei labelling with (A) DAPI (blue) and (B) SATB2 (red) after Isotropic fractionator technique. Scale bars = 10 μm (C) Quantitative analysis for cortical absolute number of cells (DAPI staining in blue) in infected and control tissue. N= MOCK (5) ZIKV (5). (D) Total number of callosal neurons - SATB2 staining (red) on cell nuclei isolated with the isotropic fractionator technique in ZIKV and MOCK animals. N= MOCK (5) ZIKV (5). (E) Quantification of pyknotic nuclei using the isotropic fractionator technique. N= MOCK (5) ZIKV (5). Student’s t-test ** = p < 0.005; *** = p < 0.0005.

**Supplementary Figure 2.**
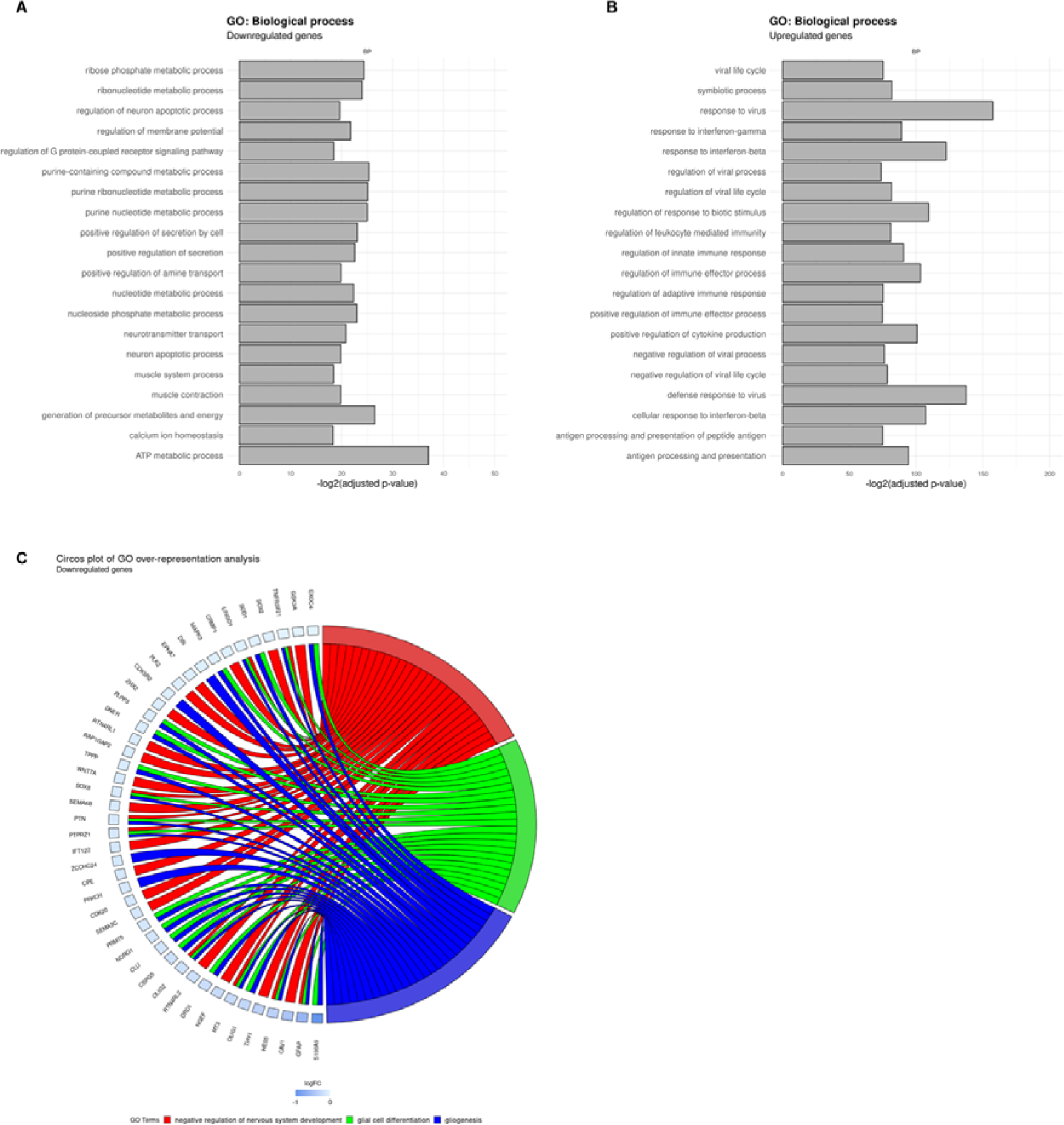
Zika virus congenital infection at E15 deregulates gene expression in embryonic brains after 3dpi. (A, B) RNA-seq analyses of brains of ZIKV infected-animals compared to MOCK at E15 and harvest at E18. Enrichment analysis of GO terms for the 20 most down-regulated genes (A) and up-regulated genes (B). (C). Circos plot illustrating the hub genes overrepresented in ZIKV-infected brain when compared to MOCK sharing GO biological process related to Nervous System arranged in order of their expression level. N= MOCK (5) ZIKV (3).

**Supplementary Table 1.**
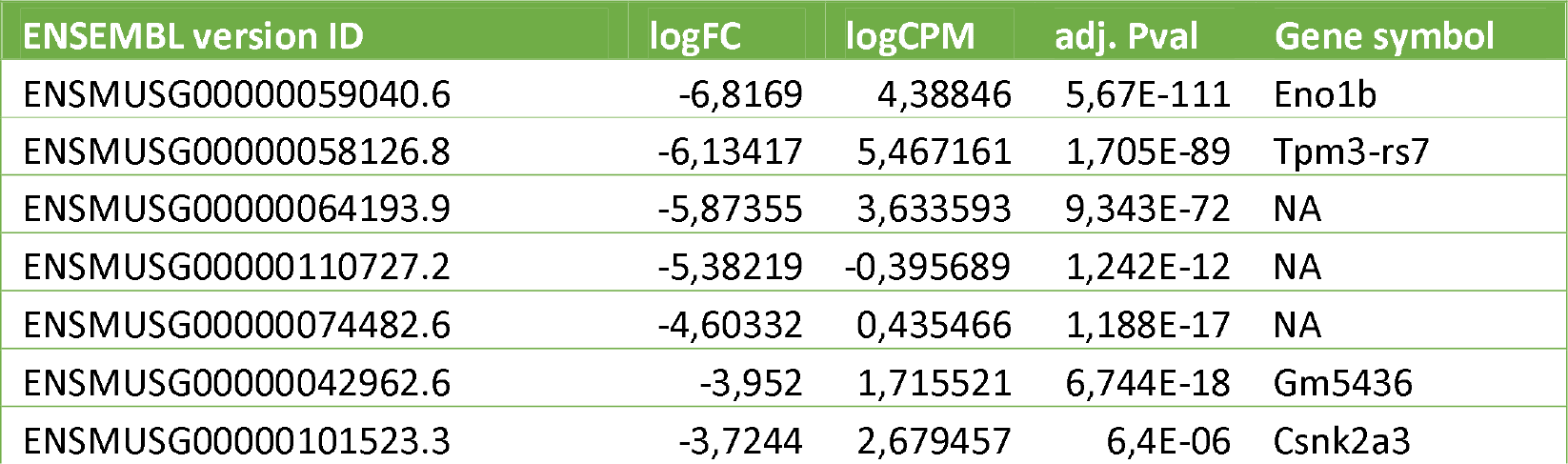
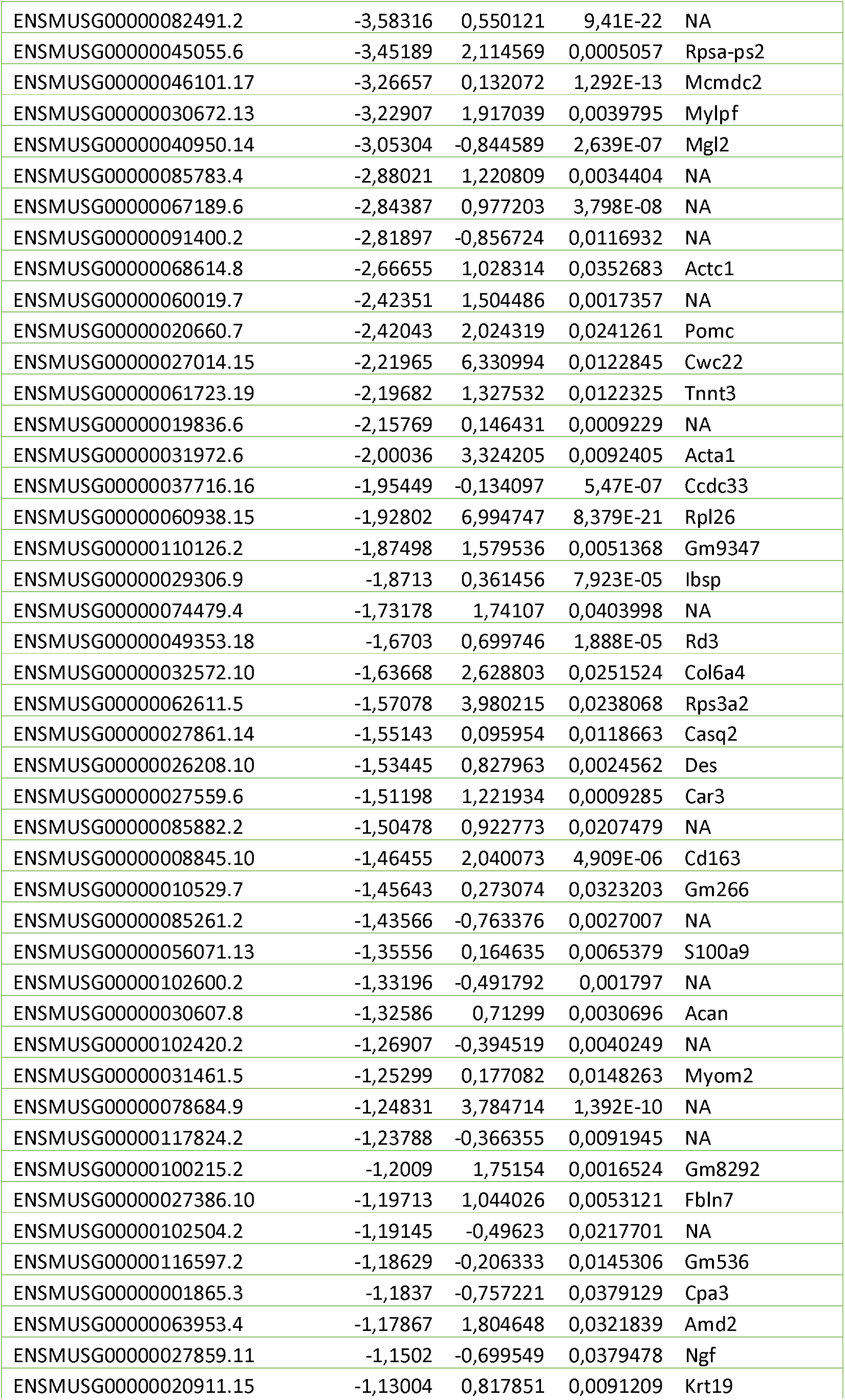
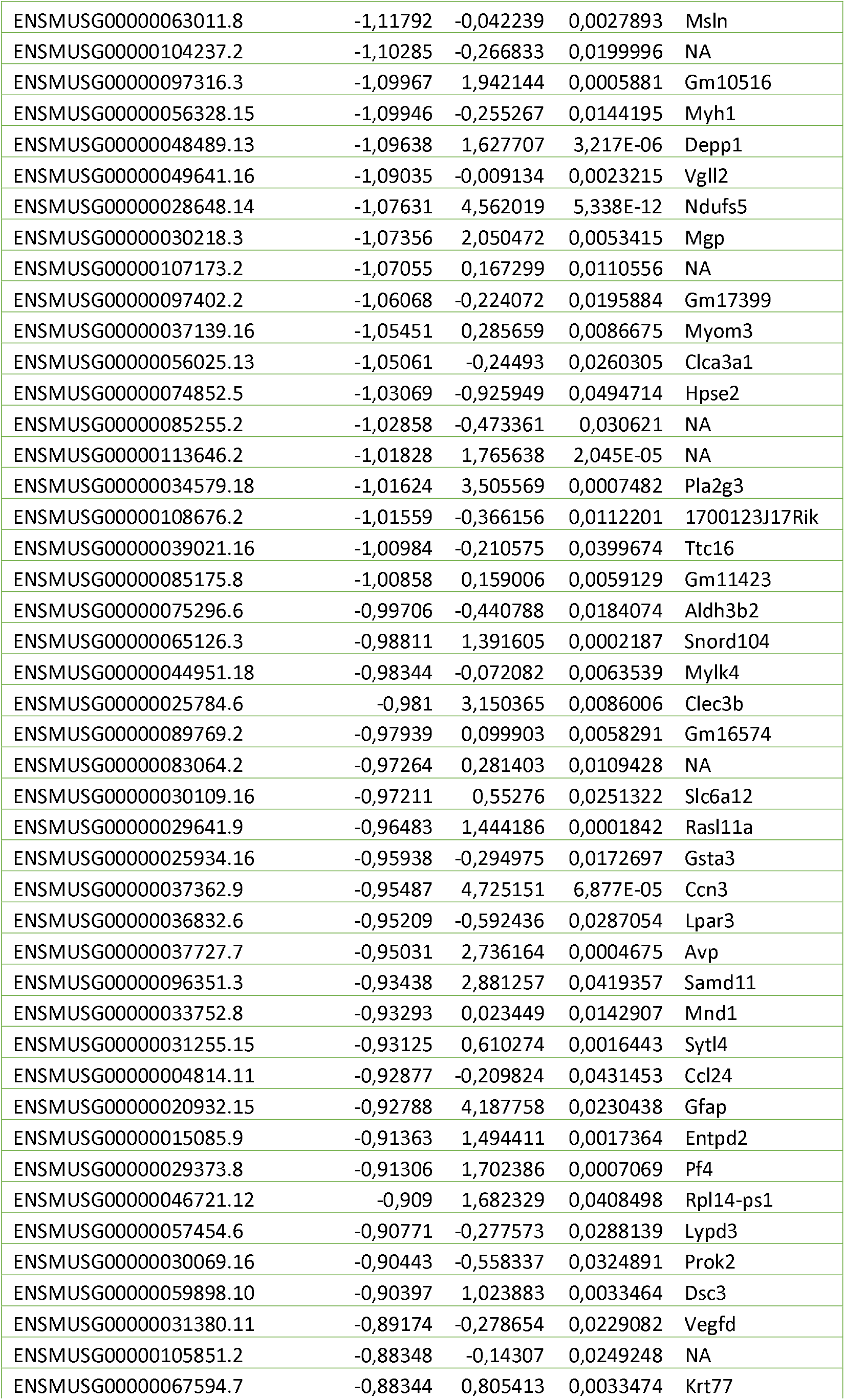
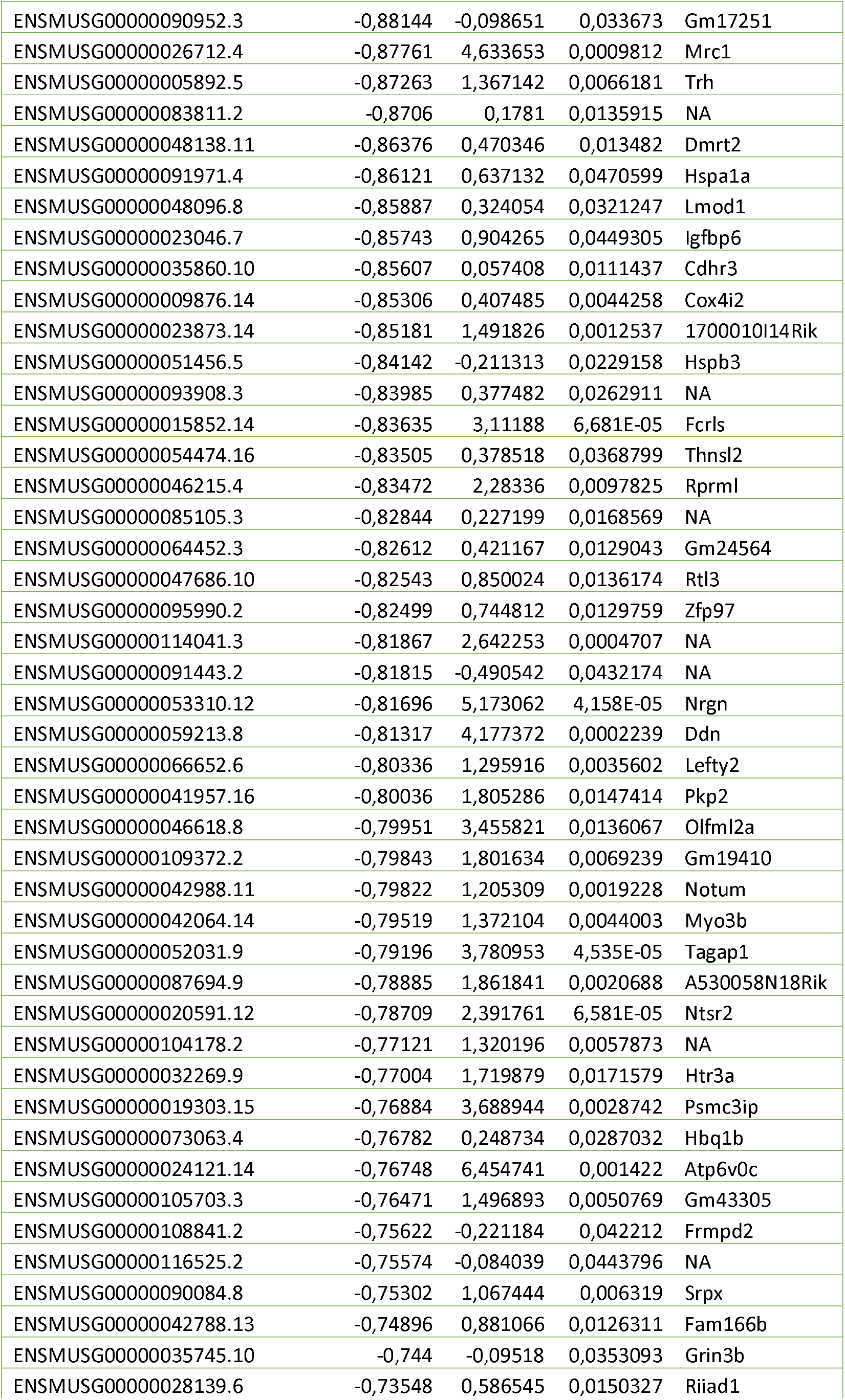
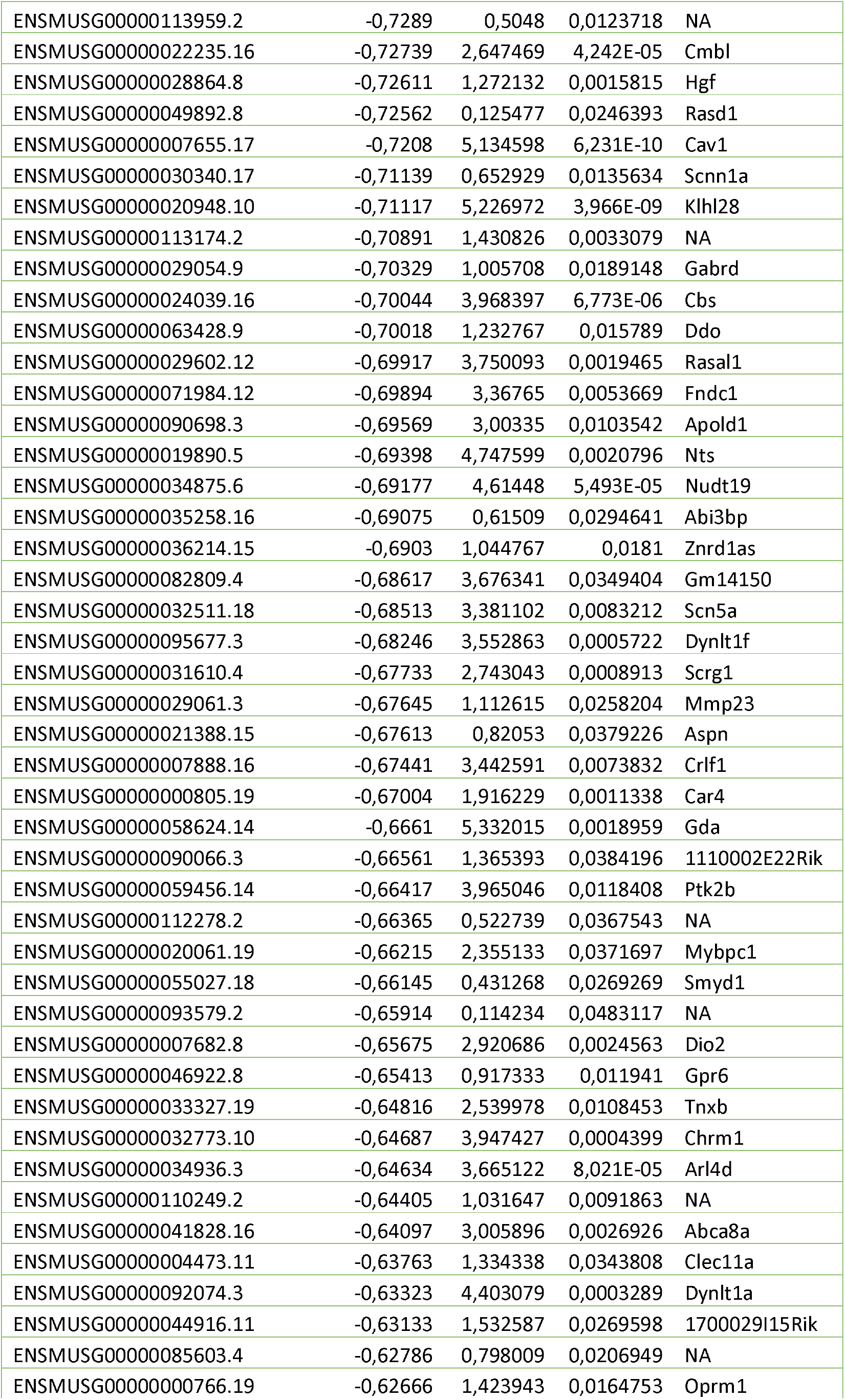
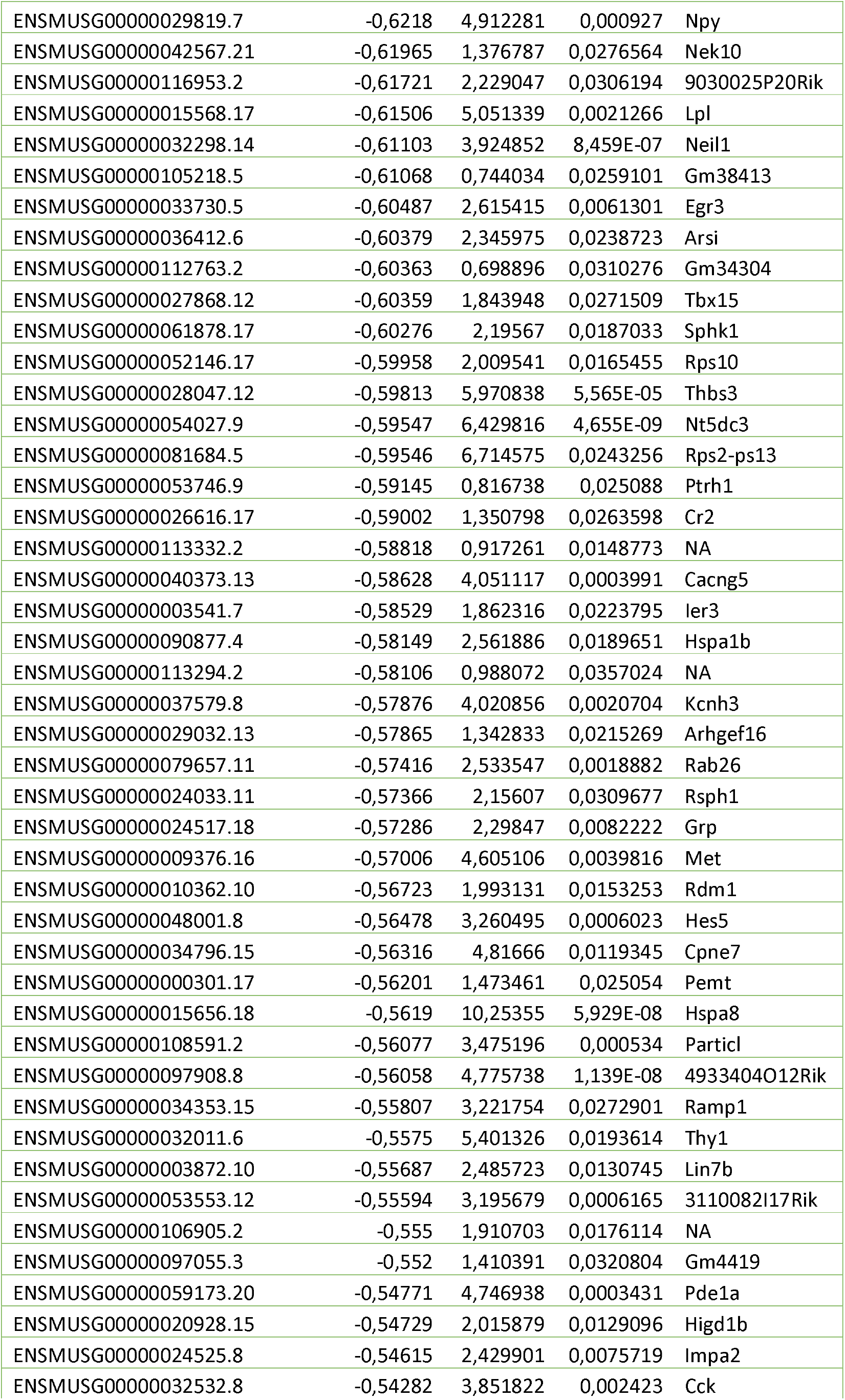
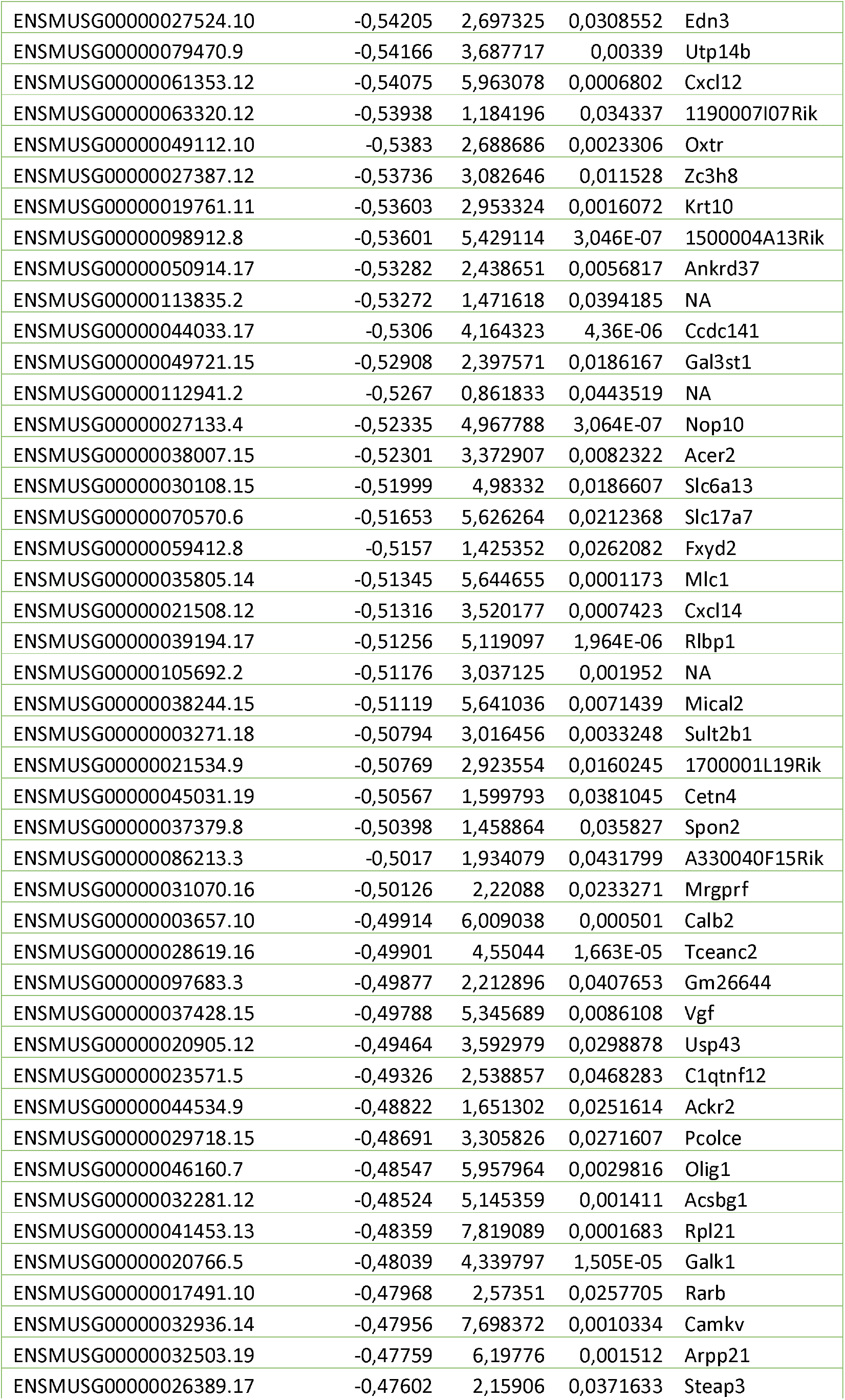
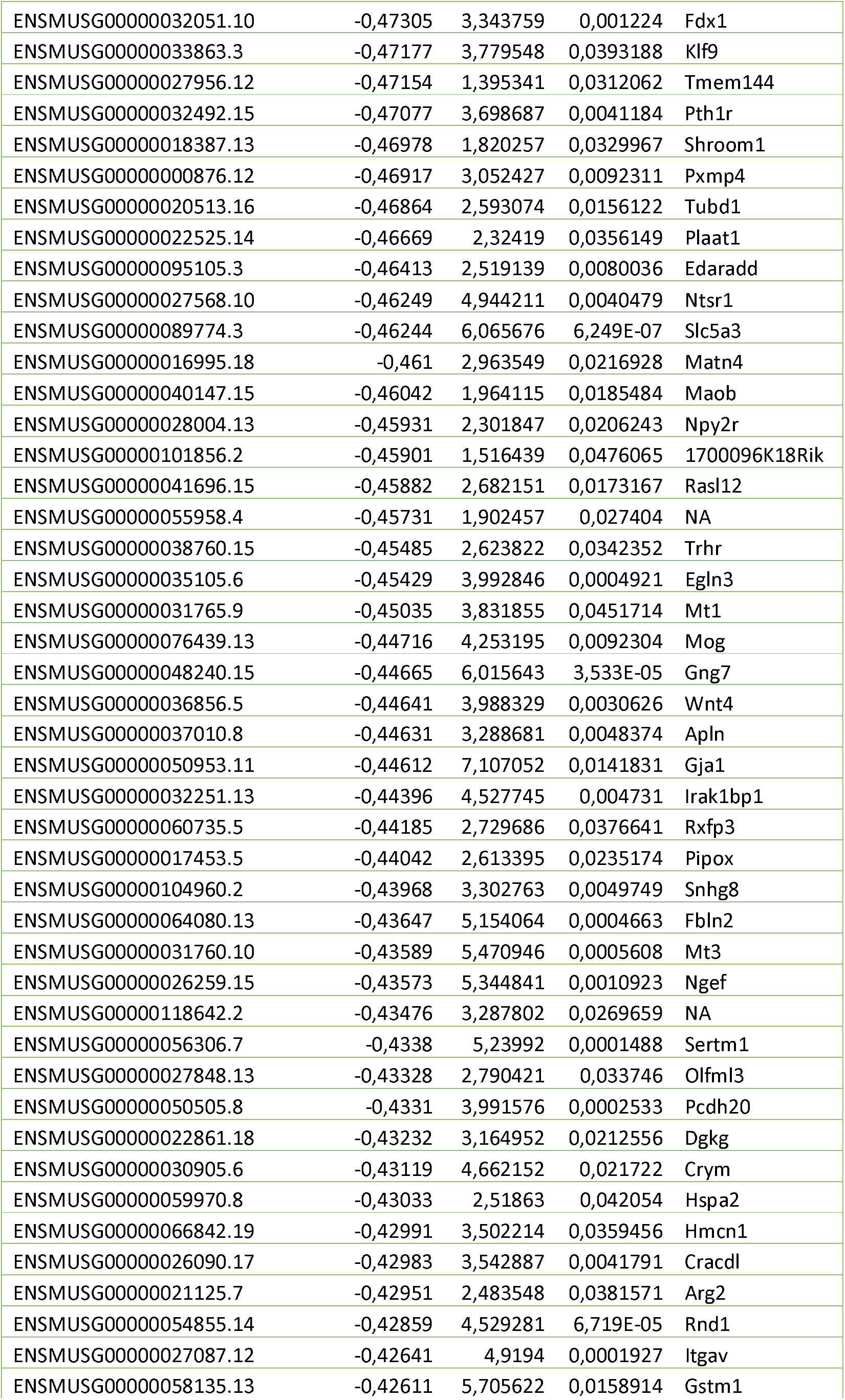
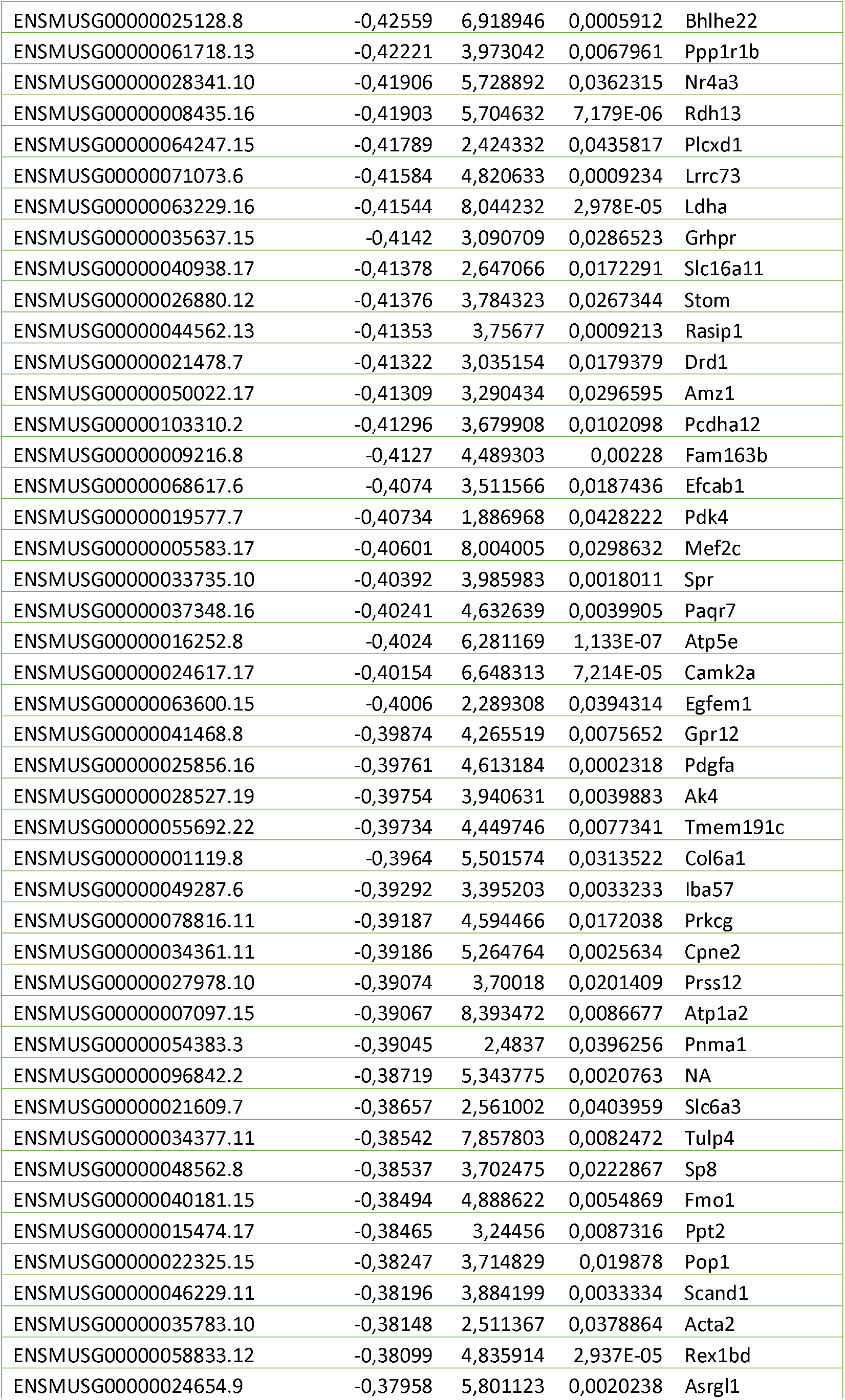
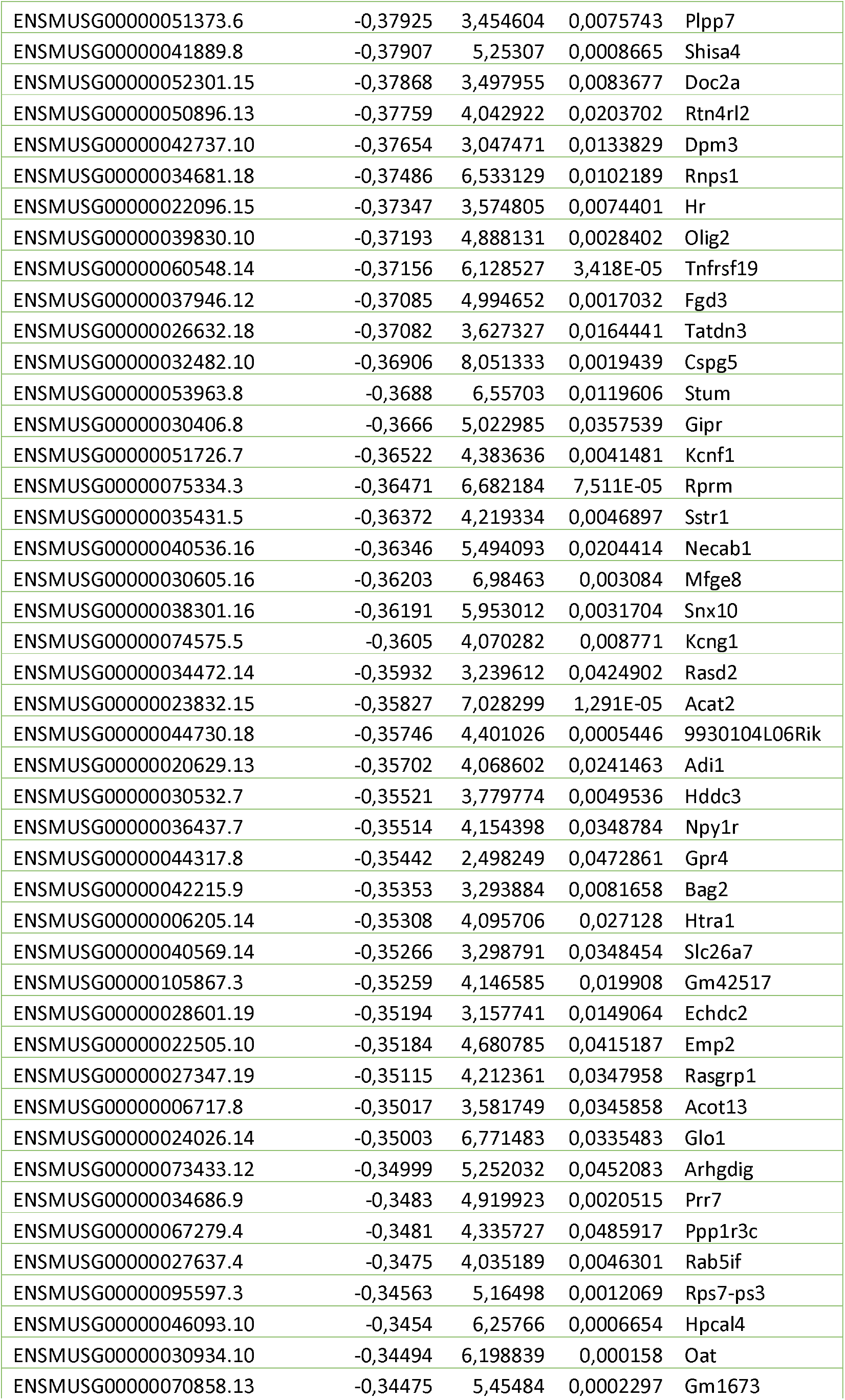
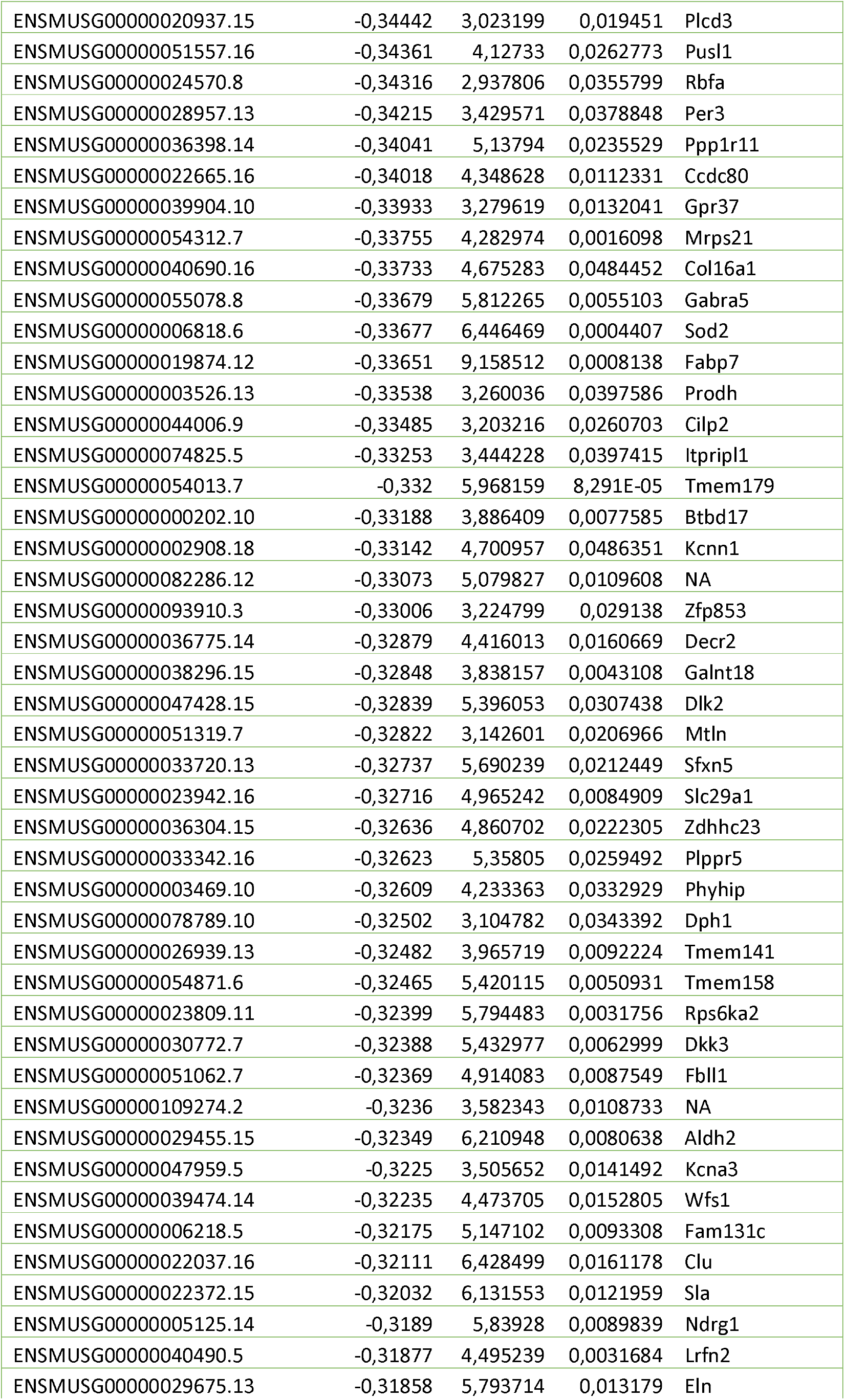
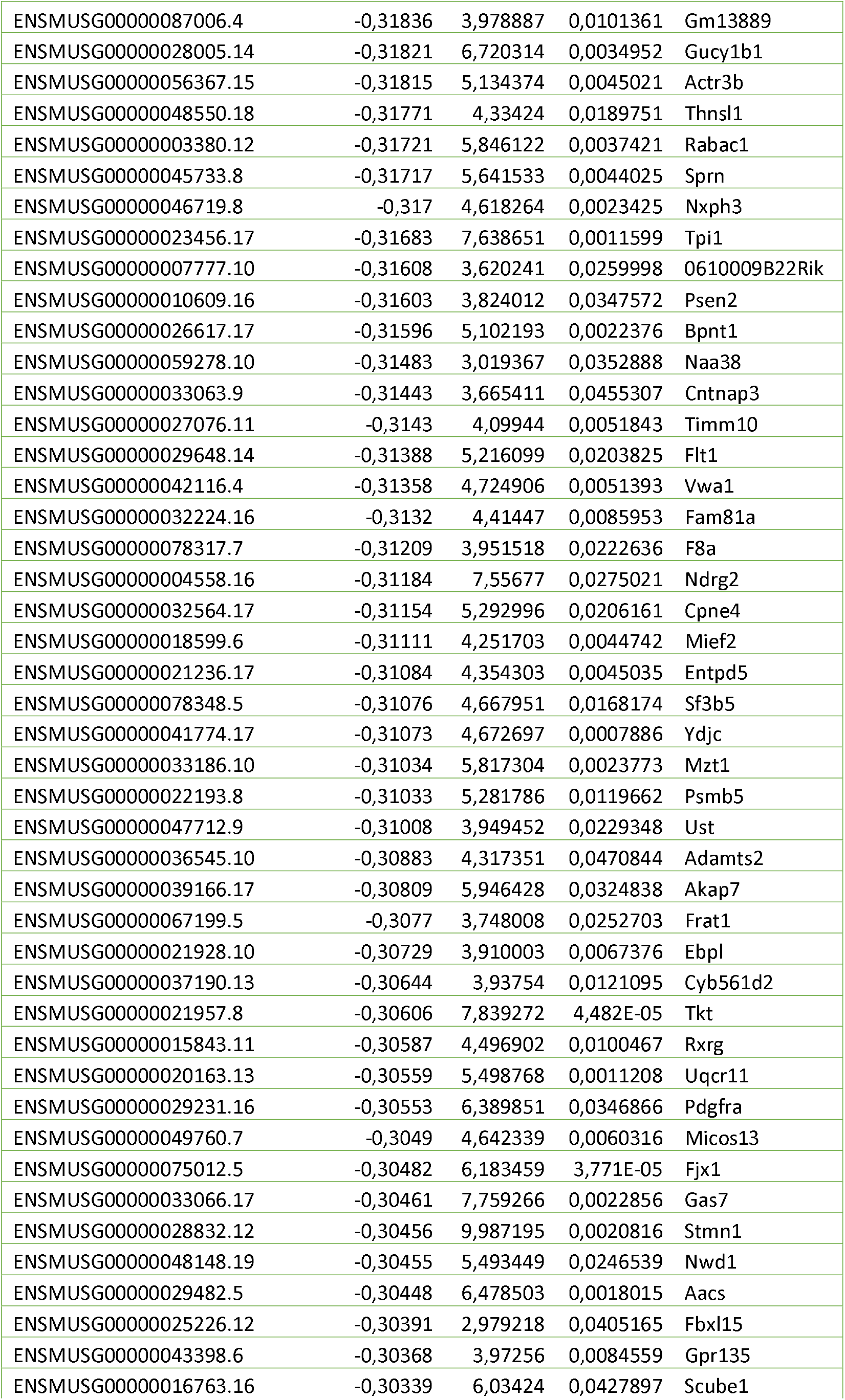
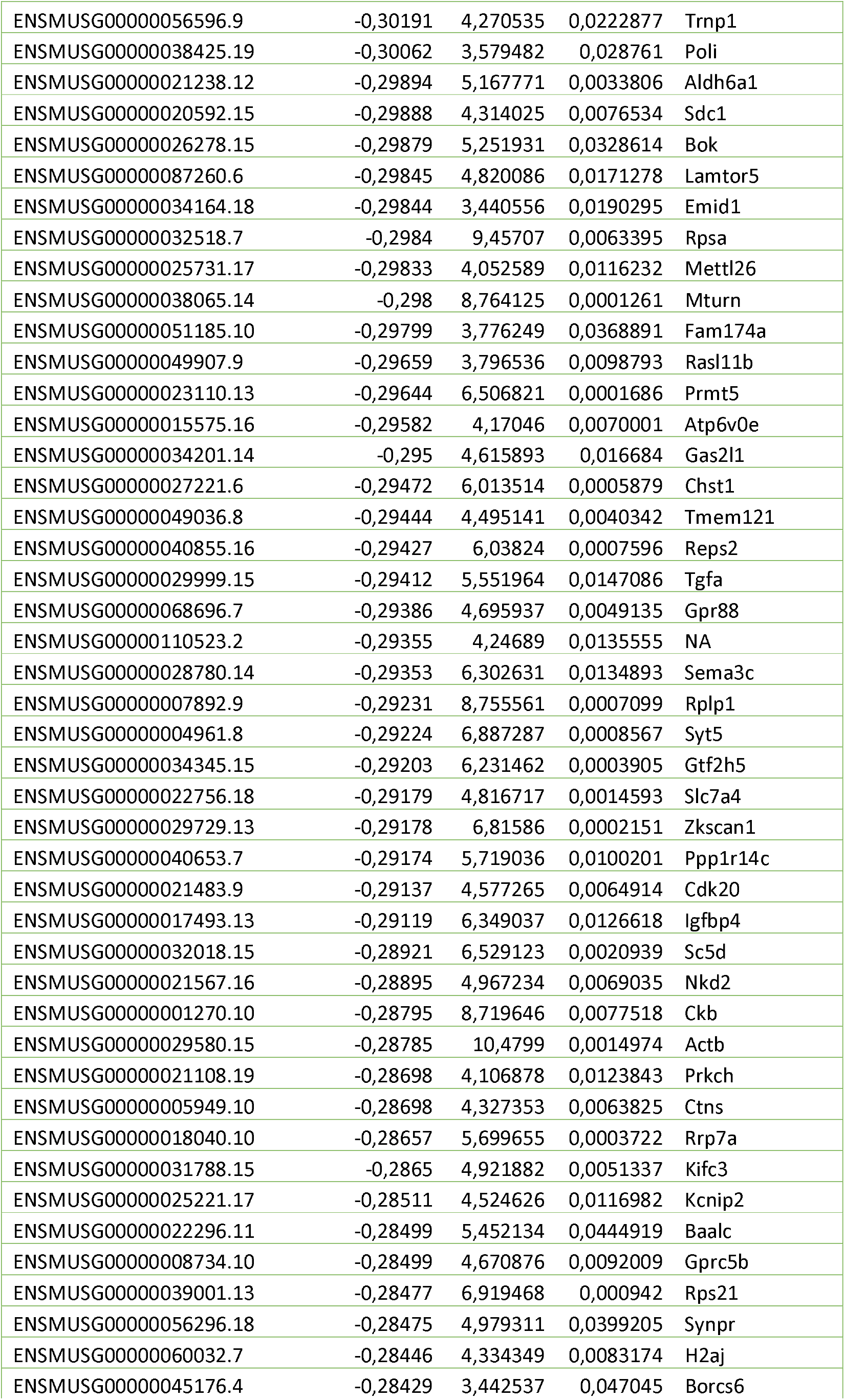
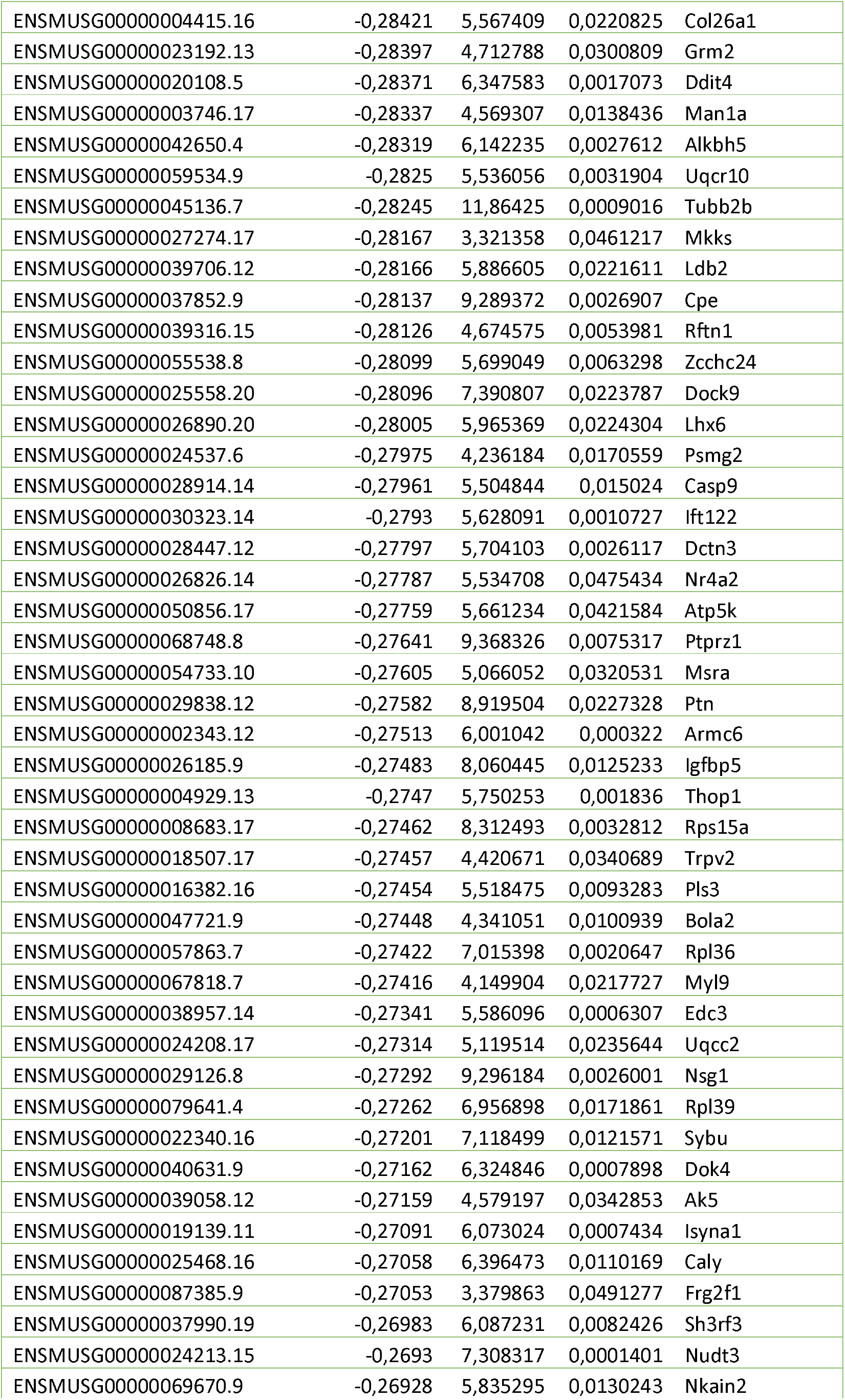
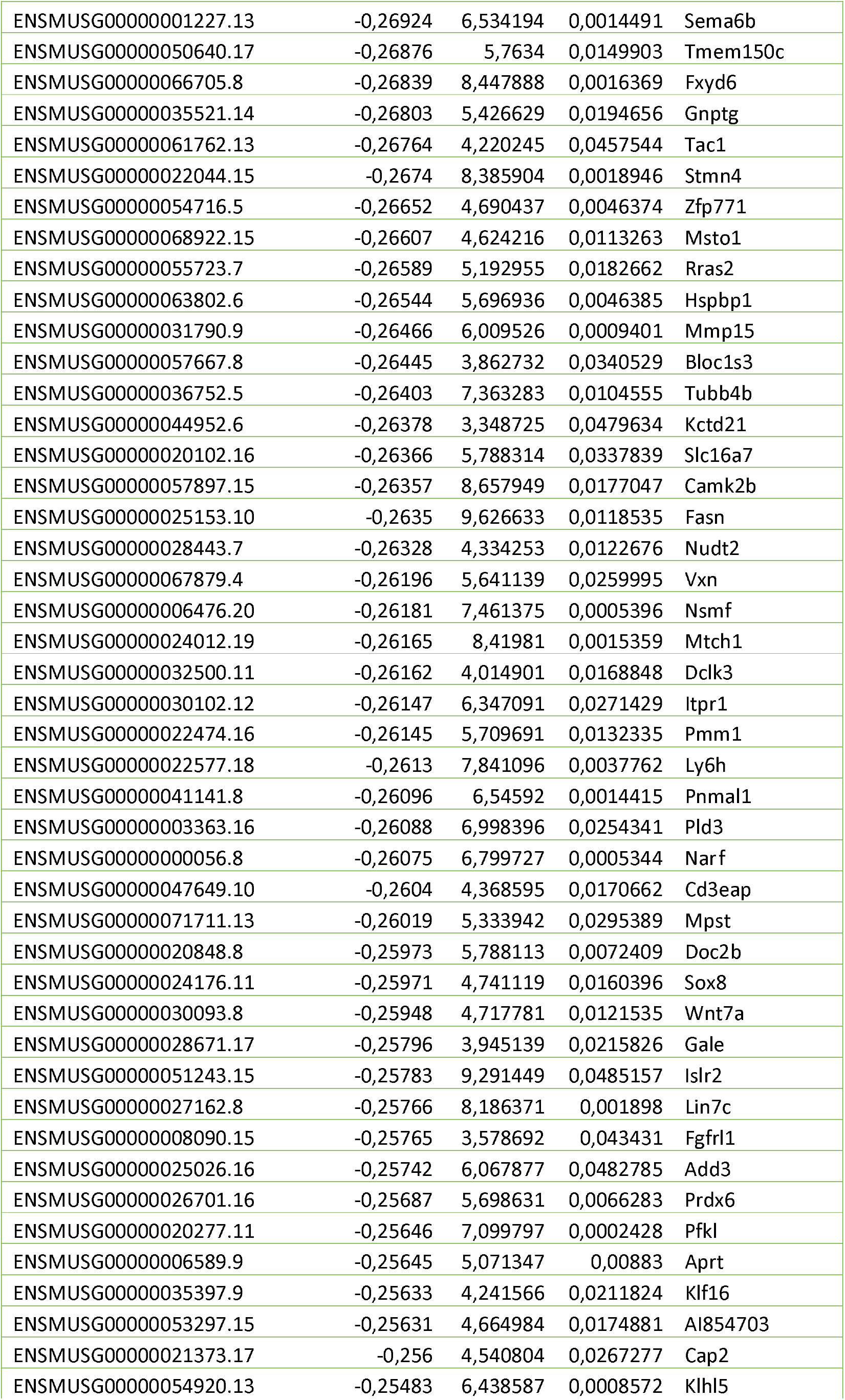
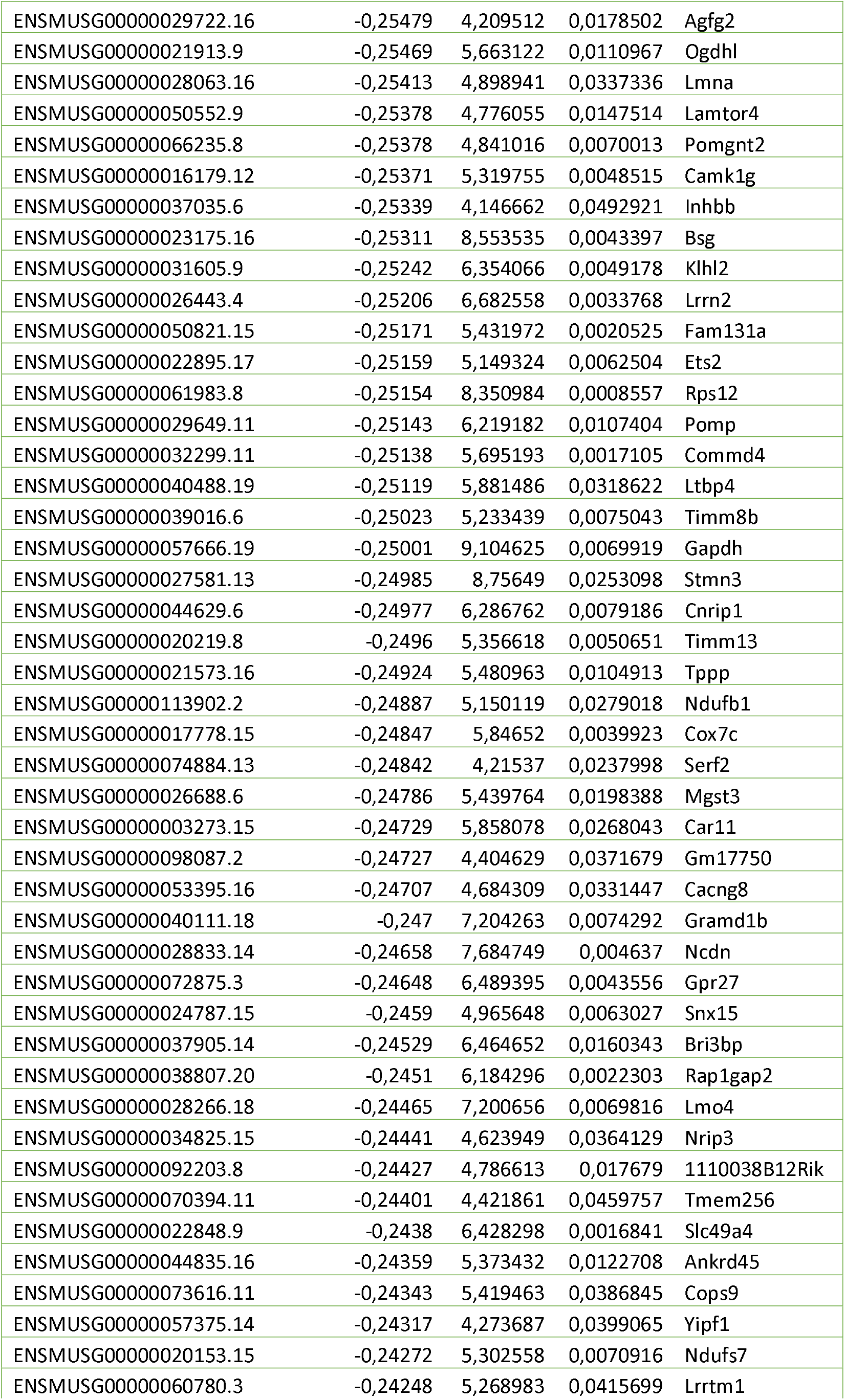
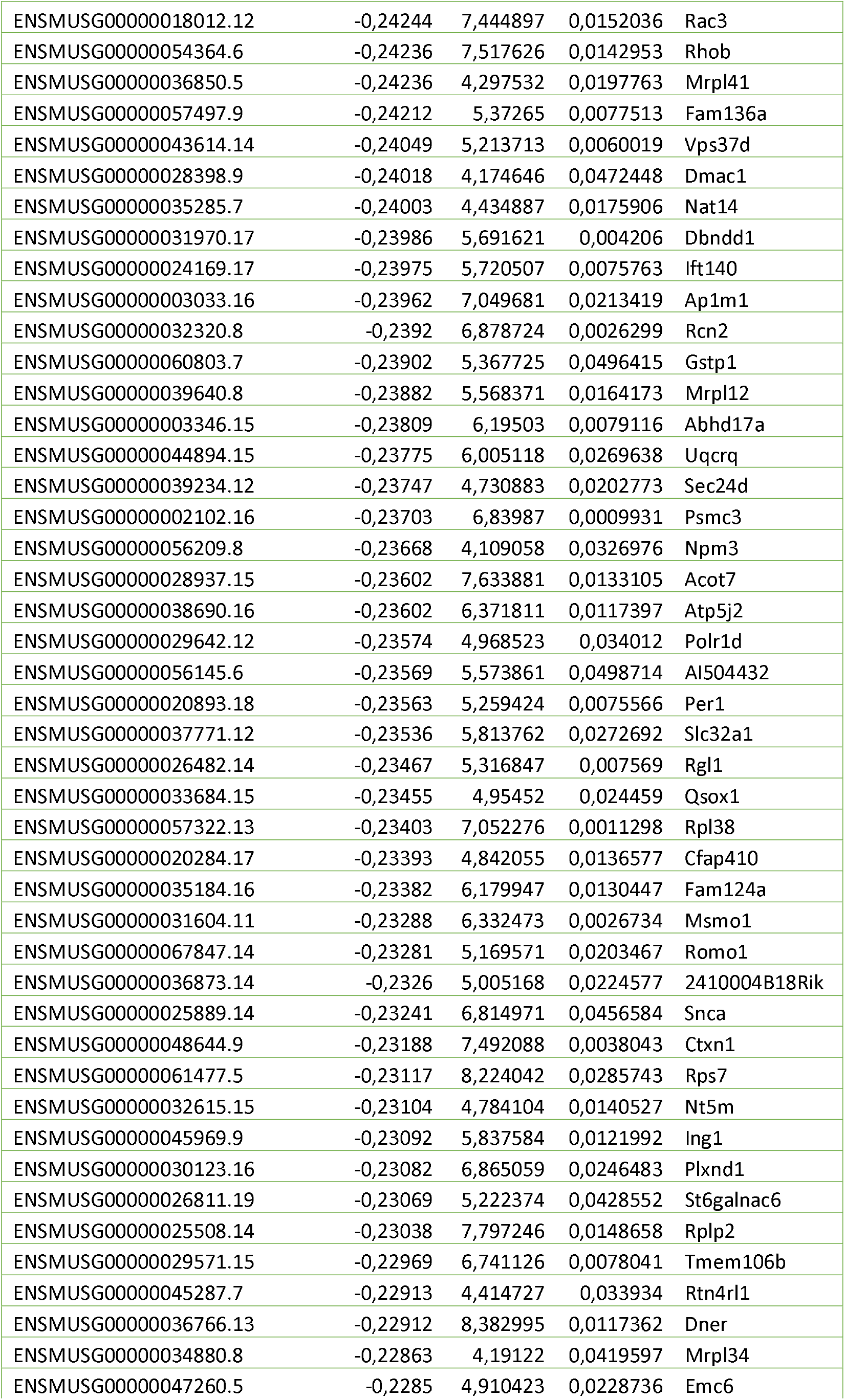
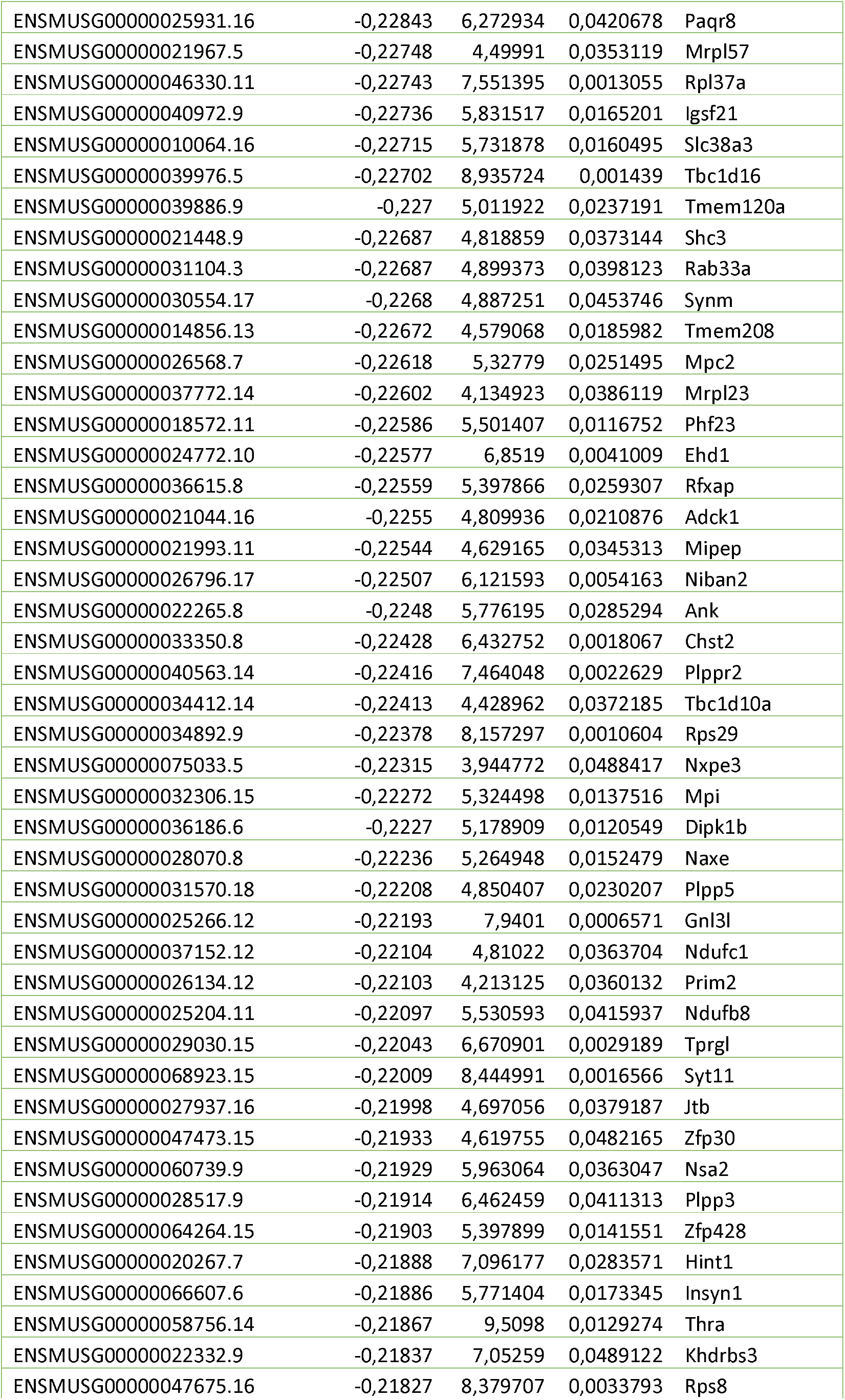
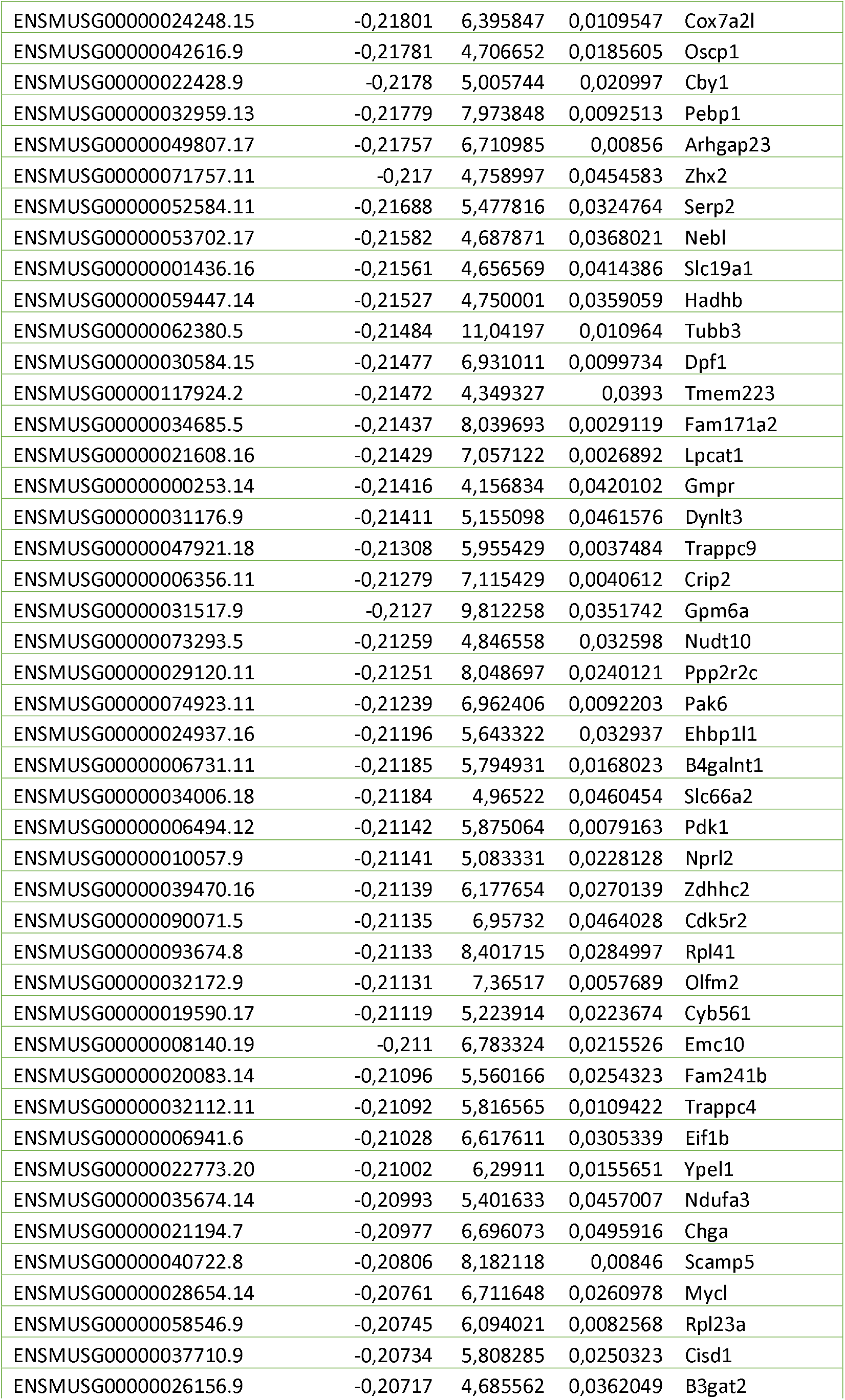
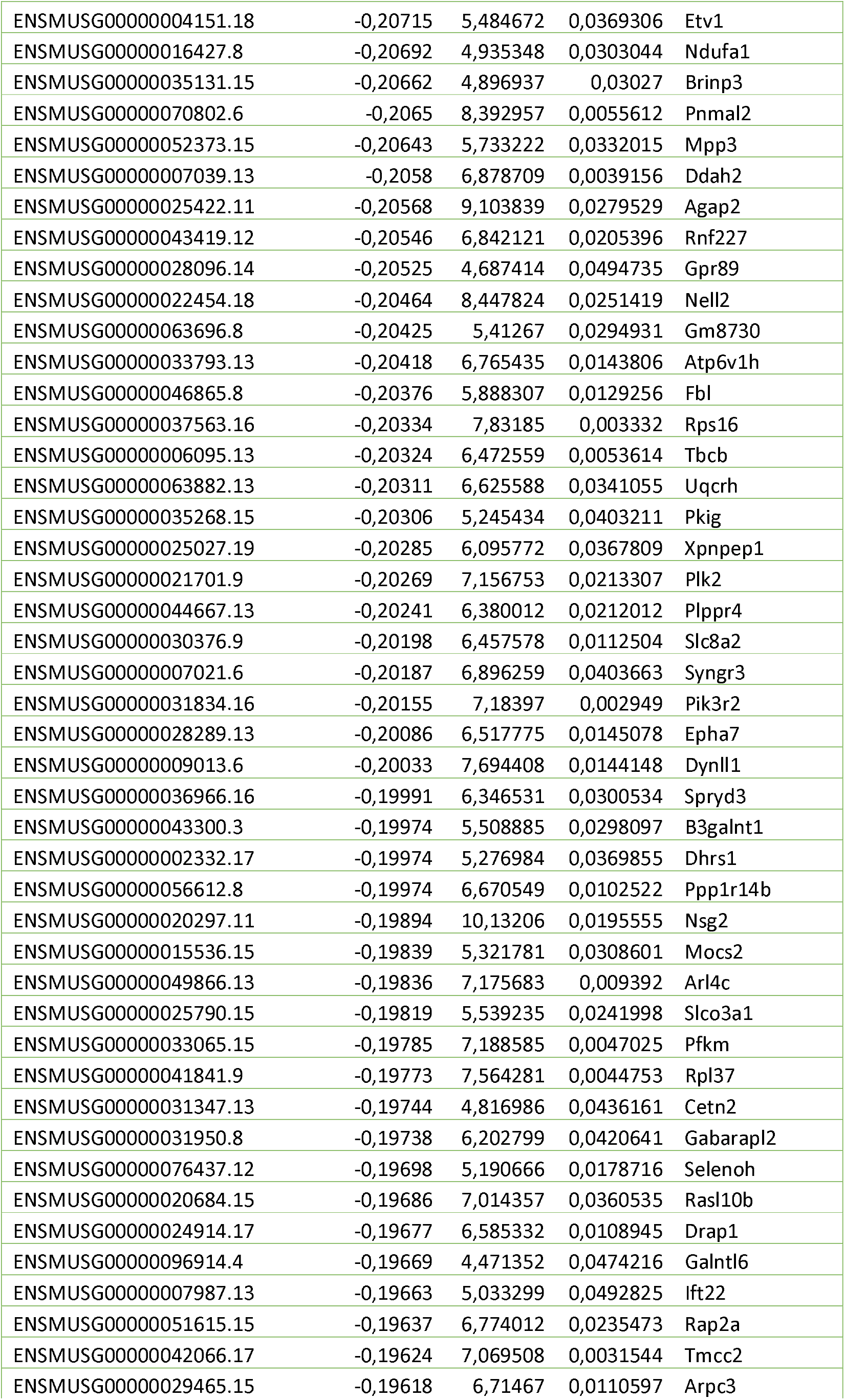
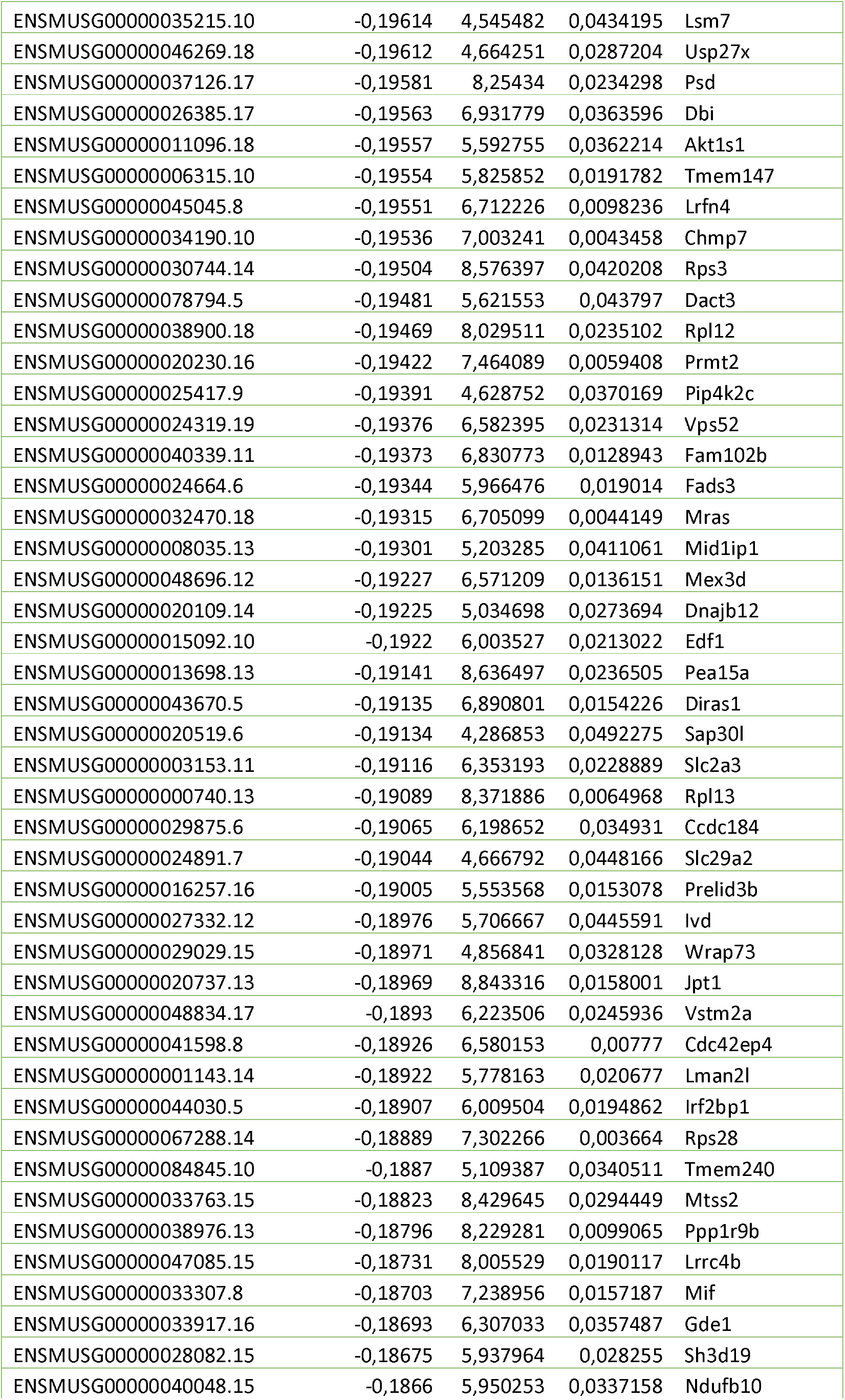
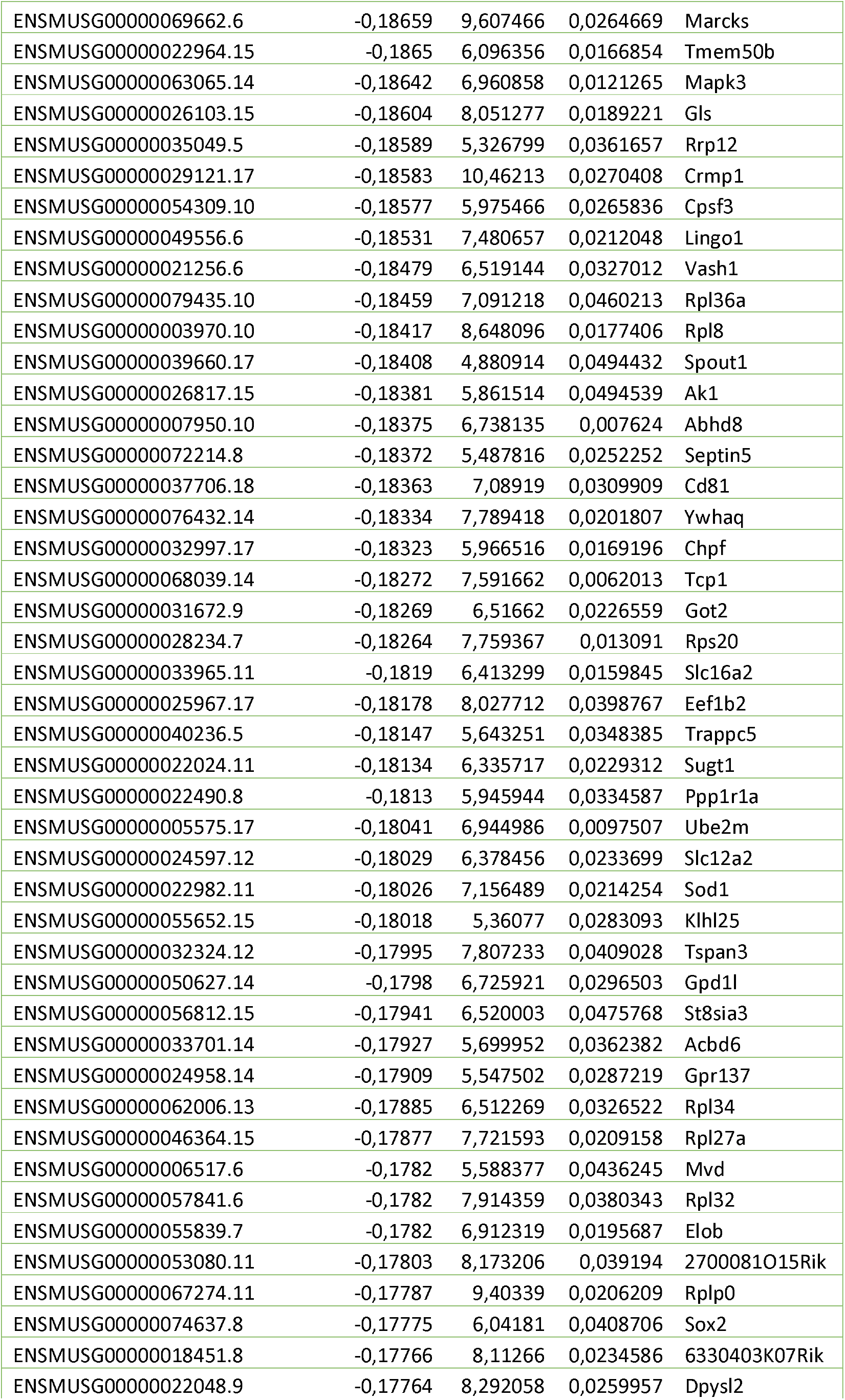
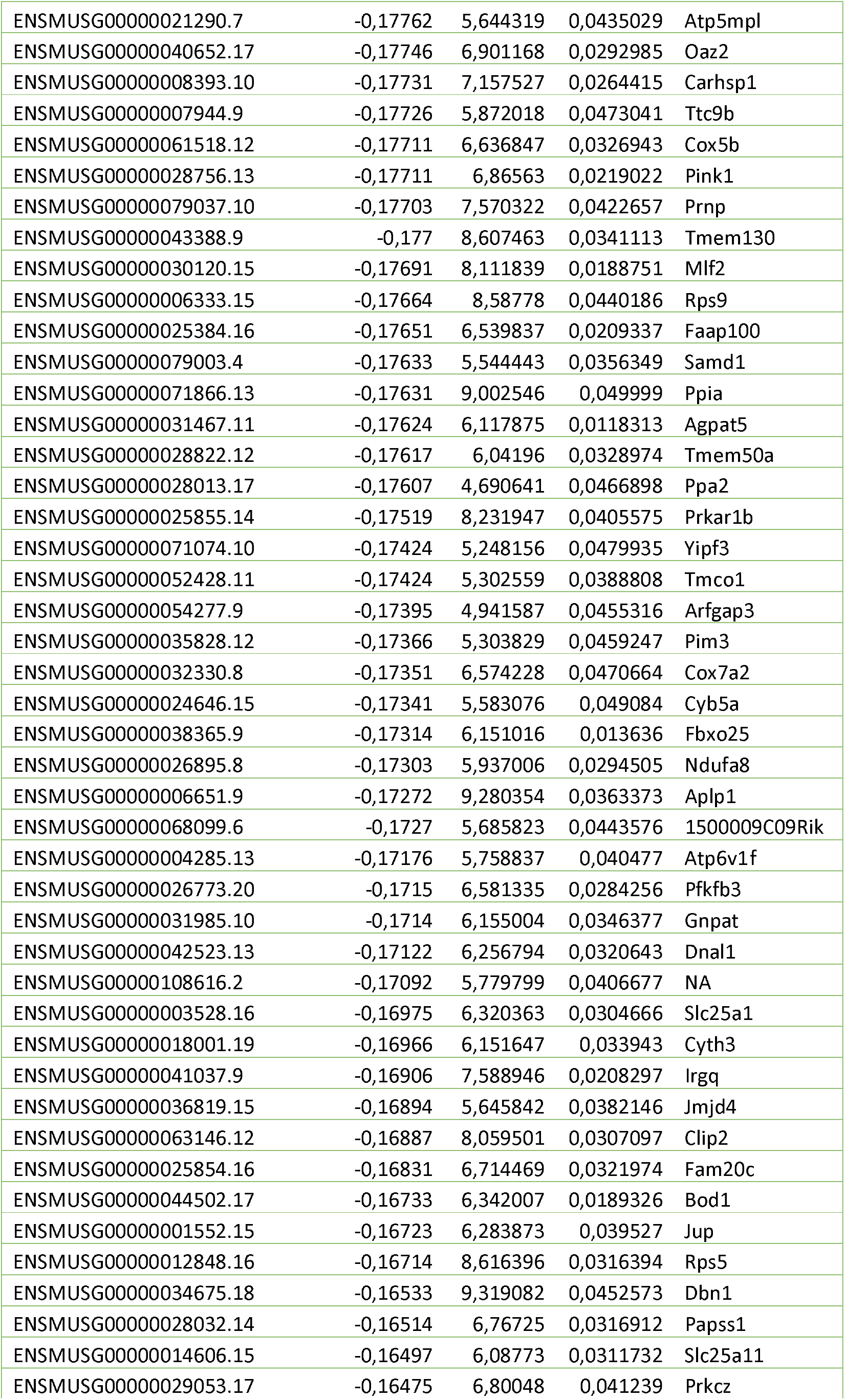
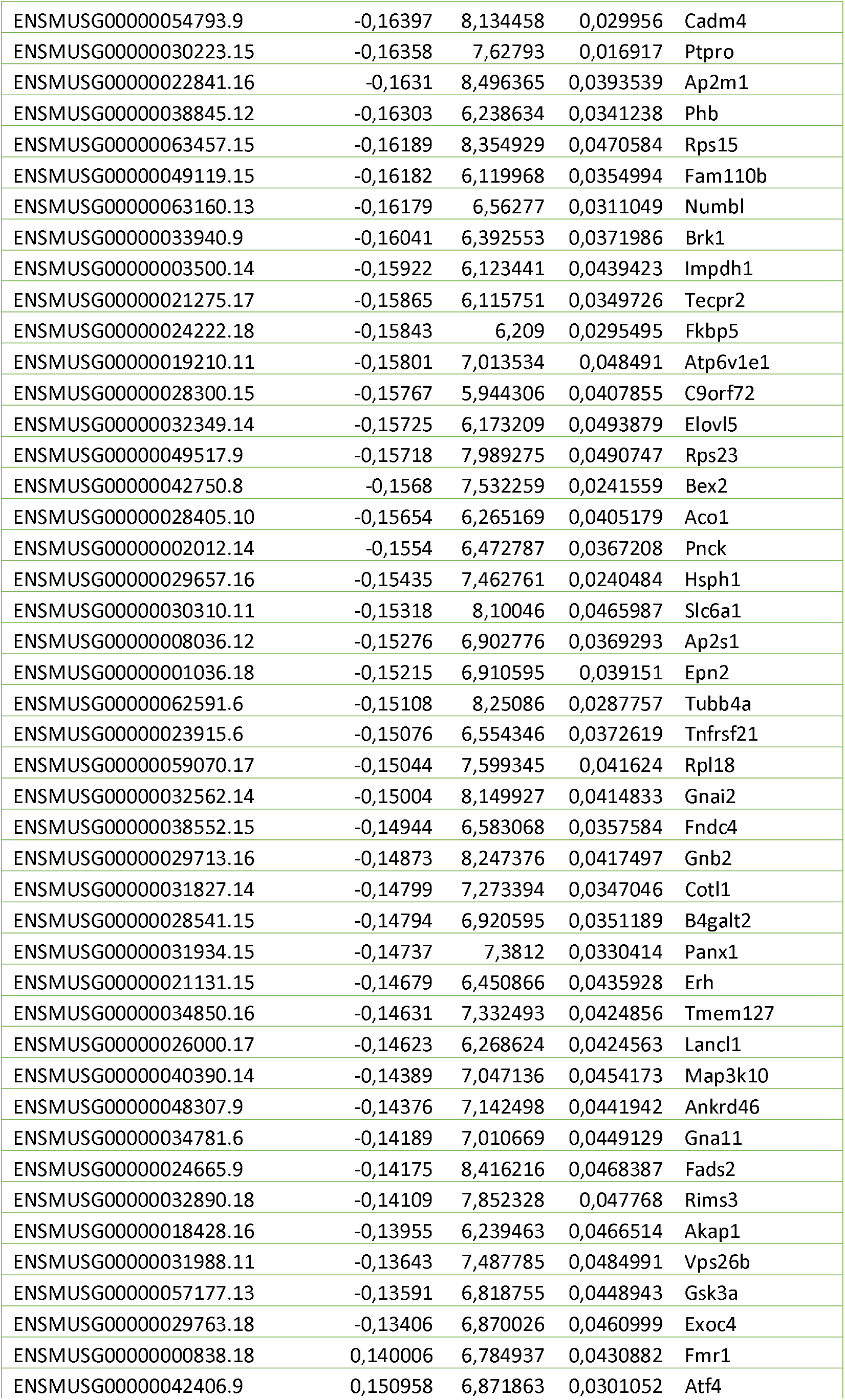
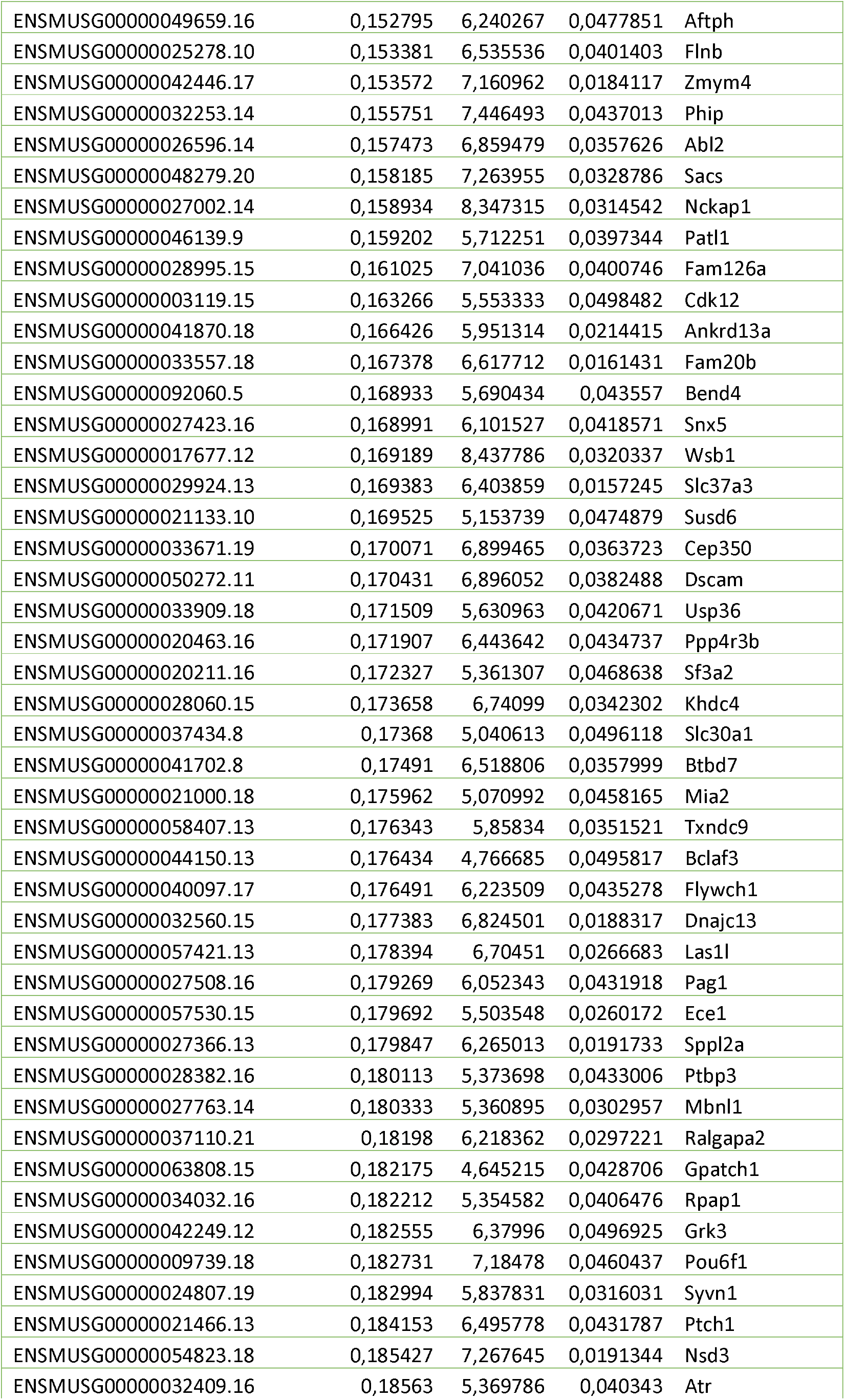
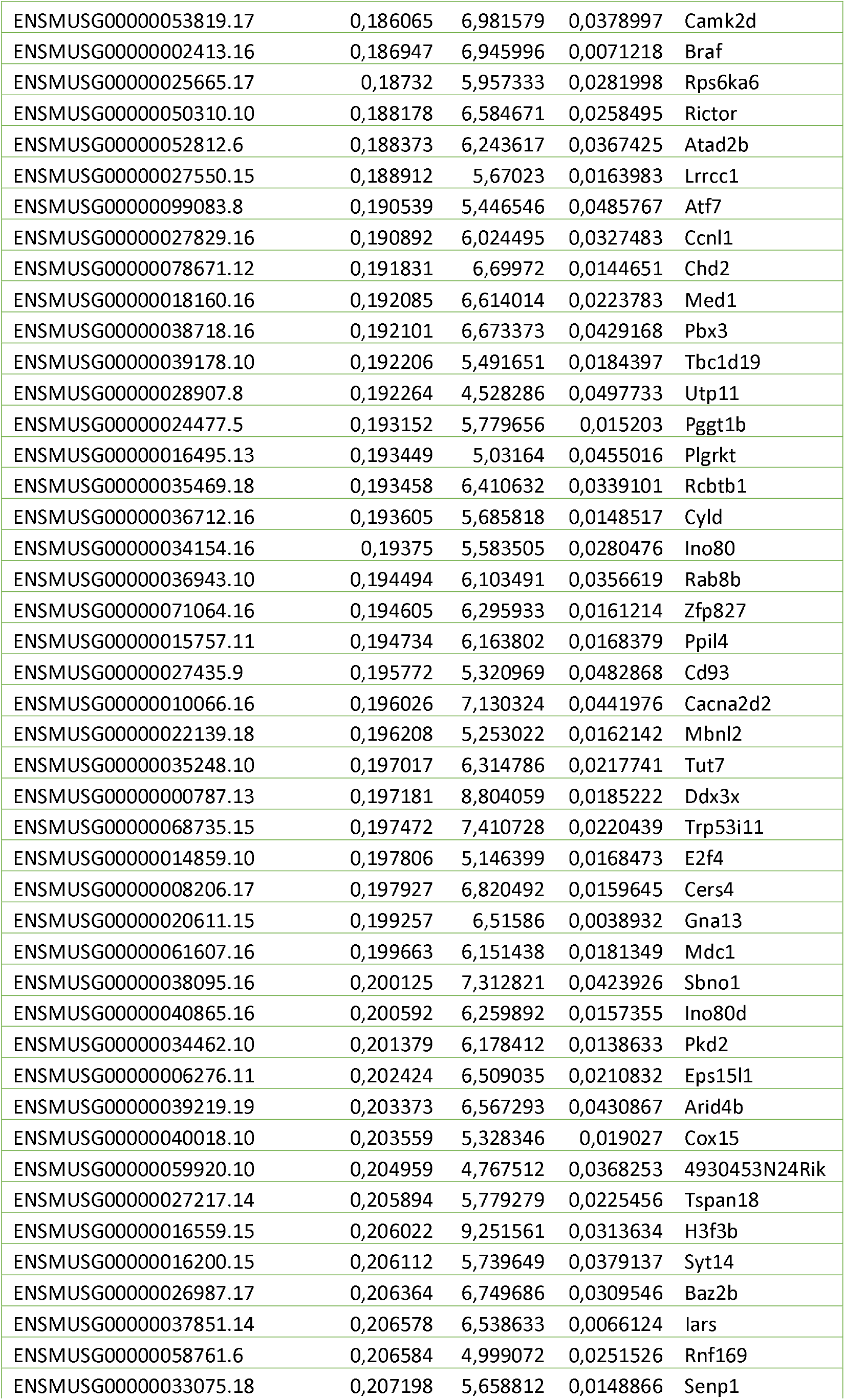
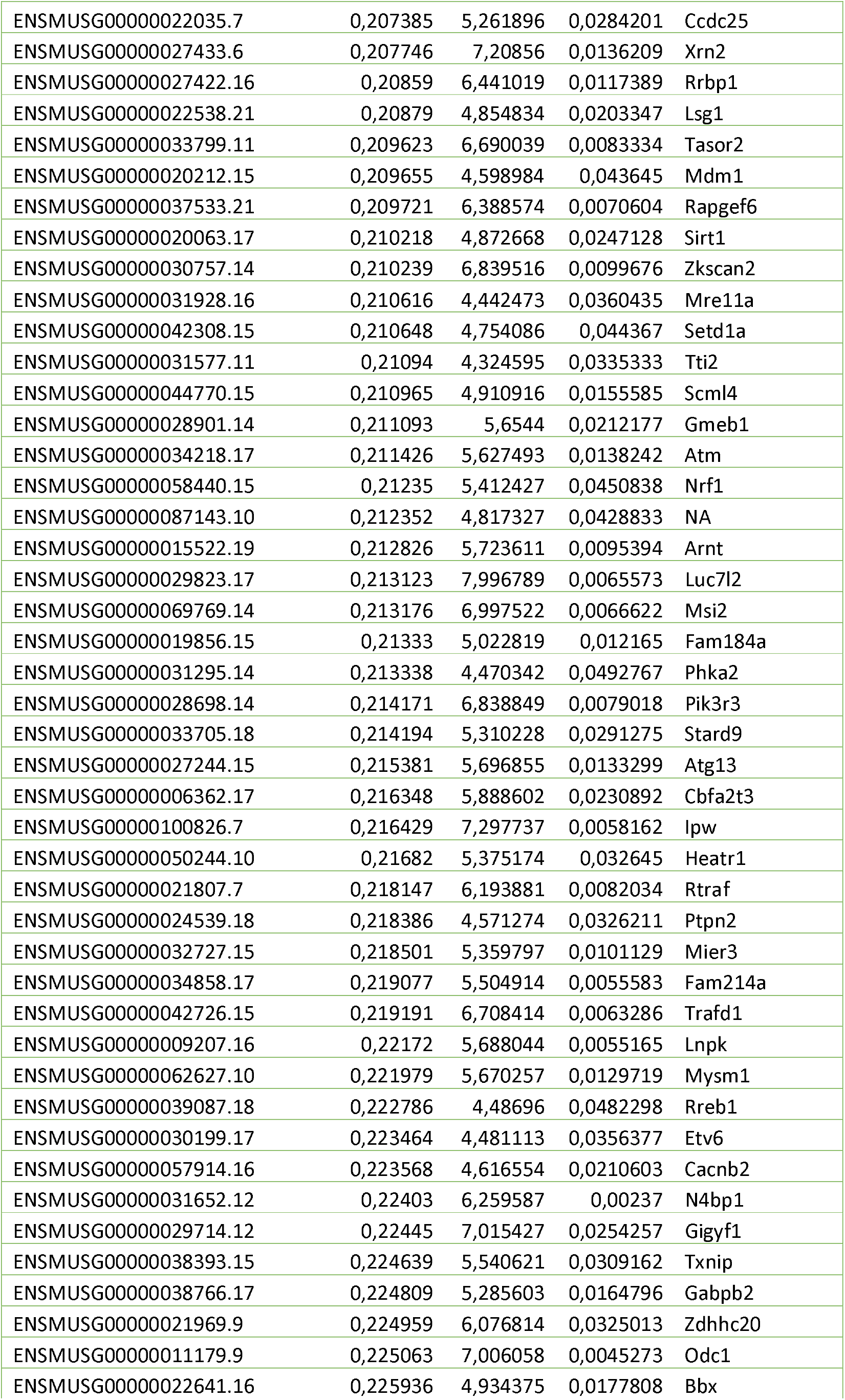
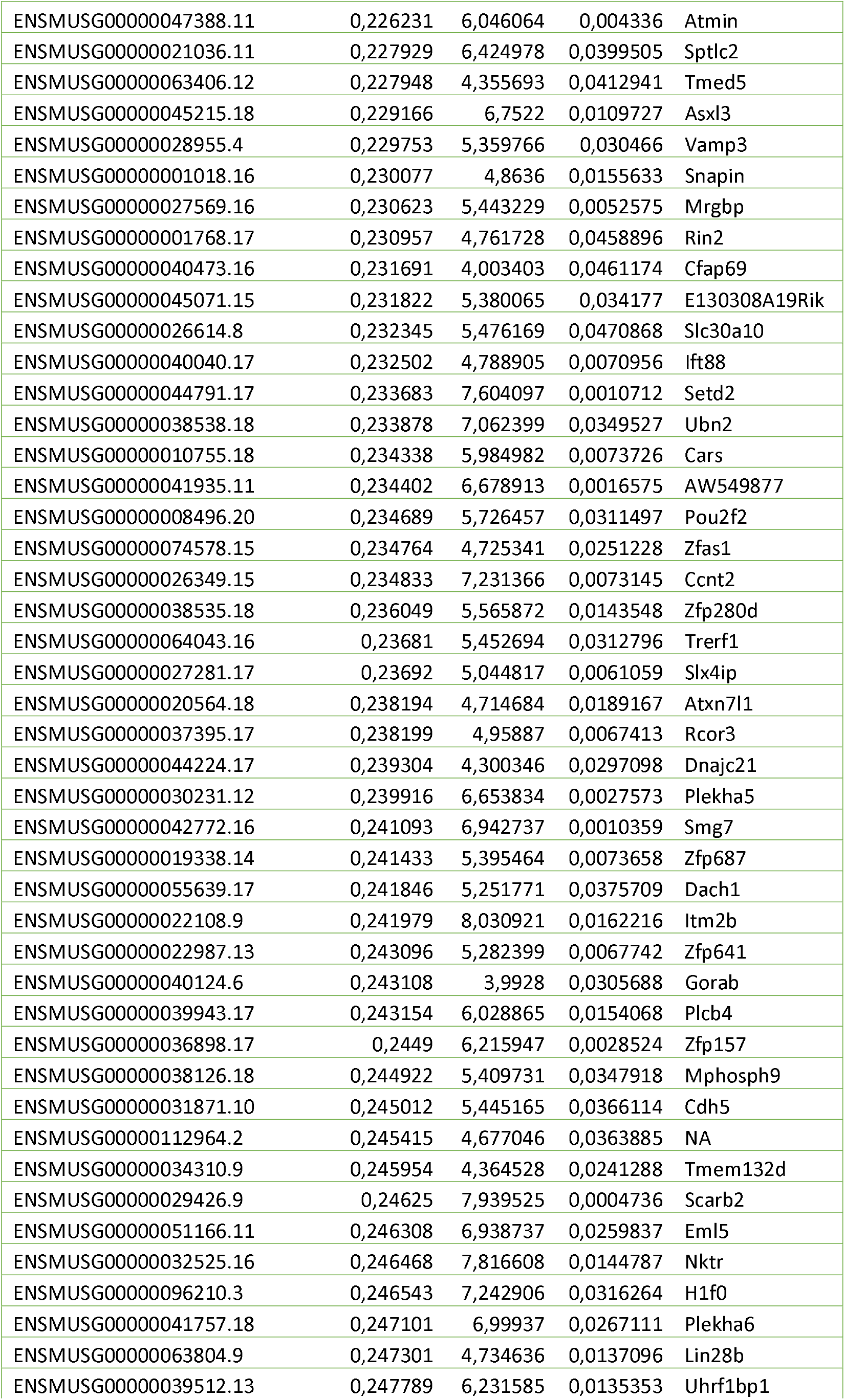
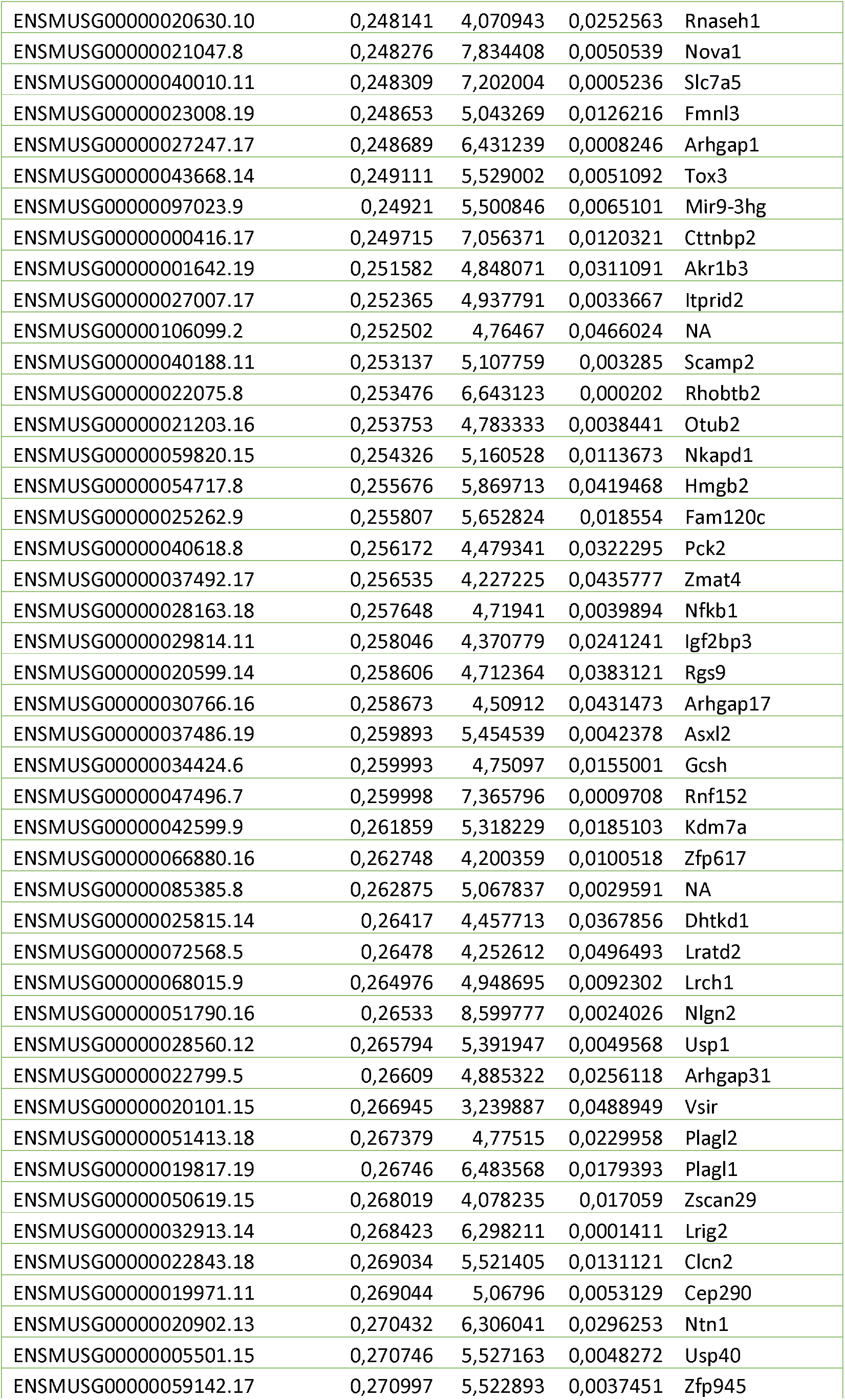
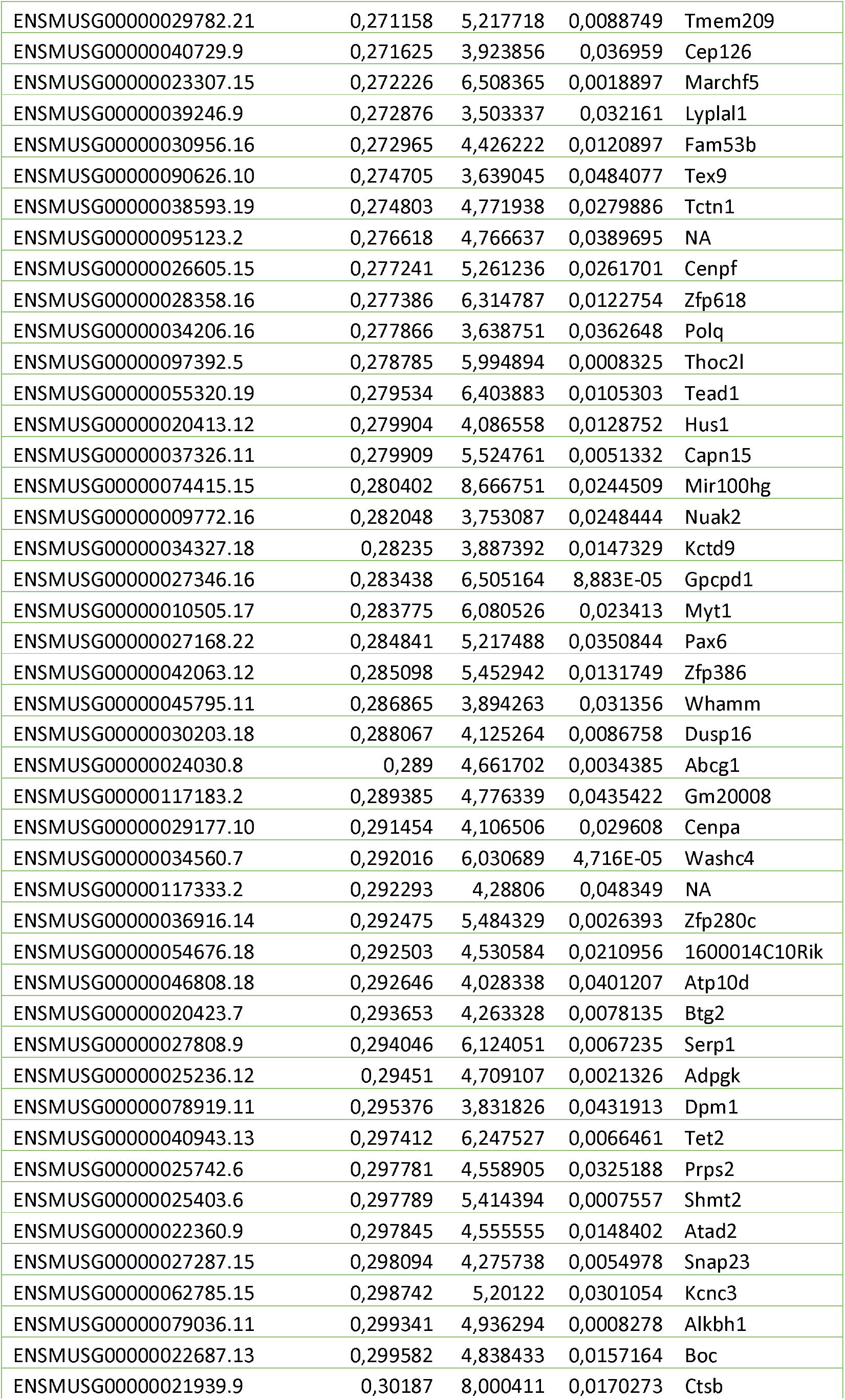
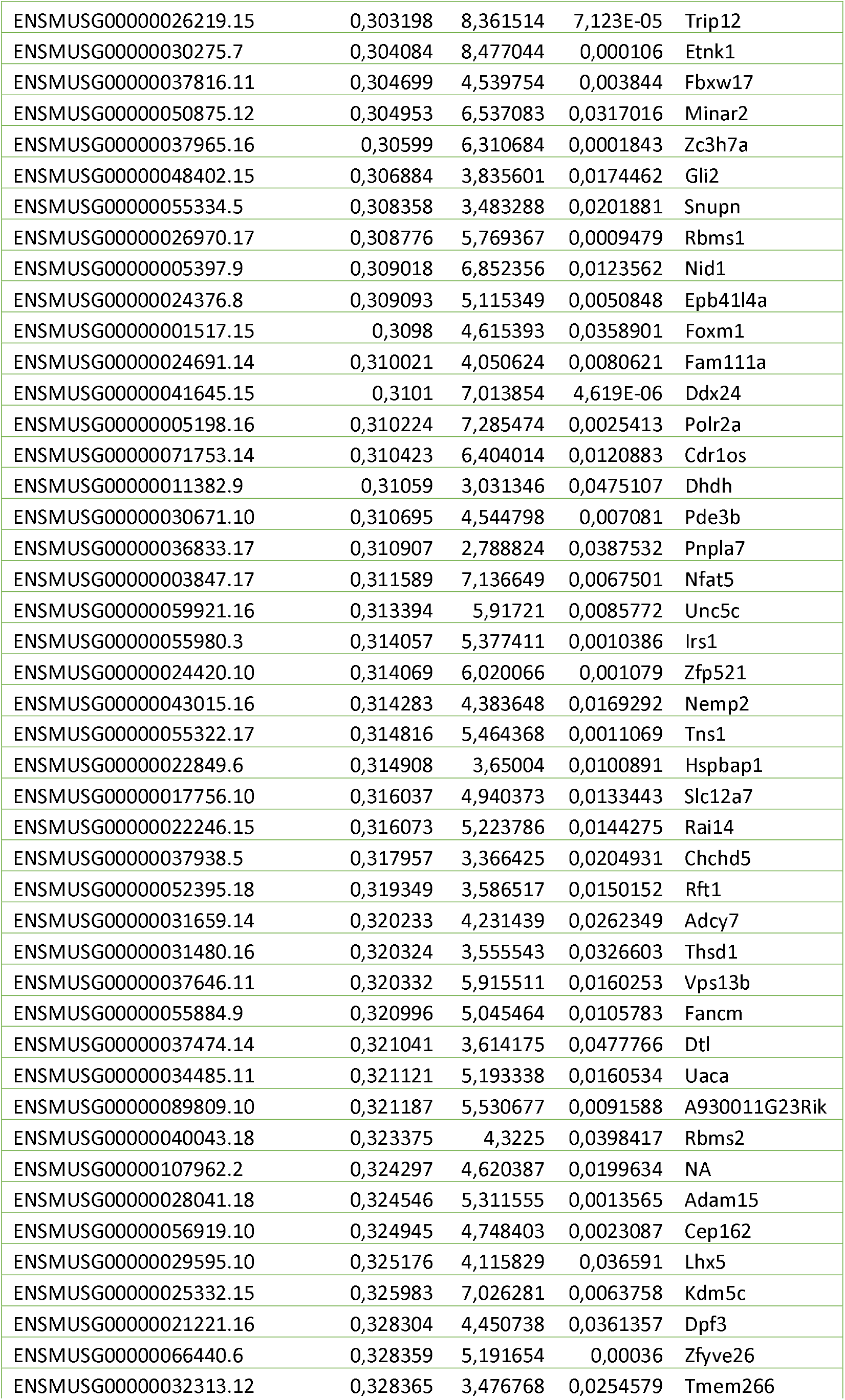
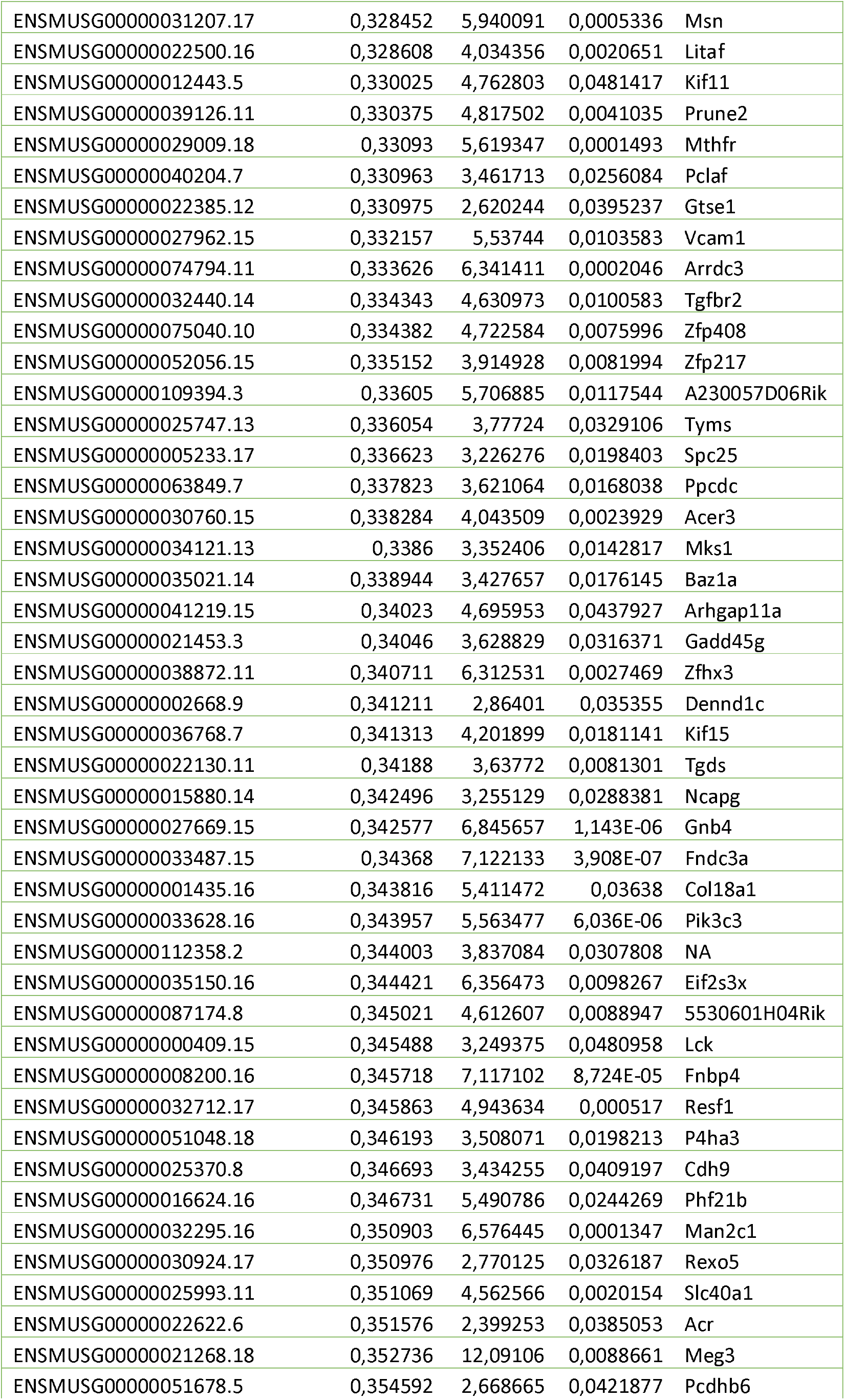
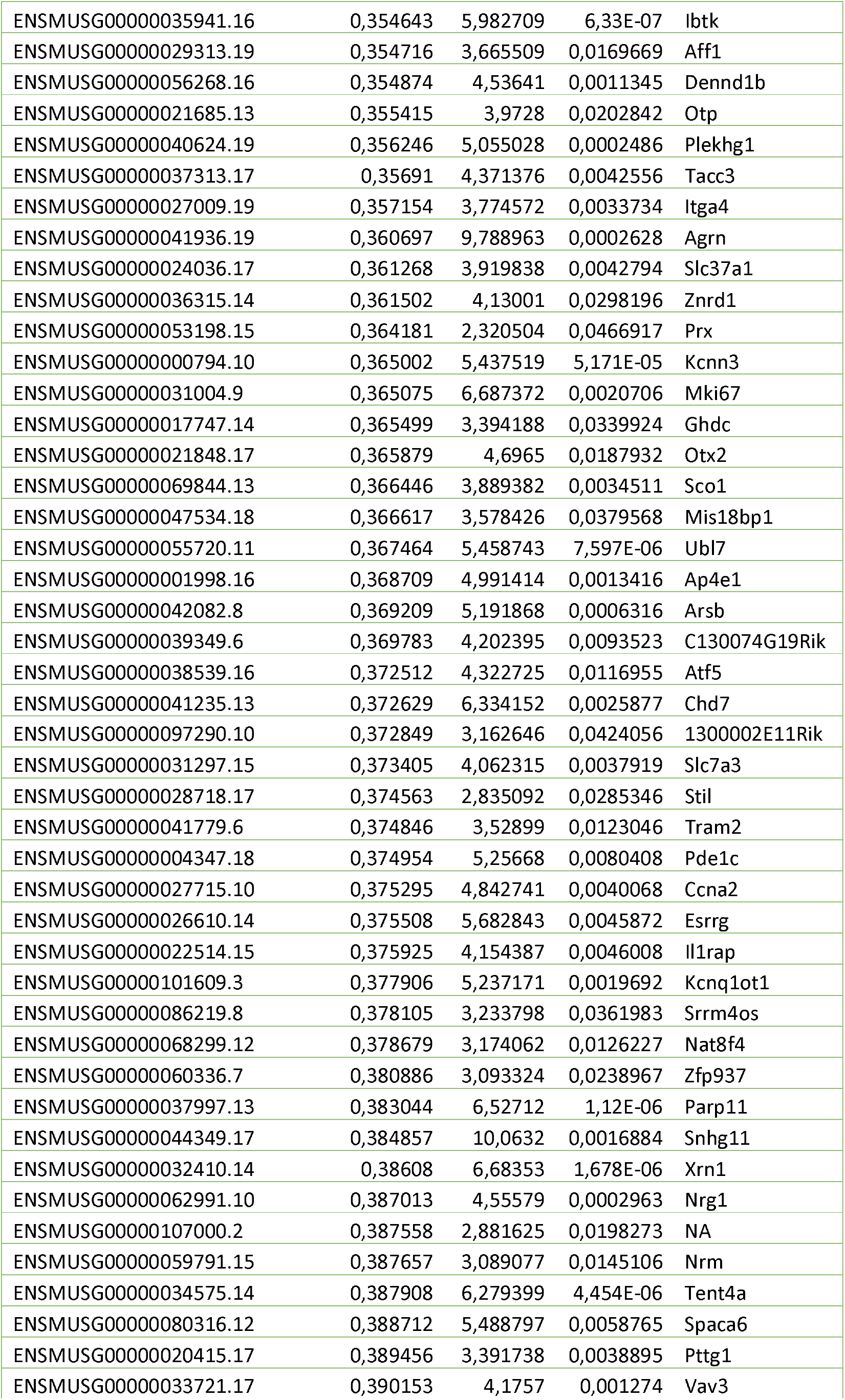
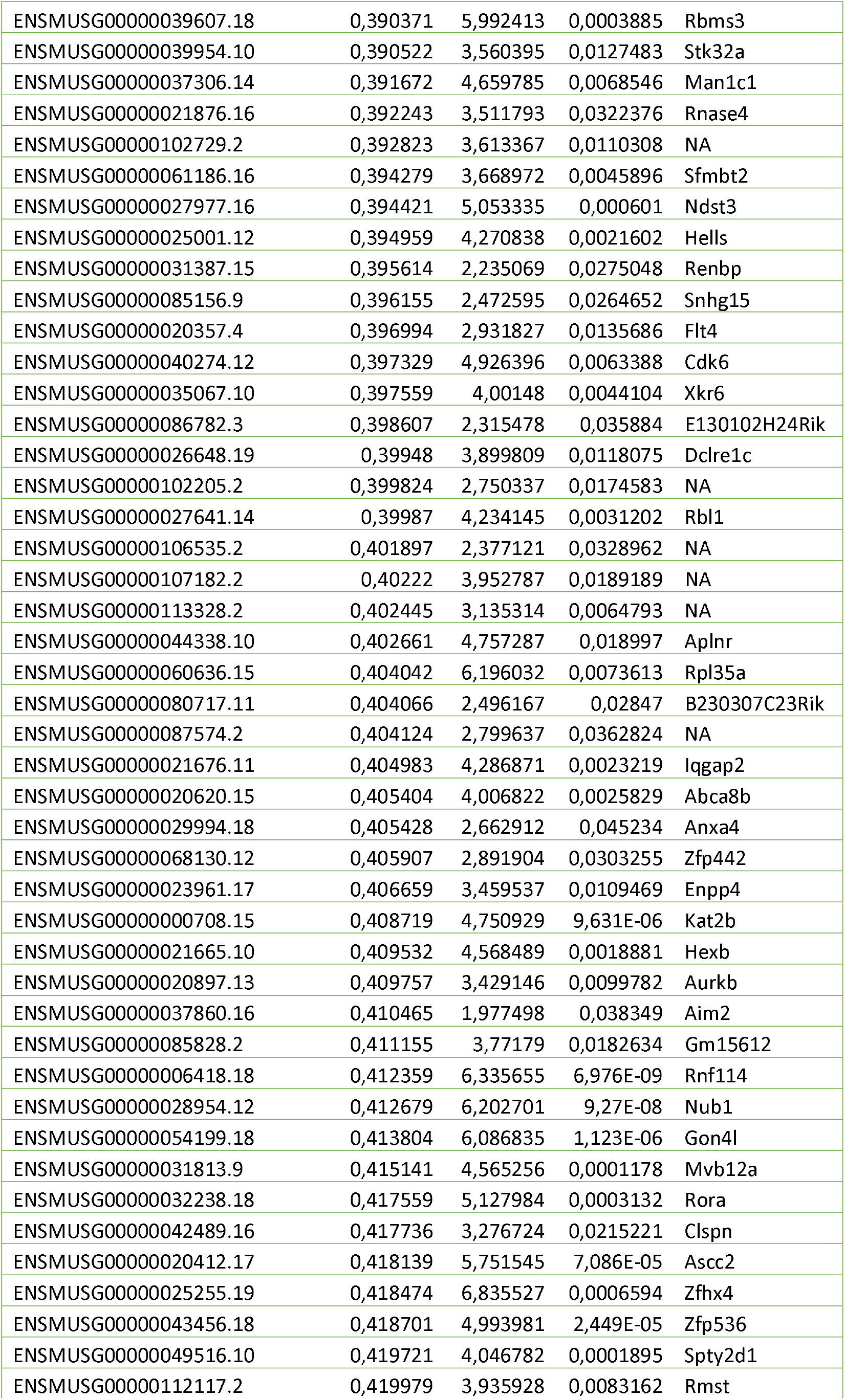
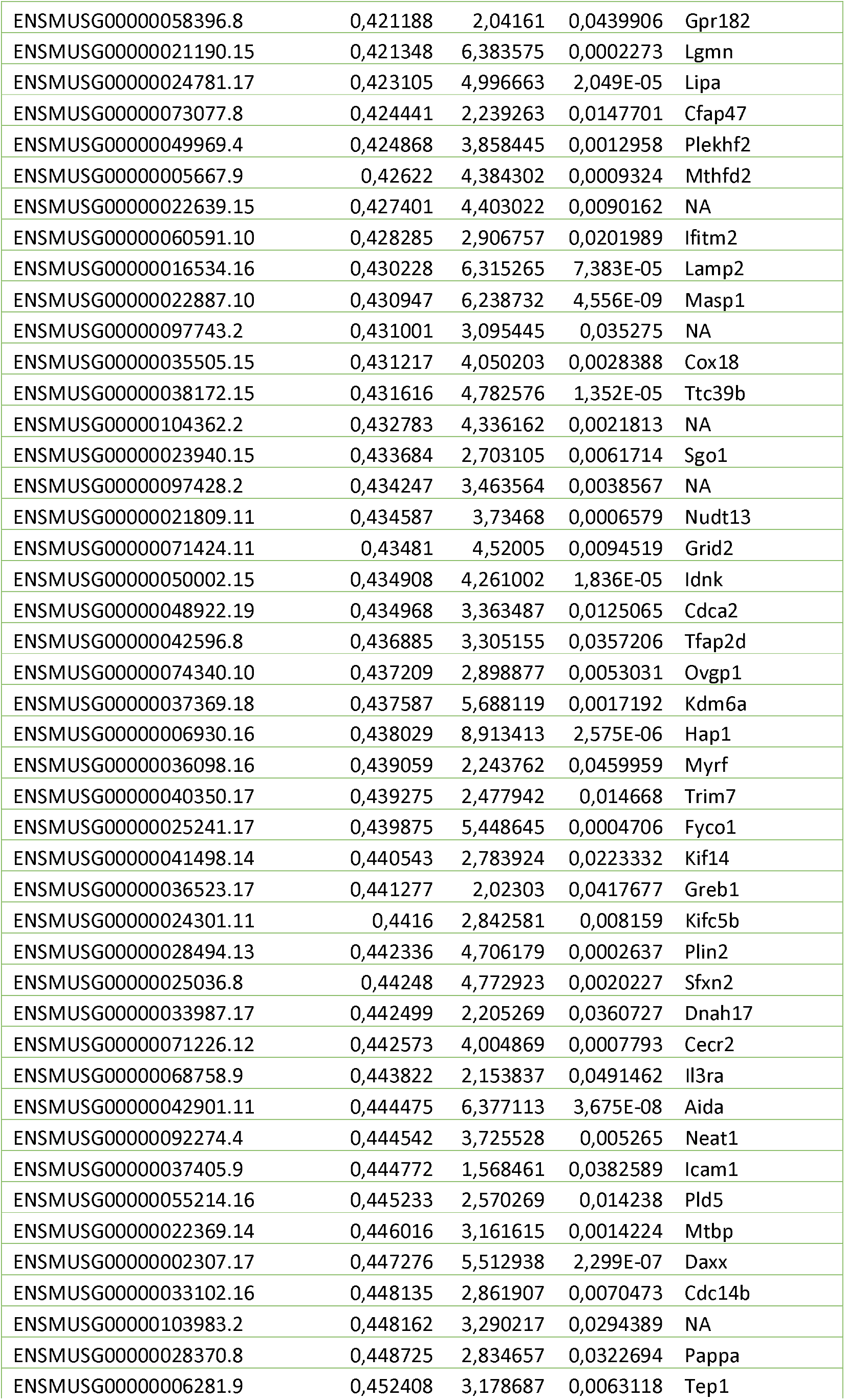
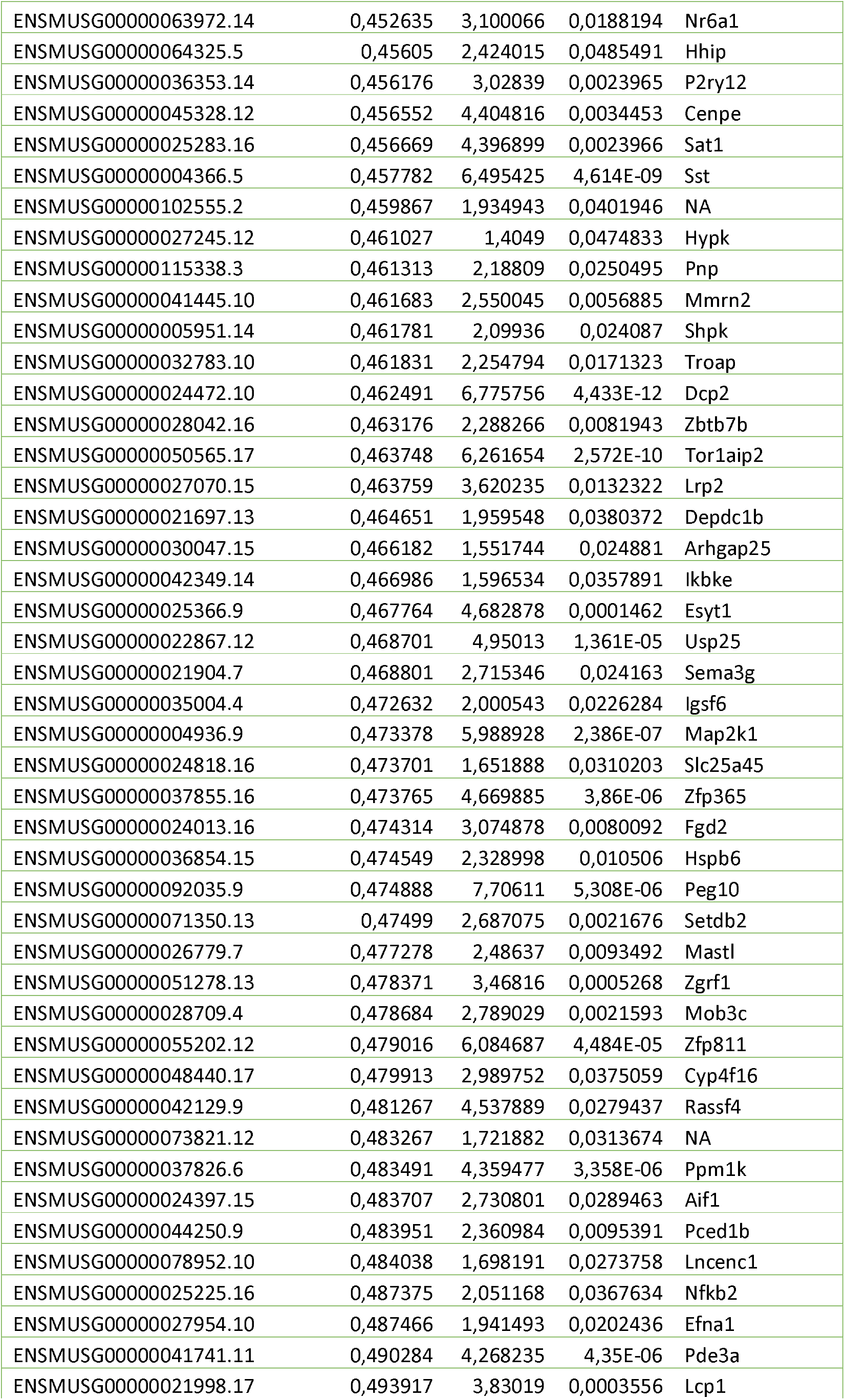
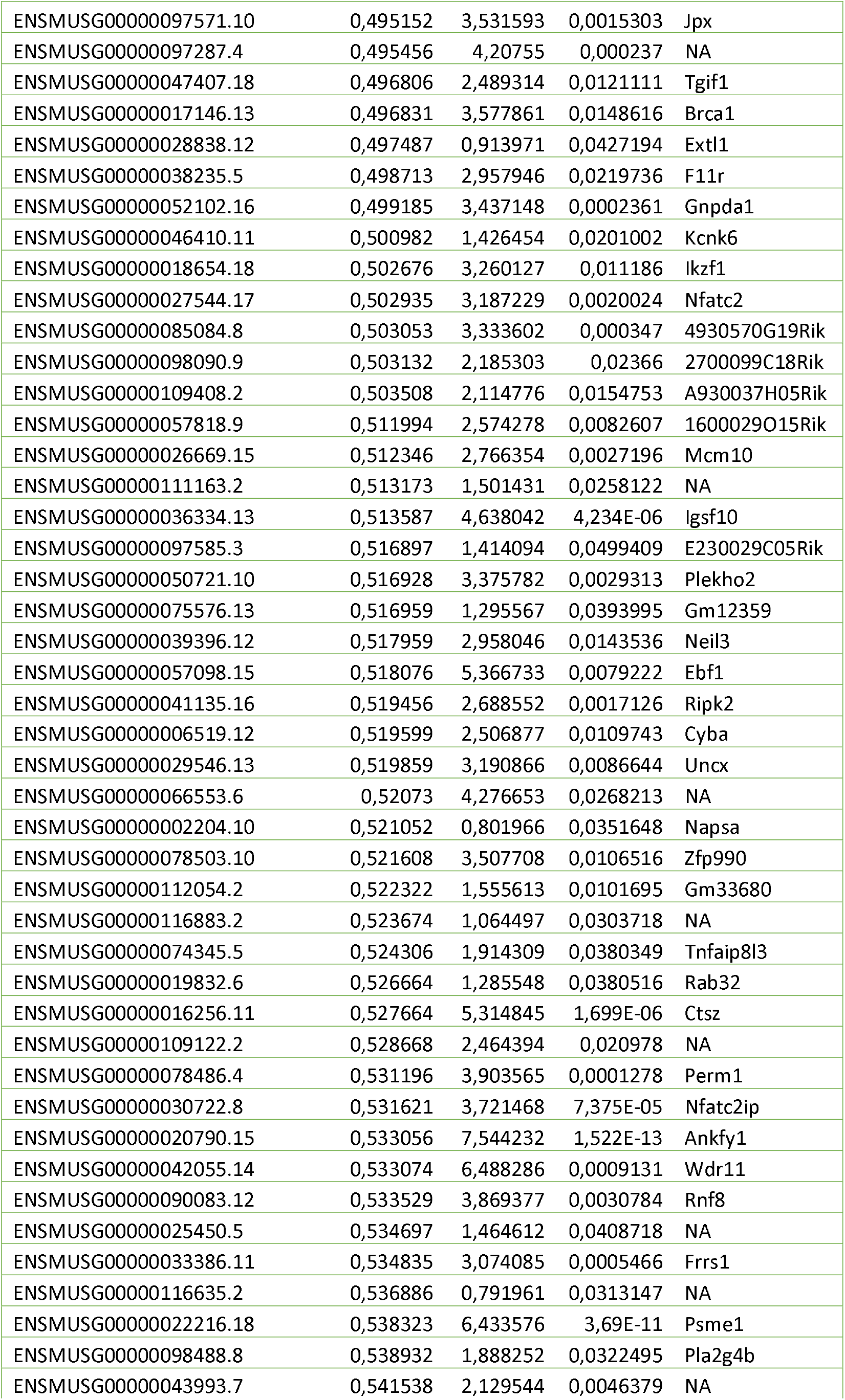
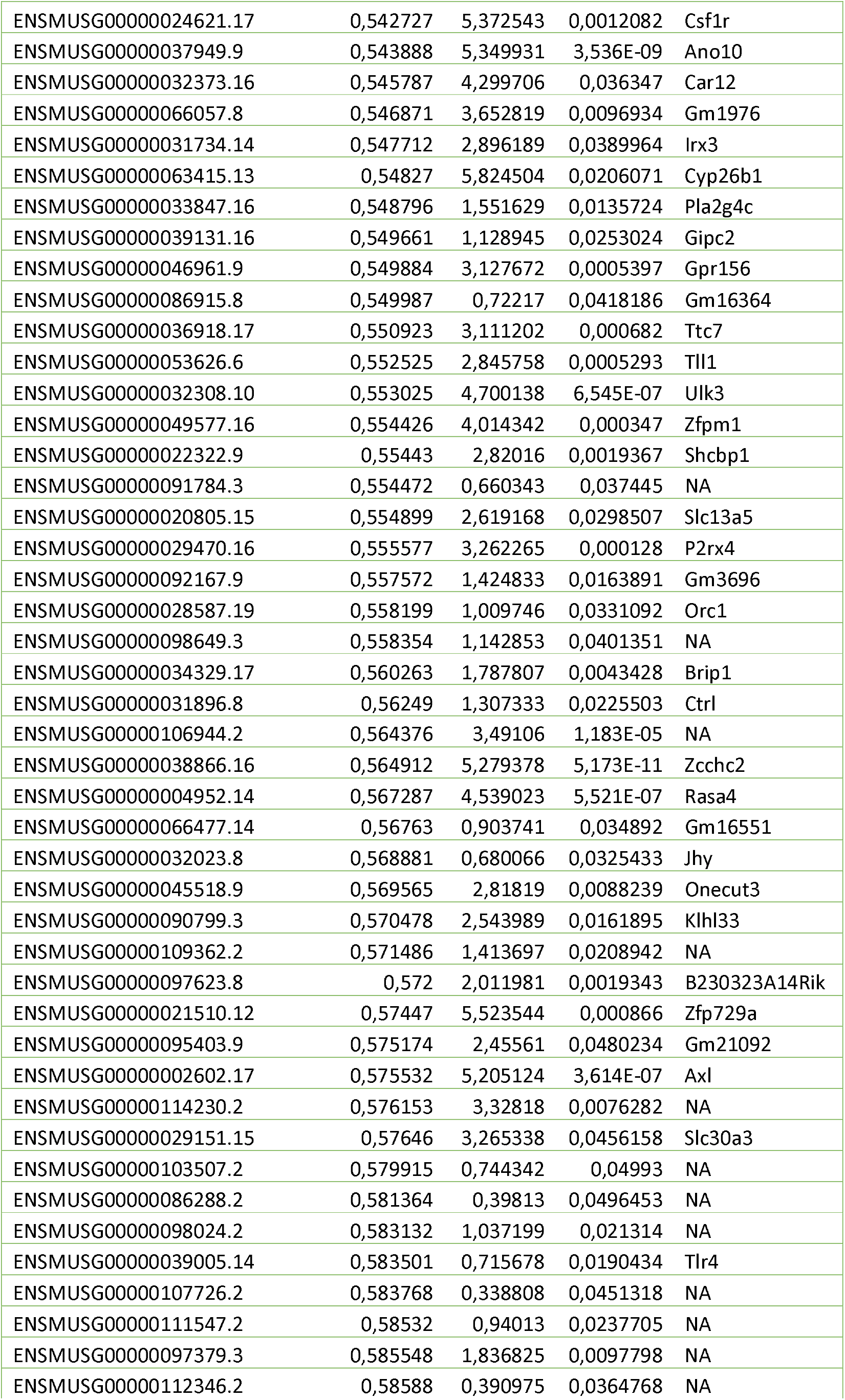
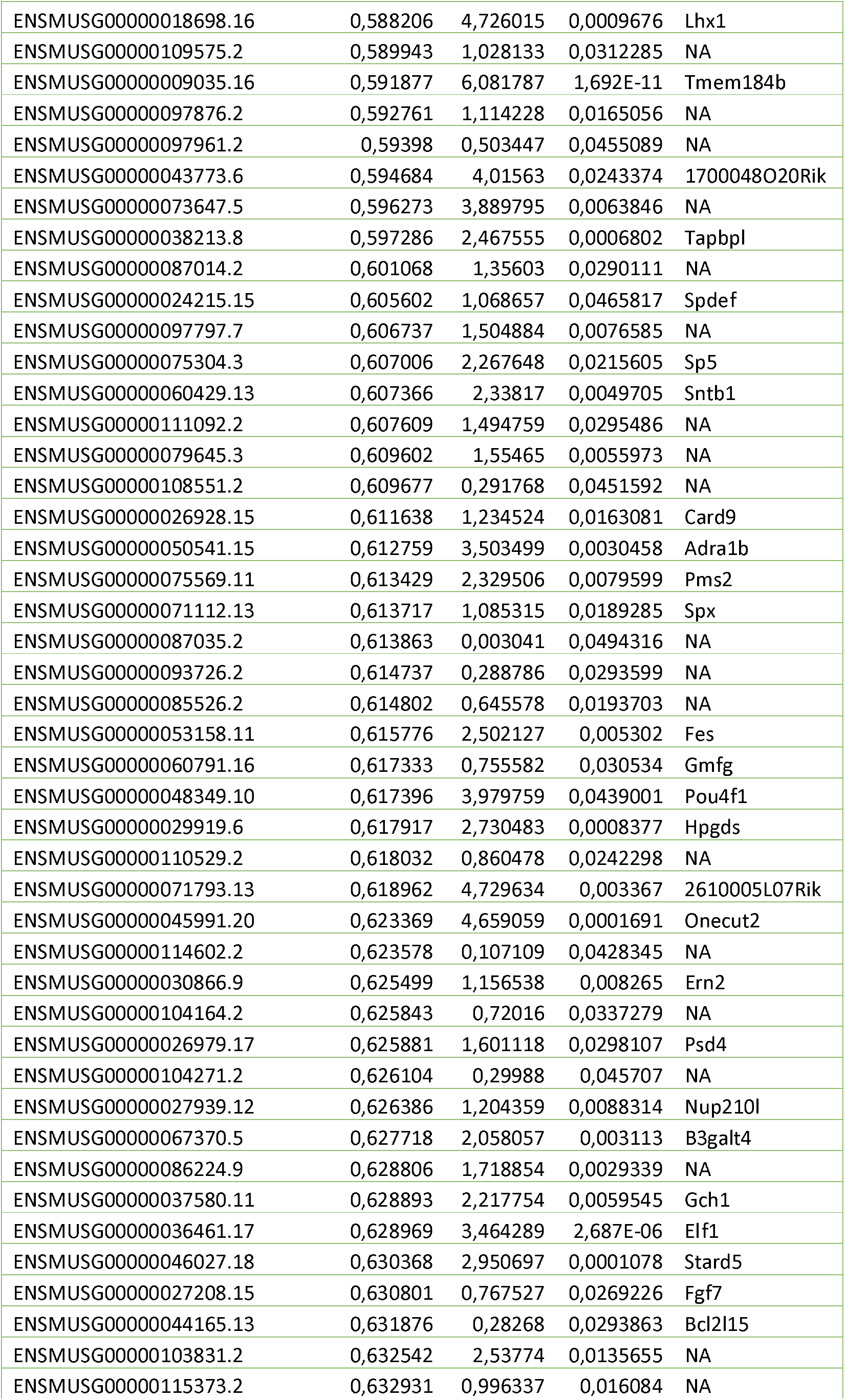
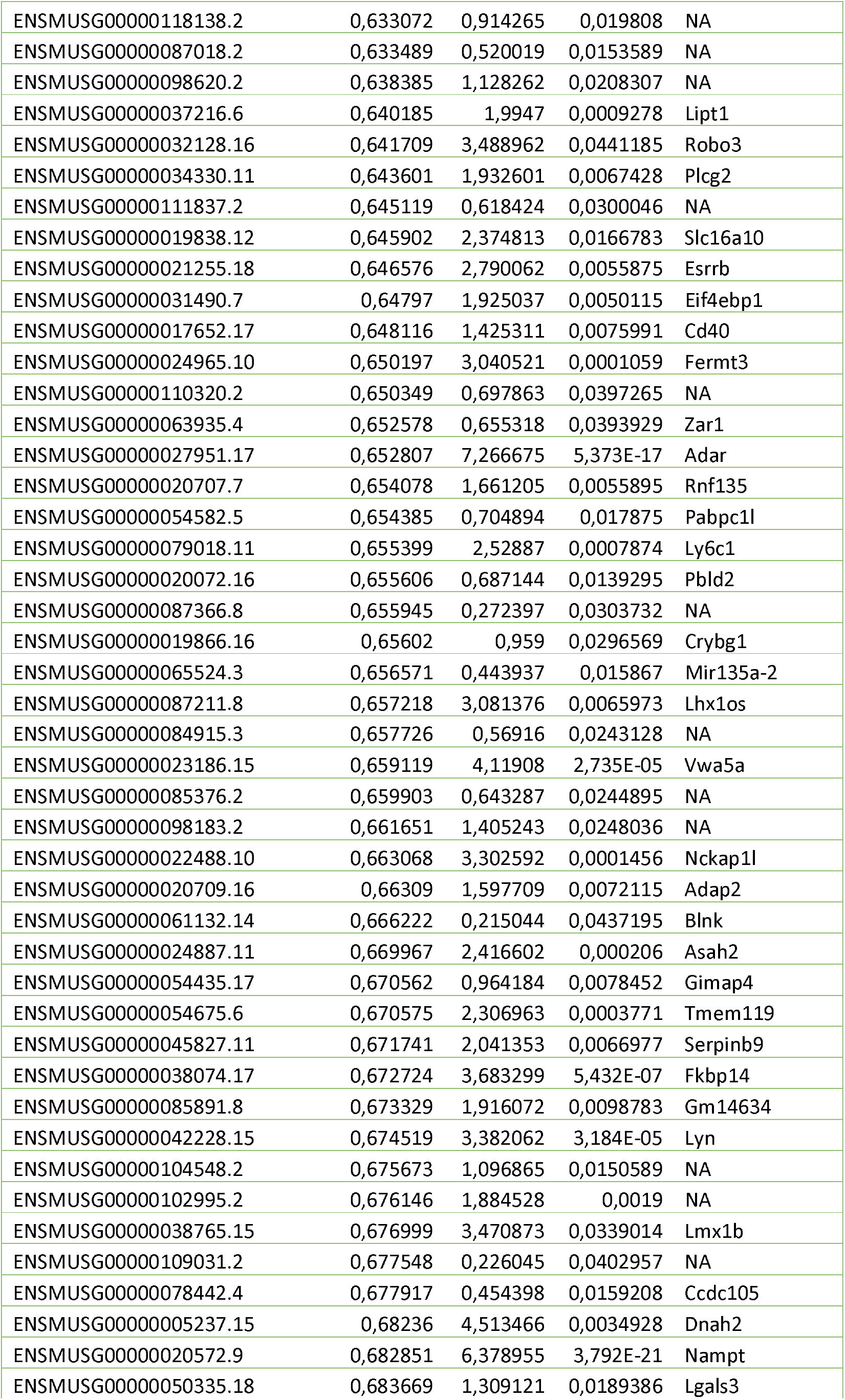
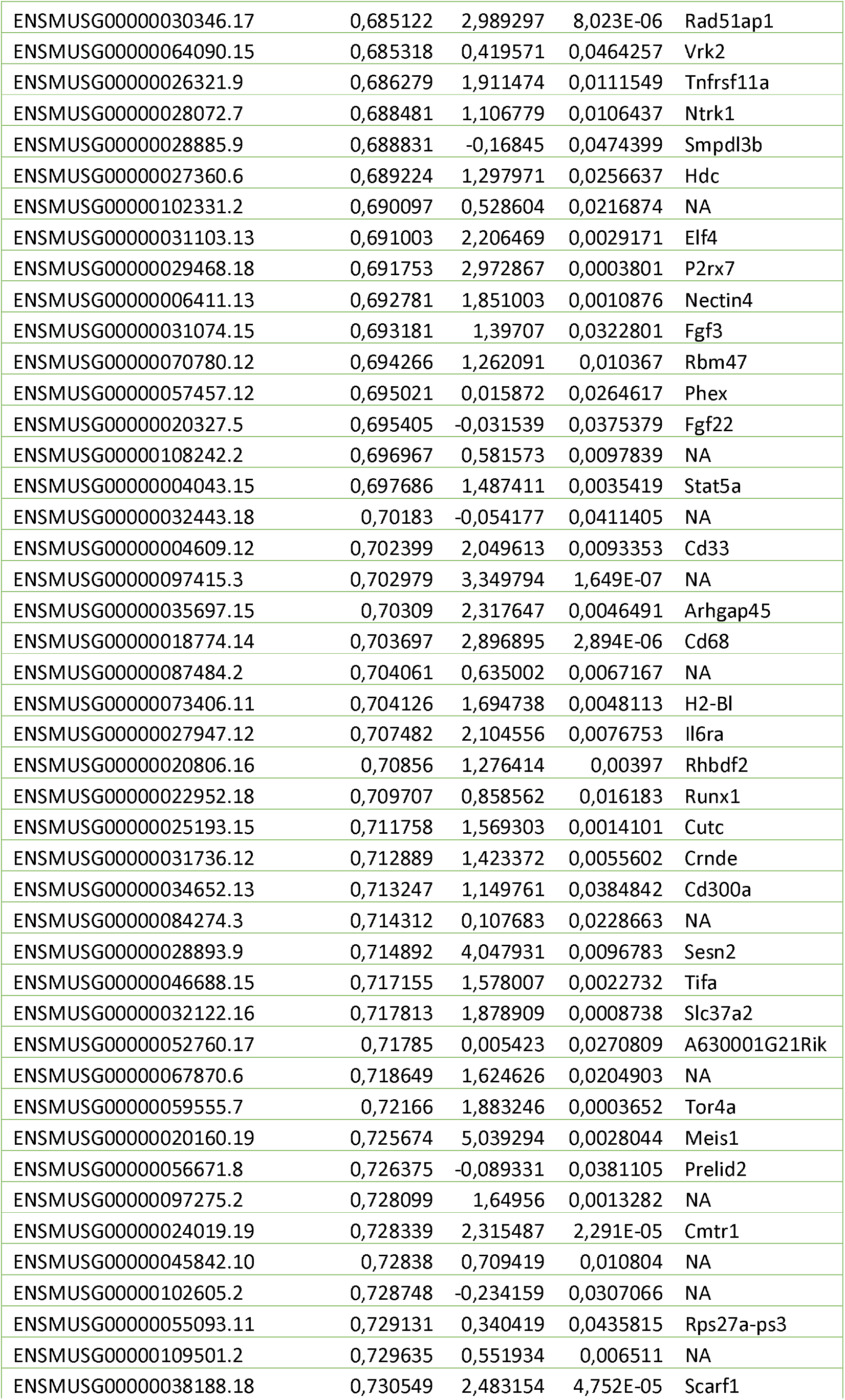
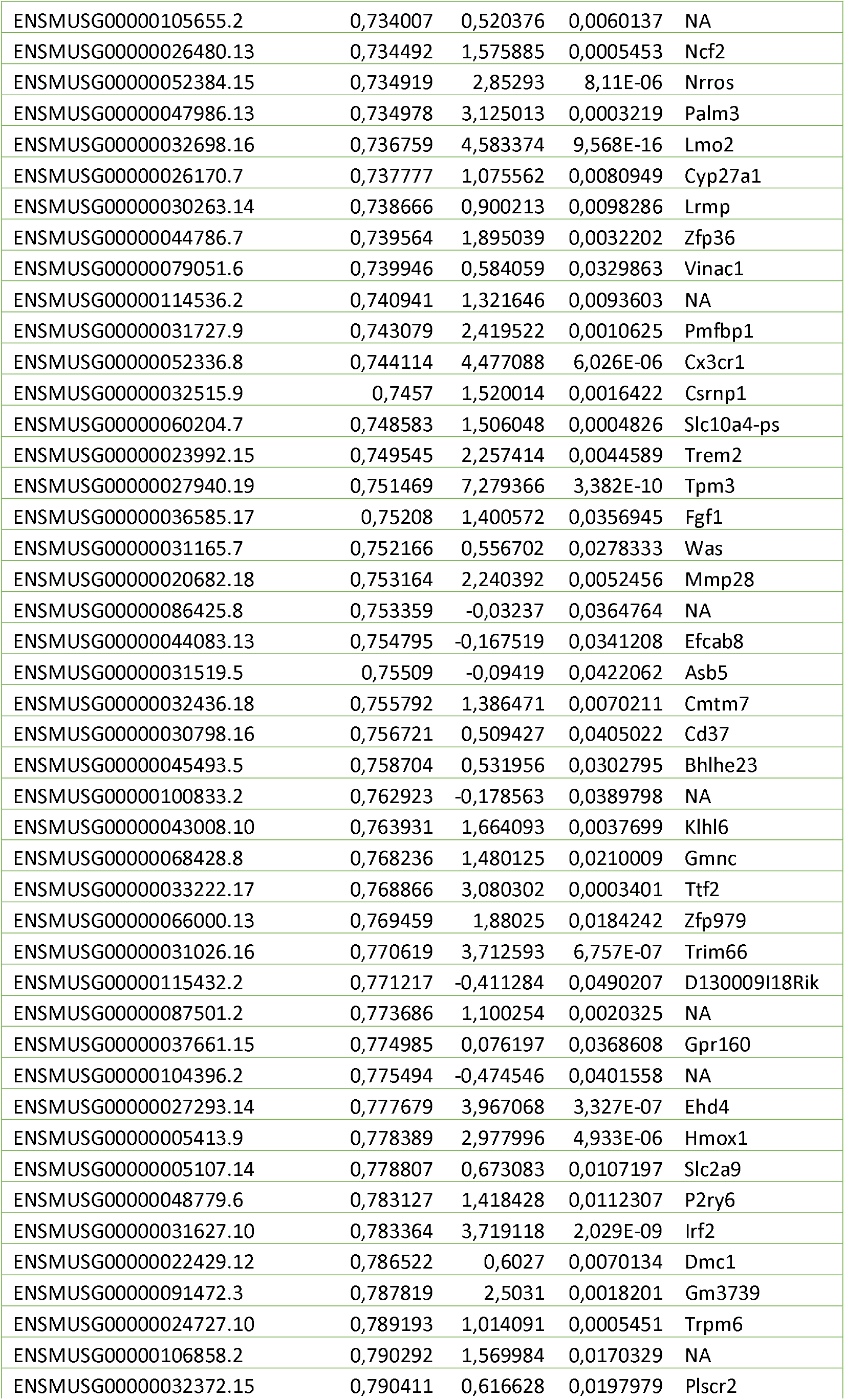
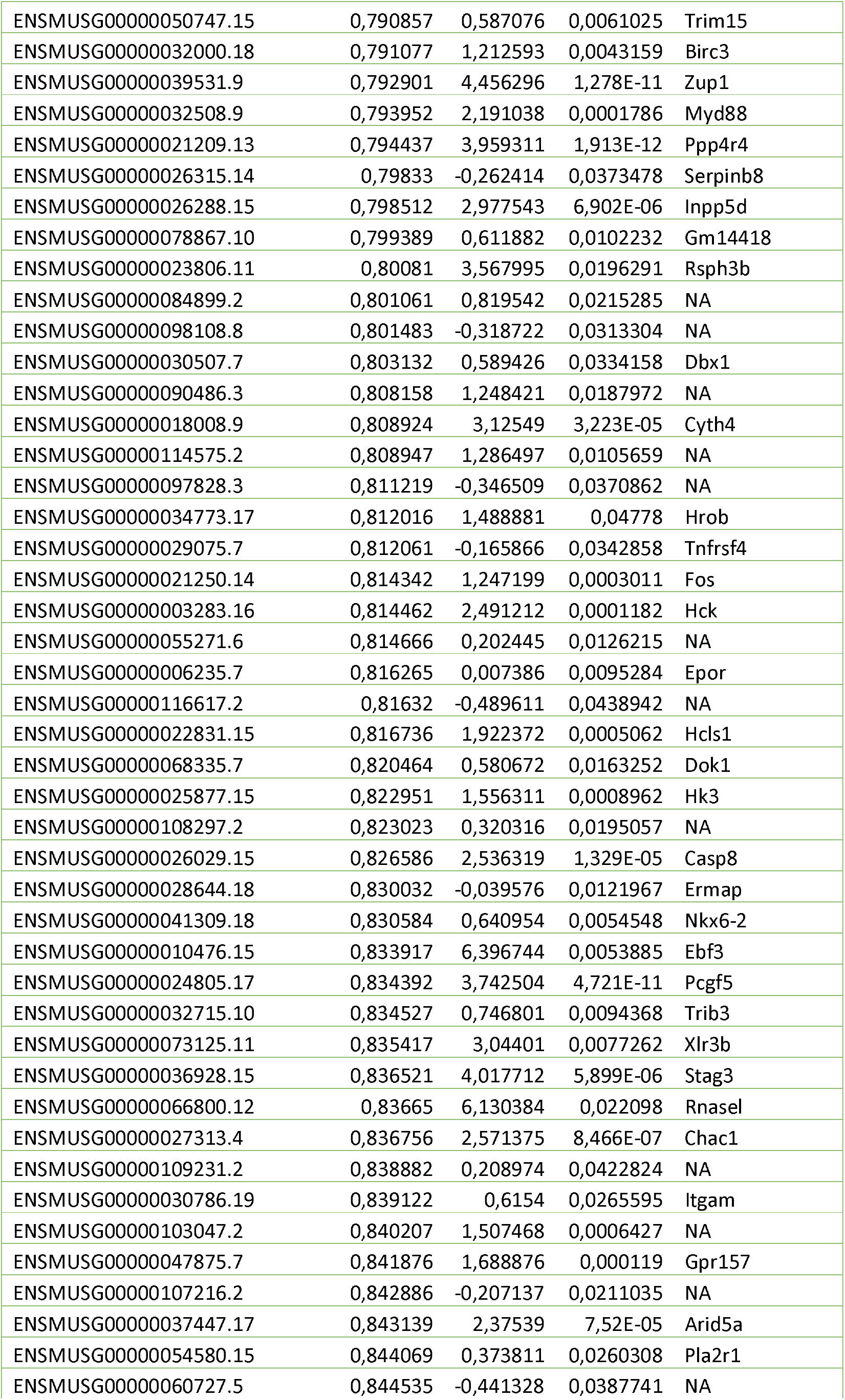
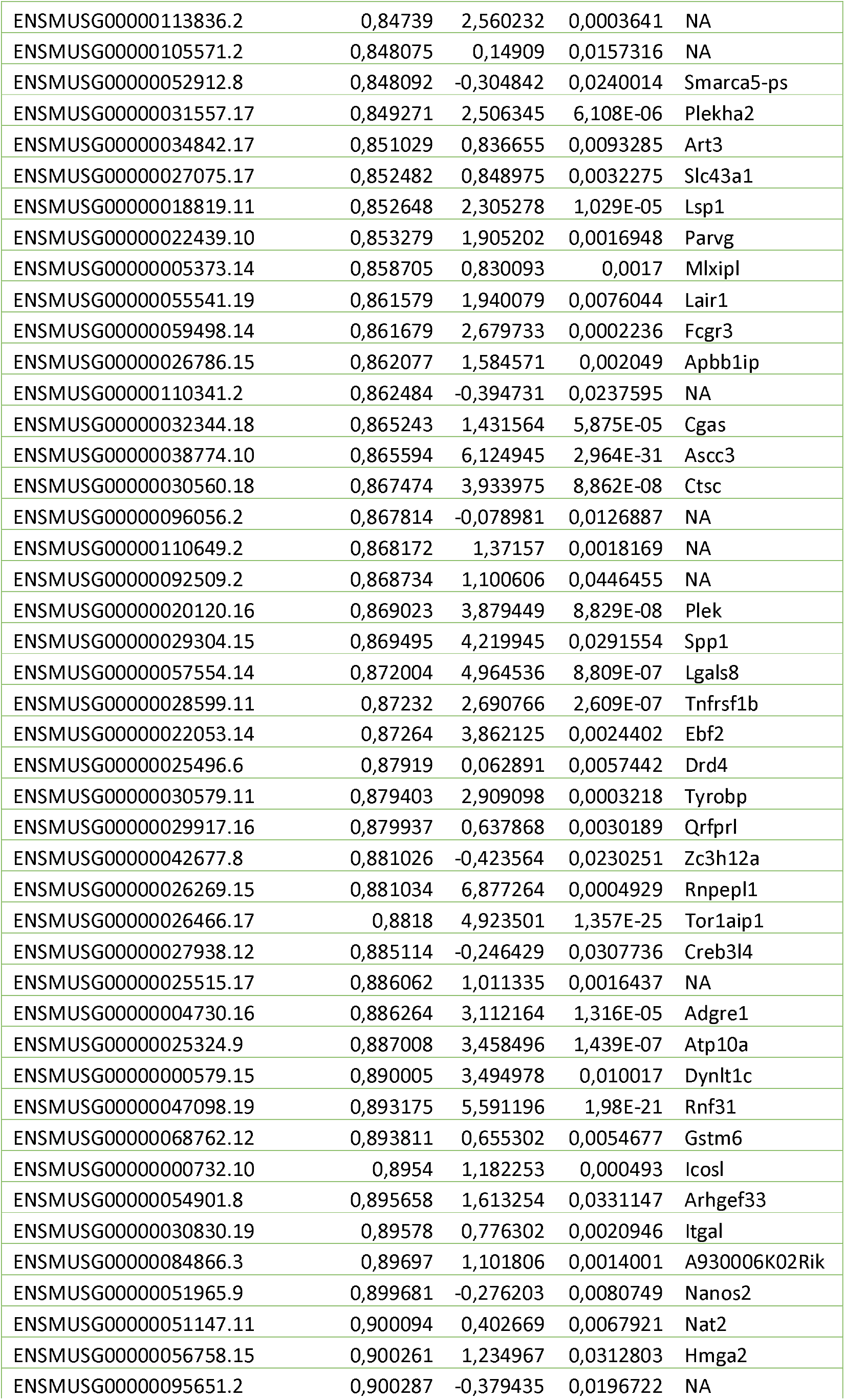
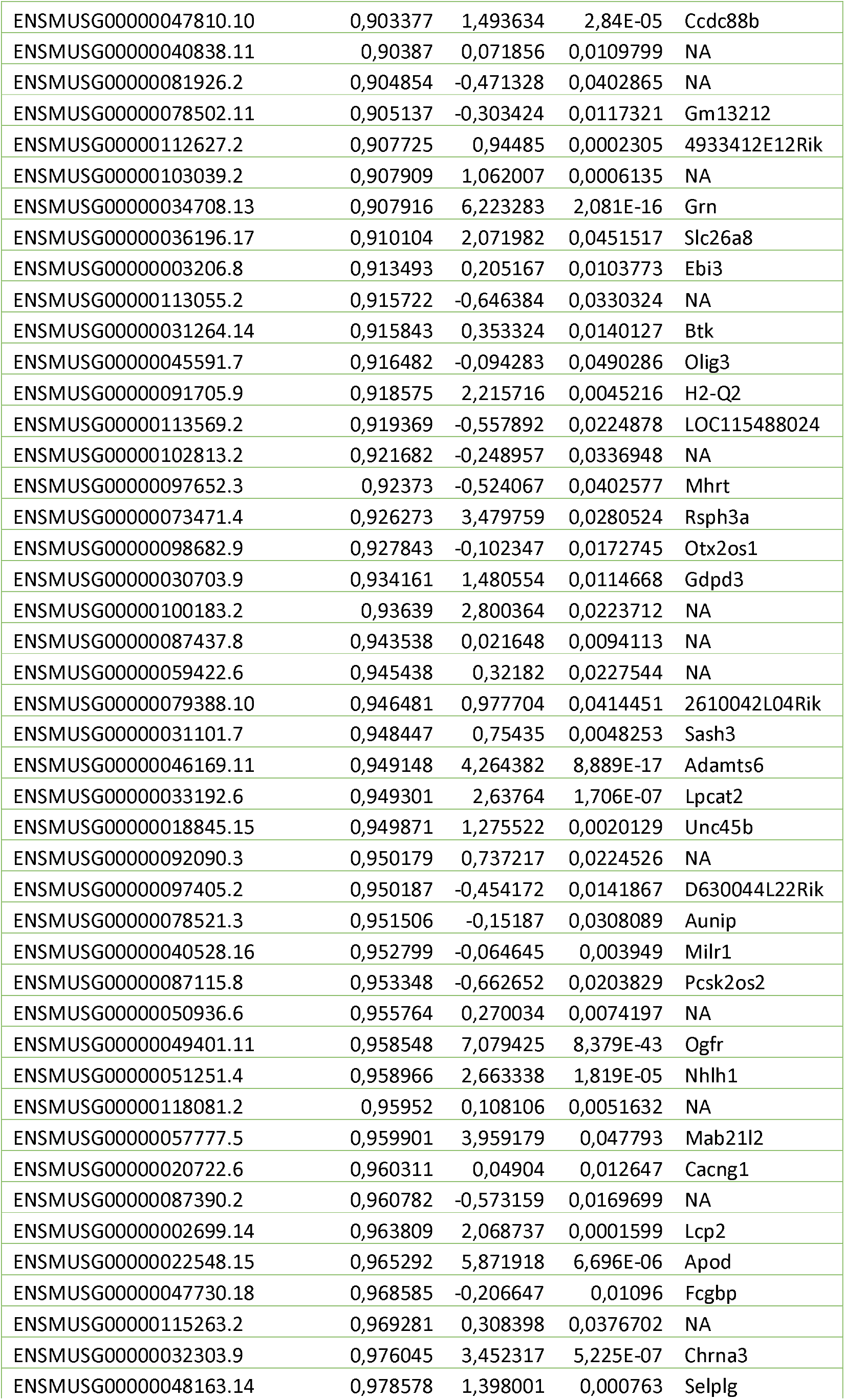
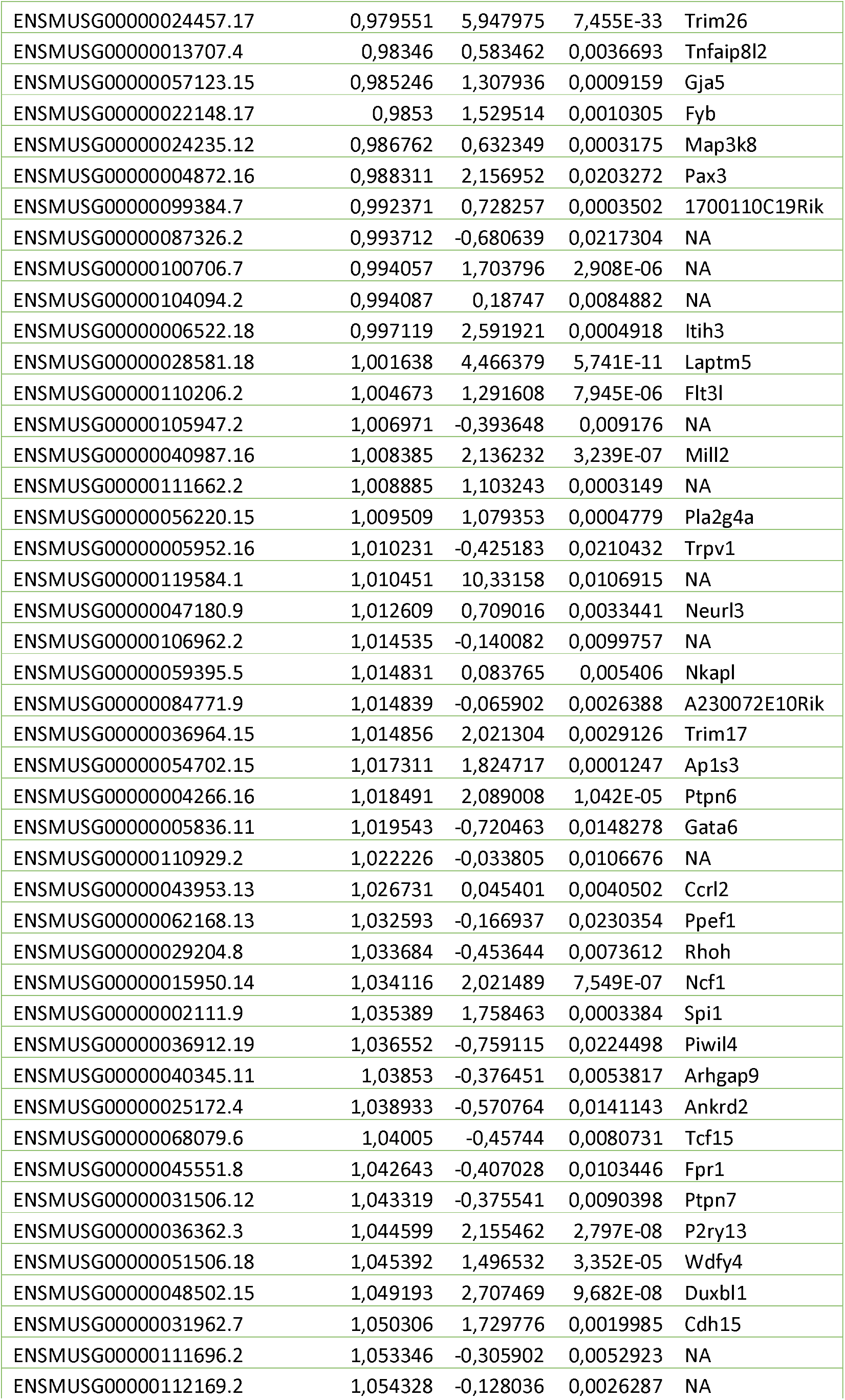
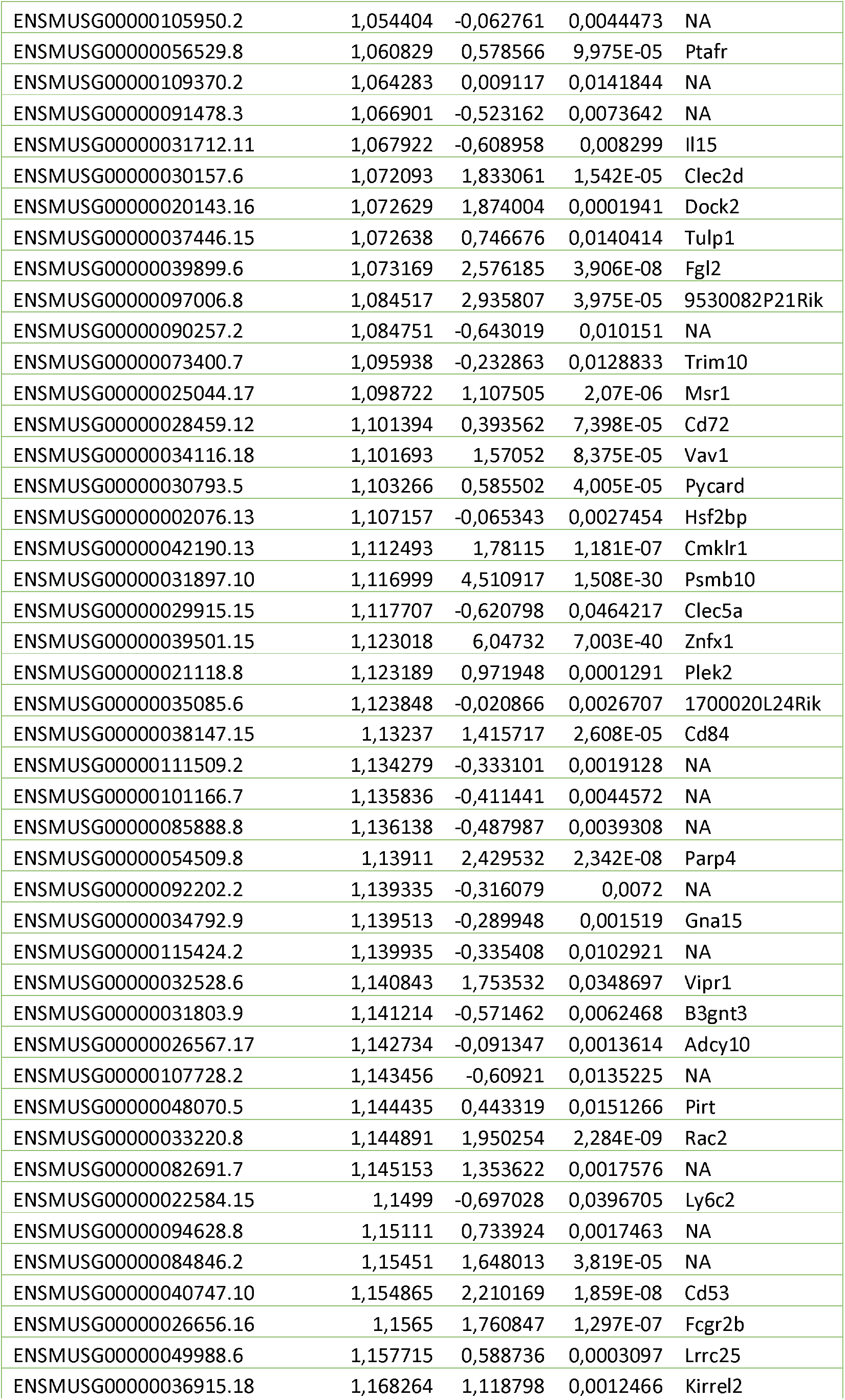
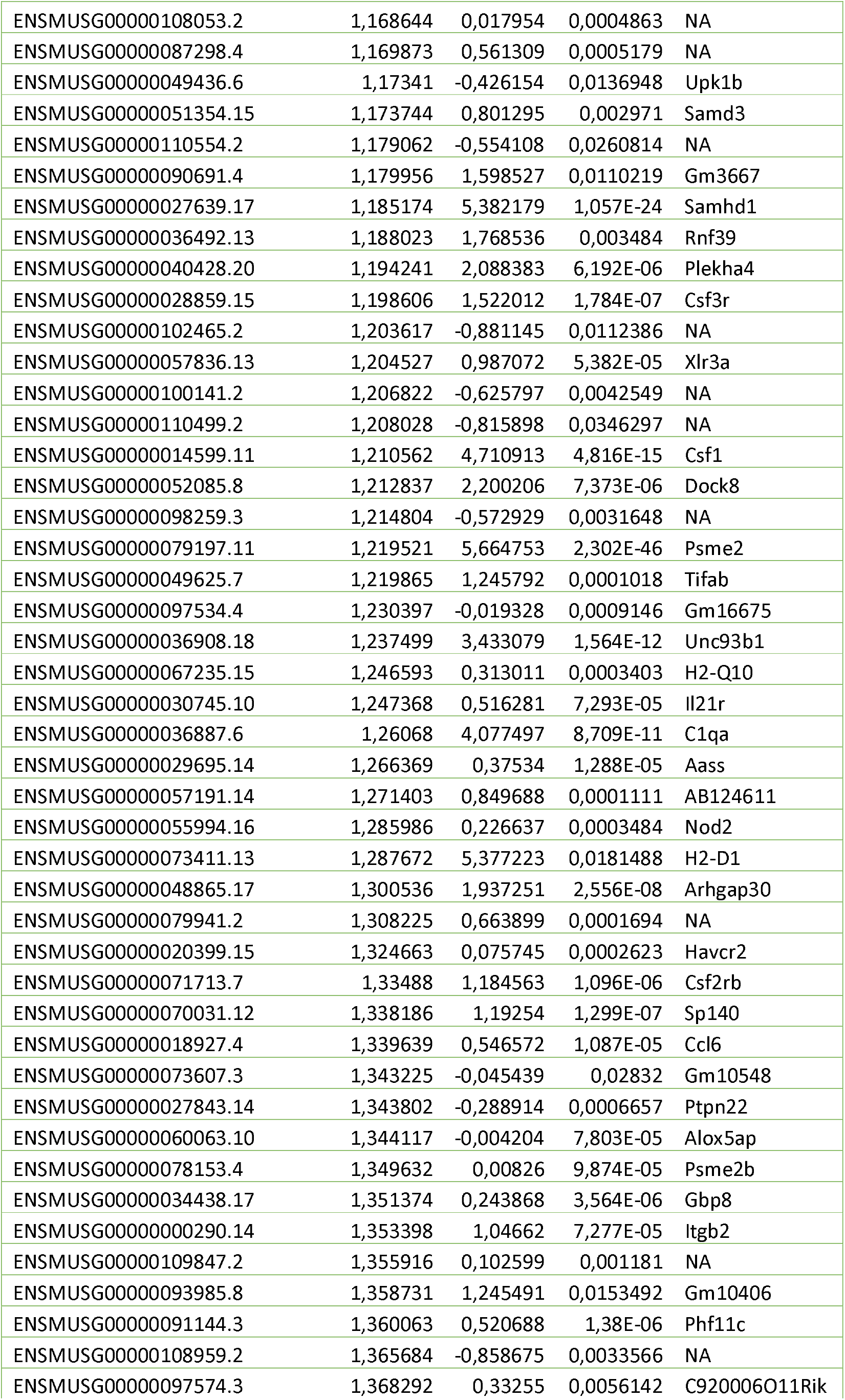
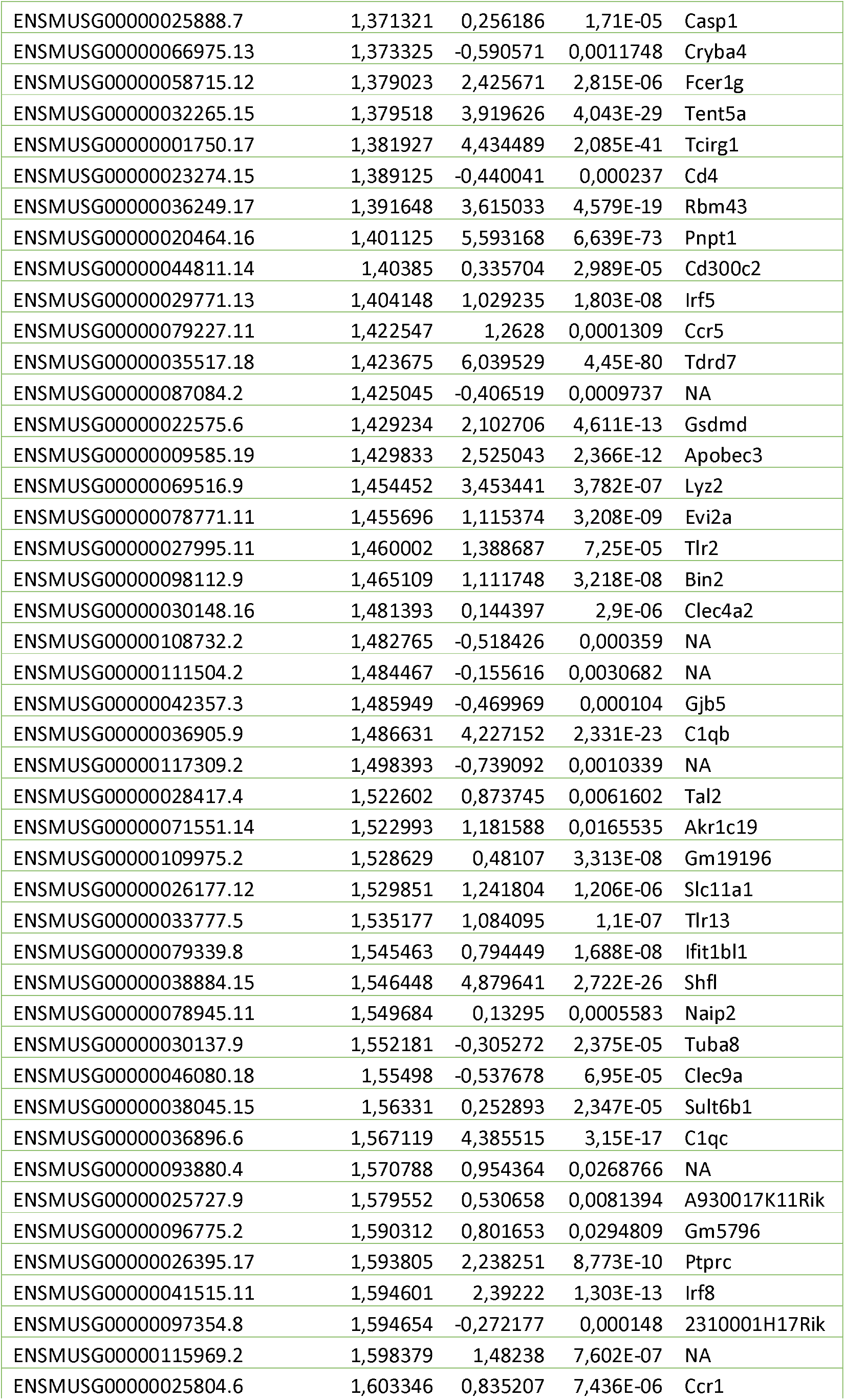
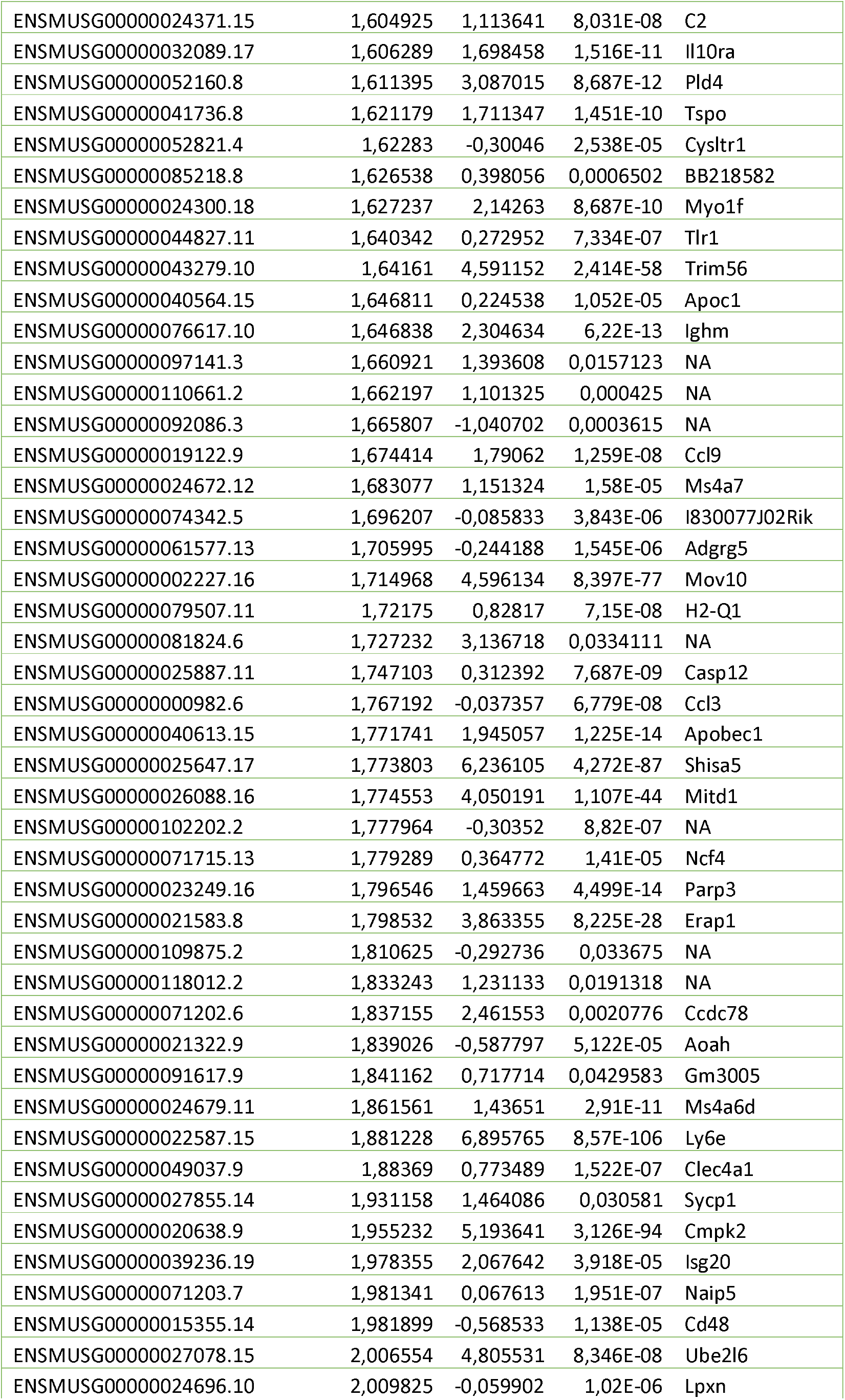
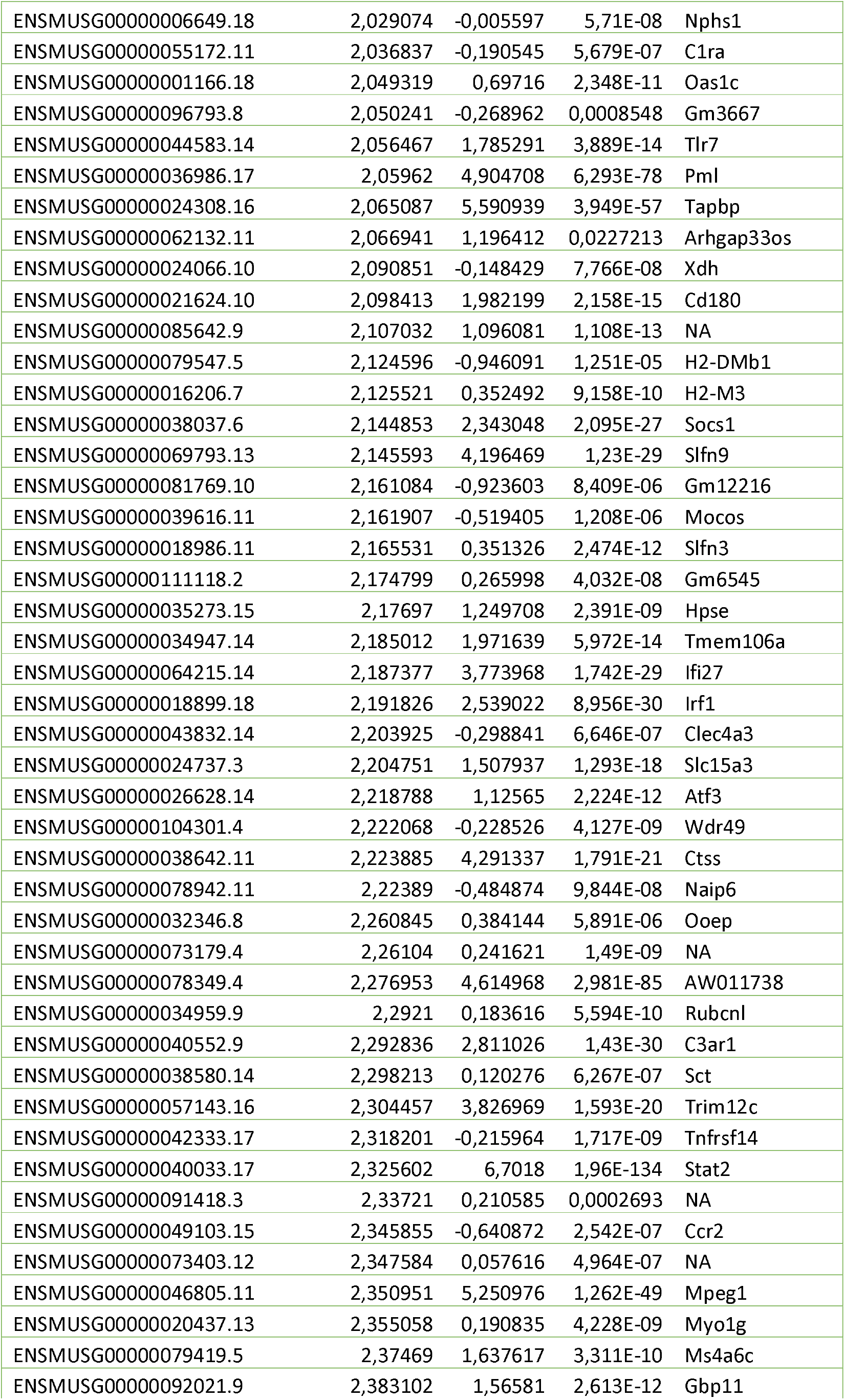
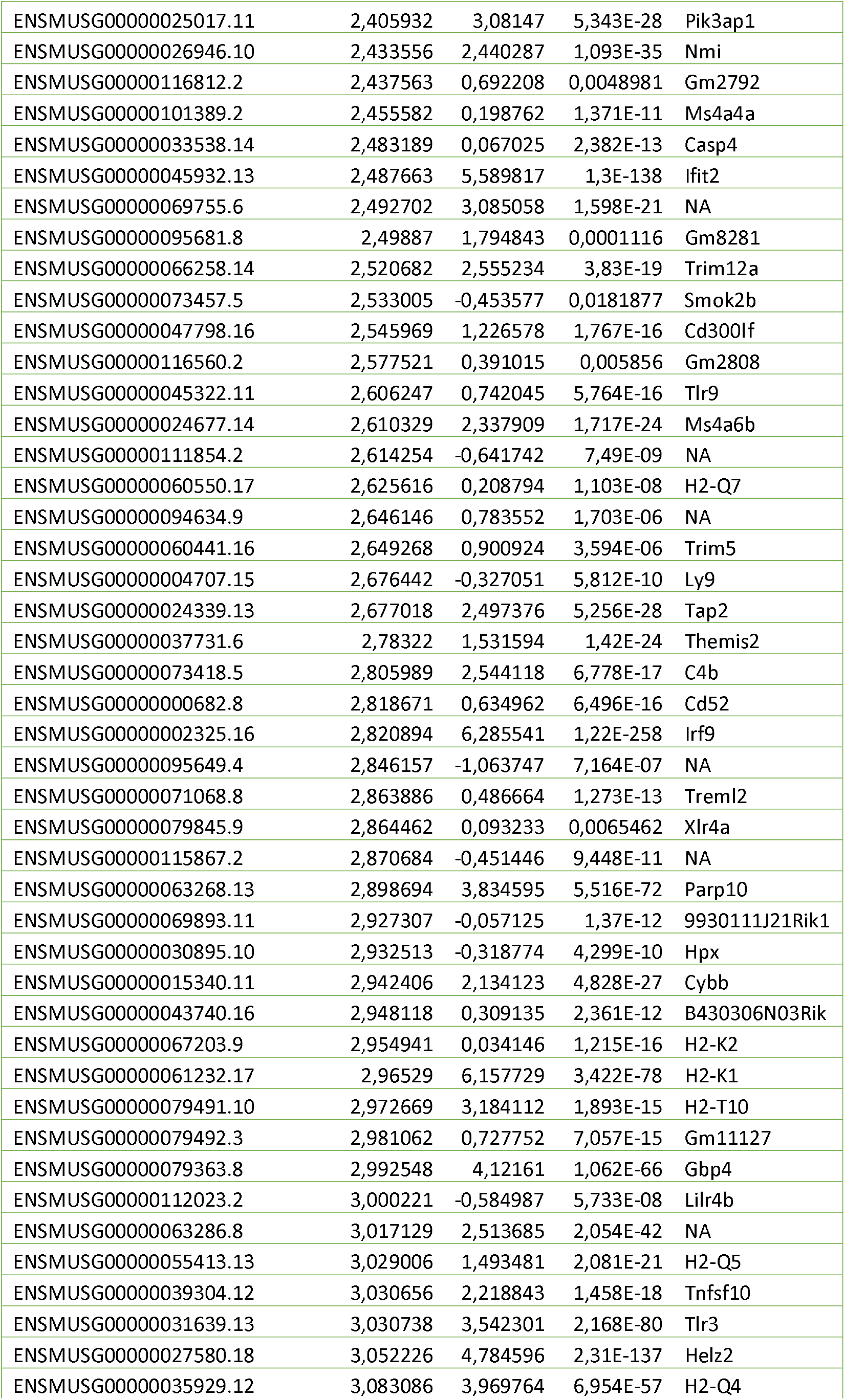
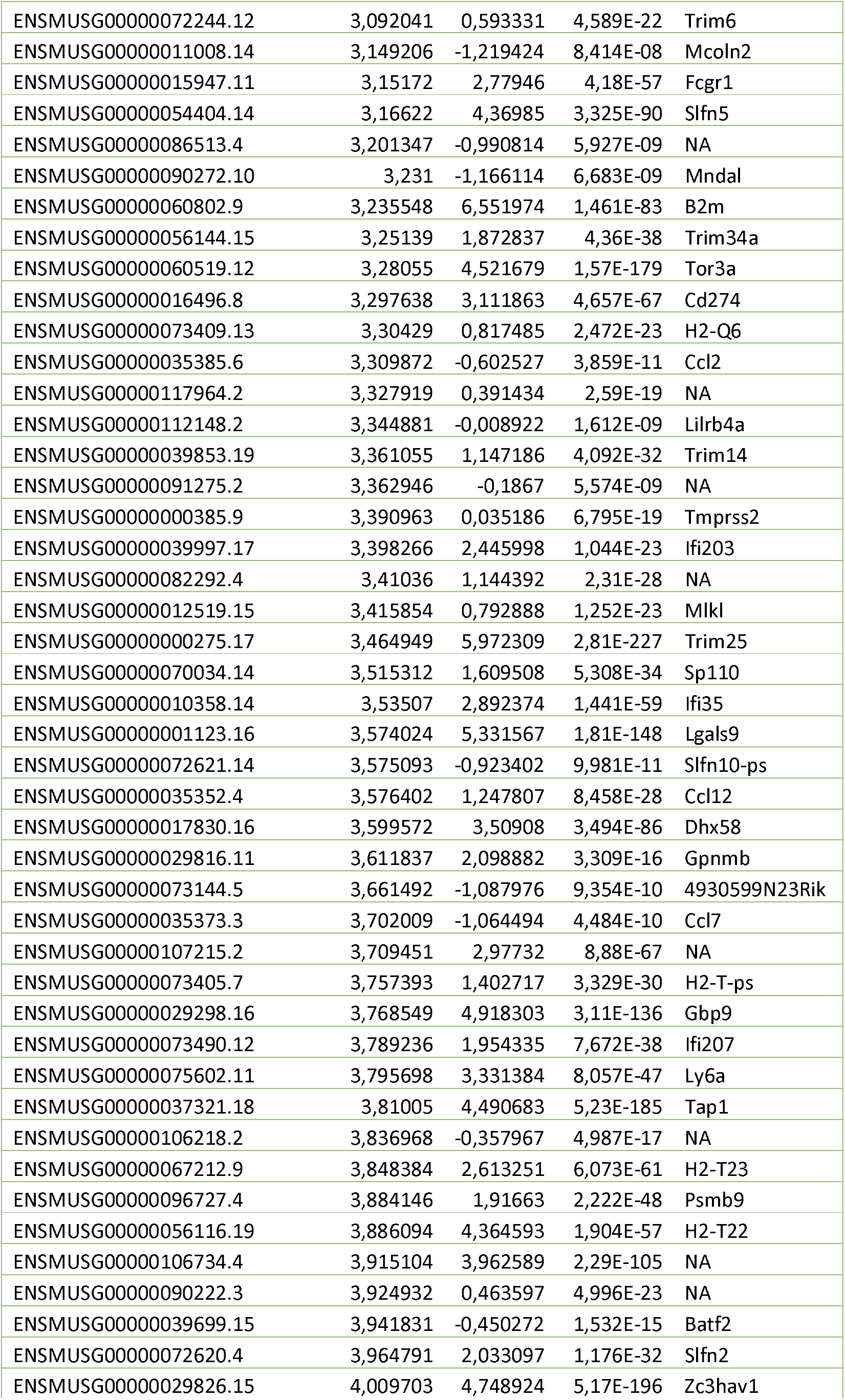
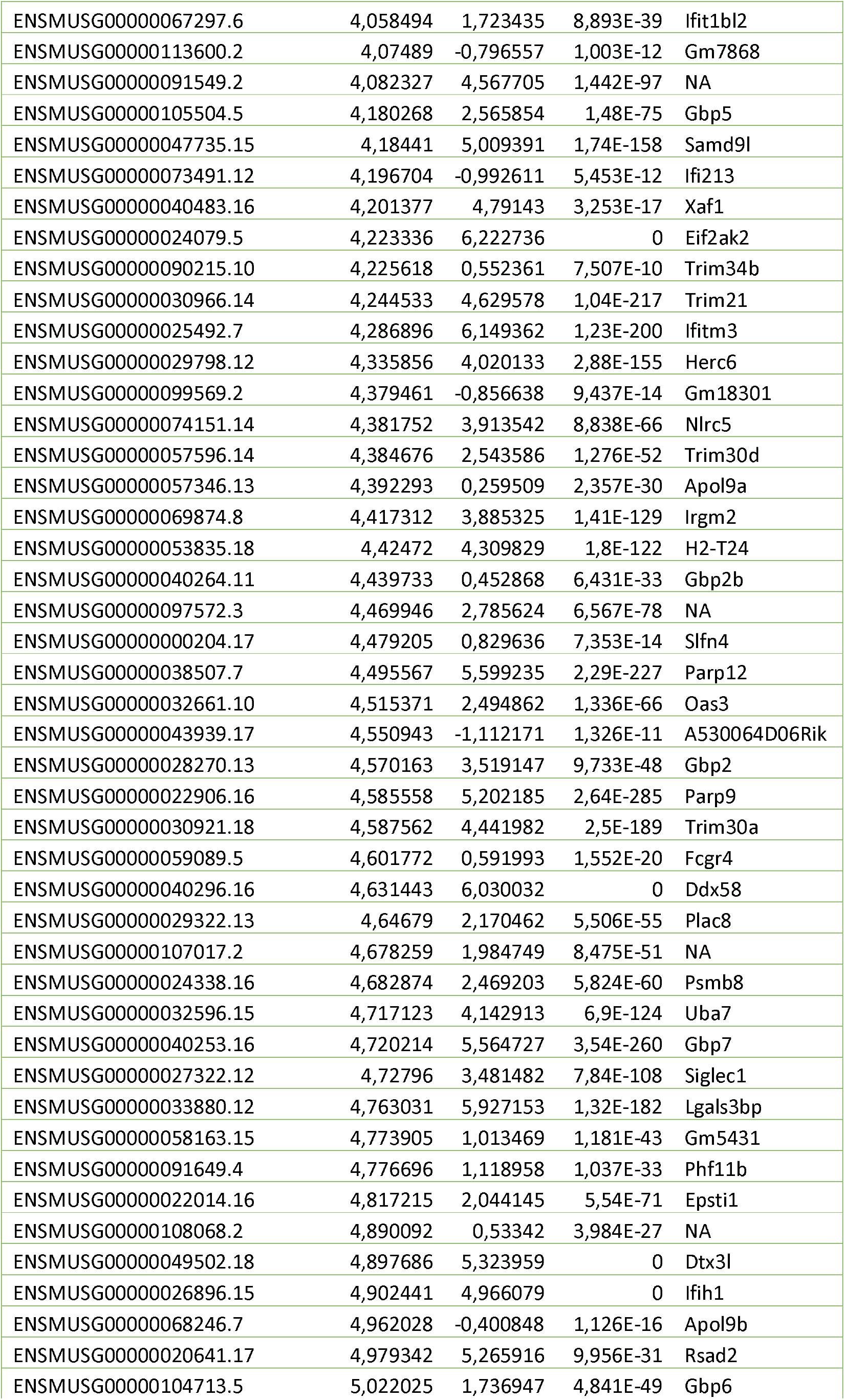
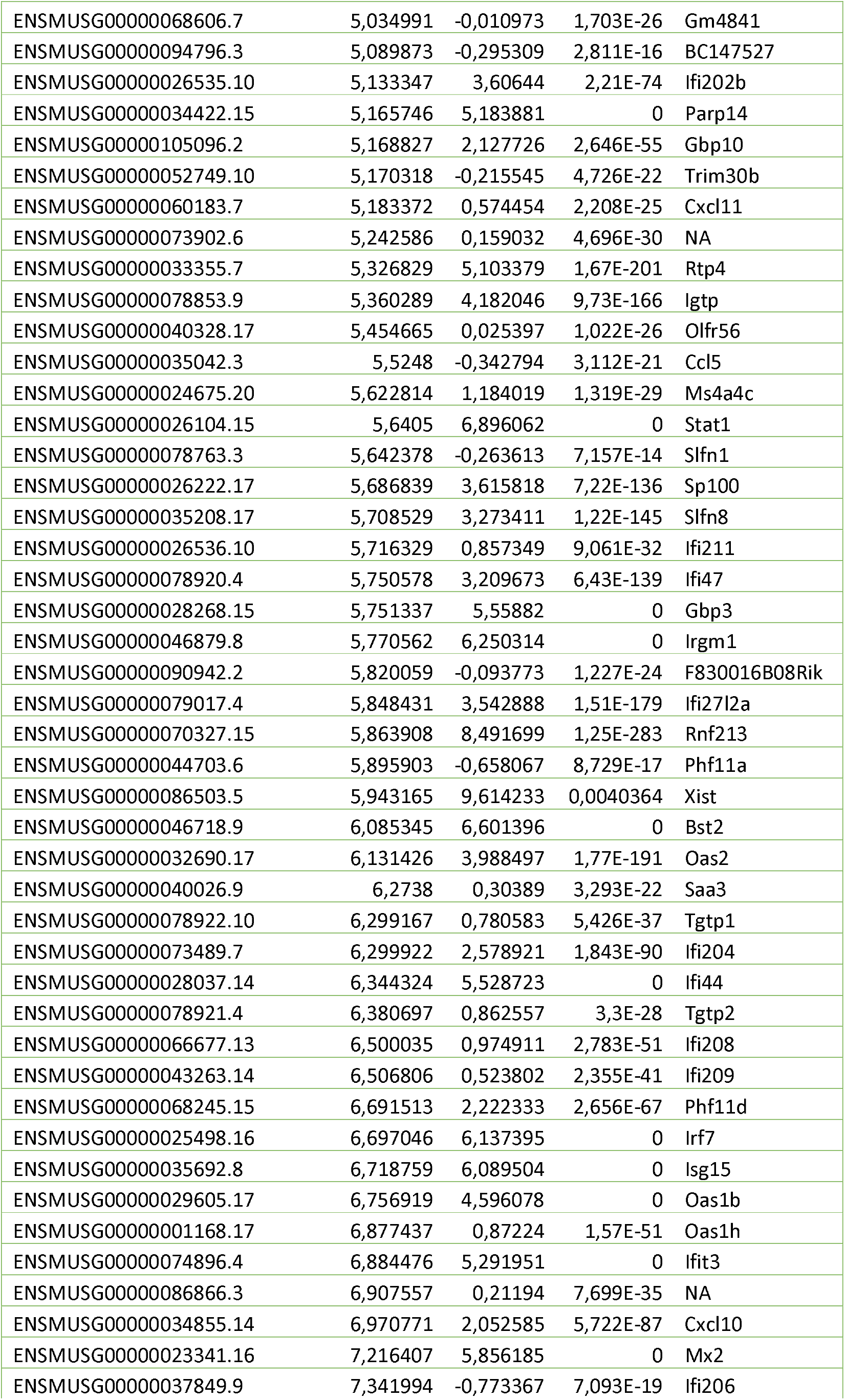
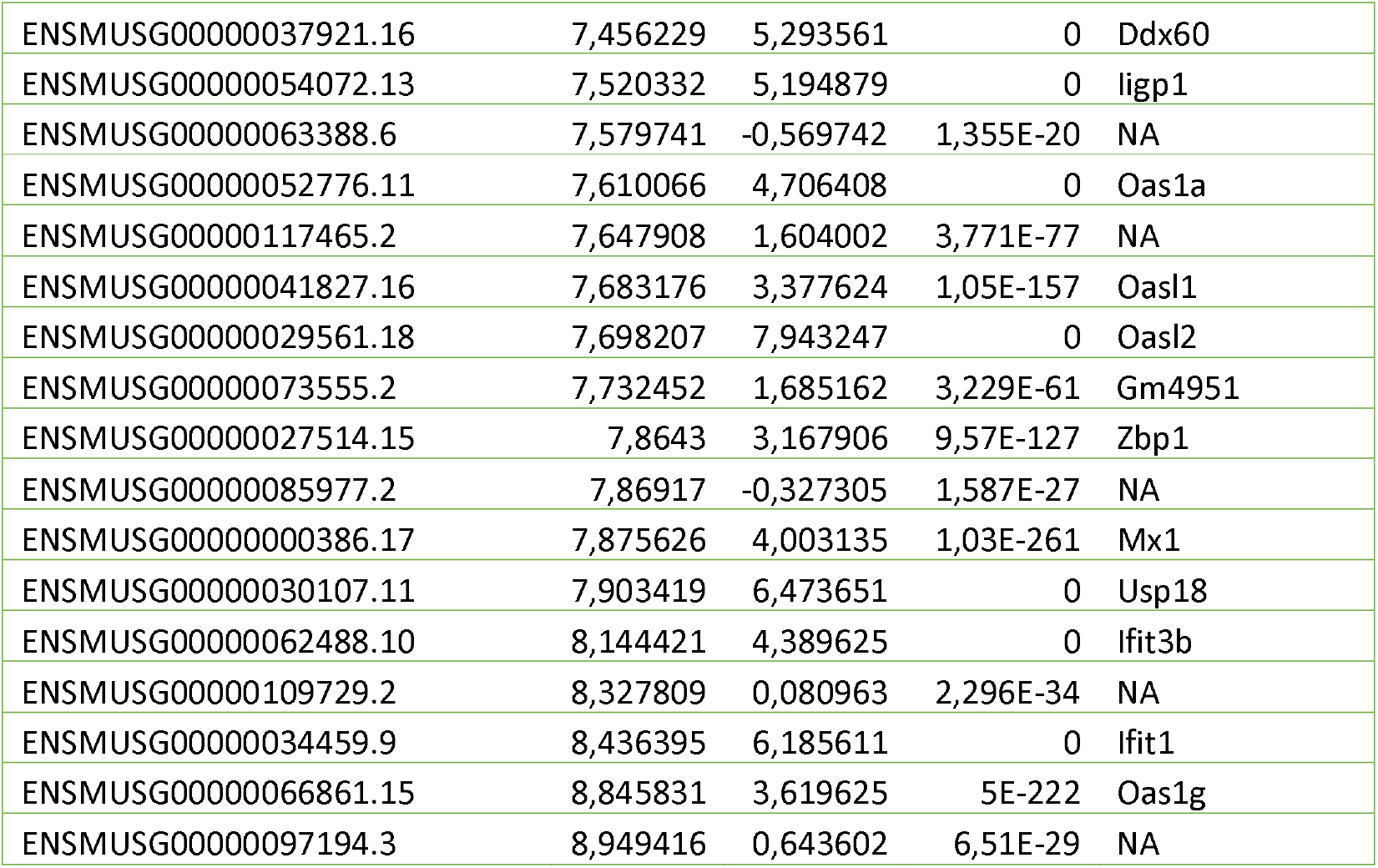
Differentially expressed genes between MOCK and ZIKV-infected brains. Embryos were infected *in utero* in E15 and harvested at E18. N= MOCK (5) ZIKV (3).

